# KDM4 Orchestrates Epigenomic Remodeling of Senescent Cells and Potentiates the Senescence-Associated Secretory Phenotype

**DOI:** 10.1101/2020.08.03.235465

**Authors:** Boyi Zhang, Qilai Long, Shanshan Wu, Shuling Song, Qixia Xu, Liu Han, Min Qian, Xiaohui Ren, Jing Jiang, Qiang Fu, Jianming Guo, Xiaoling Zhang, Xing Chang, Eric W-F Lam, Judith Campisi, James L. Kirkland, Yu Sun

**Affiliations:** CAS Key Laboratory of Tissue Microenvironment and Tumor, Shanghai Institute of Nutrition and Health, Shanghai Institutes for Biological Sciences, University of Chinese Academy of Sciences, Chinese Academy of Sciences, Shanghai 200031, China; Department of Urology, Zhongshan Hospital, Fudan University, Shanghai 200032, China; Institute of Health Sciences, Shanghai Jiao Tong University School of Medicine & Shanghai Institutes for Biological Sciences, Chinese Academy of Sciences, Shanghai 200031, China; Department of Pharmacology, Binzhou Medical University, Yantai, Shandong 264003, China; Department of Immunology, Binzhou Medical University, Yantai, Shandong 264003, China; Department of Orthopedic Surgery, Xinhua Hospital, Shanghai Jiao Tong University School of Medicine, Shanghai 200092, China; Key Laboratory of Structural Biology of Zhejiang Province, School of Life Sciences, Westlake University, Hangzhou, Zhejiang 310024, China; Department of Surgery and Cancer, Imperial College London, London, W12 0NN, UK; Buck Institute for Research on Aging, Novato, CA 94945, USA; Life Sciences Division, Lawrence Berkeley National Laboratory, Berkeley, CA 94720, USA; Robert and Arlene Kogod Center on Aging, Mayo Clinic, Rochester, MN 55905, USA; Department of Medicine and VAPSHCS, University of Washington, Seattle, WA 98195, USA

## Abstract

Cellular senescence restrains the expansion of neoplastic cells through several layers of regulation, including epigenetic decoration of chromatin structure and functional modulation of bioactive components. Here we report that expression of the histone H3-specific demethylase KDM4 is upregulated in human stromal cells upon cellular senescence. In clinical oncology, upregulated KDM4 and diminished H3K9/H3K36 methylation are correlated with adverse survival of cancer patients post-chemotherapy. Global chromatin accessibility mapping *via* ATAC-seq and expression profiling through RNA-seq reveal extensive reorganization of chromosomes and spatiotemporal reprogramming of the transcriptomic landscape, events responsible for development of the senescence-associated secretory phenotype (SASP). Selectively targeting KDM4 dampens the SASP of senescent stromal cells and enhances the apoptotic index of cancer cells in the treatment-damaged tumor microenvironment (TME), together prolonging overall survival of experimental animals. Our study supports the dynamic change of H3K9/H3K36 methylation marks during cellular senescence, identifies an unusually permissive chromatin state, unmasks KDM4 as a key modulator of the SASP, and presents a novel therapeutic avenue to manipulate cellular senescence and curtail age-related pathologies.

## Introduction

Senescence is a self-control mechanism that limits cell hyper-proliferation and prevents neoplastic progression by implementing a stable, durable, albeit generally irreversible growth arrest. Senescent cells display profound alterations in nuclear and chromatin structure, expression patterns and metabolic activities ^1^. Senescent cells also actively secrete a large array of proteins, many of which are pro-inflammatory factors *per se*, a property collectively termed the senescence-associated secretory phenotype (SASP)^2–4^.

Several lines of evidence suggest that the SASP is essential for tissue repair, wound healing, embryonic development, and immune surveillance to clear senescent cells ^5^. However, as a matter of evolutionary pleiotropy, this phenotype is mostly detrimental in numerous pathological settings ^6^. For example, in a local tumor microenvironment (TME), the secretome of lingering senescent cells can markedly promote malignancy of neighboring cancer cells, particularly chemoresistance, and cause chronic inflammation associated with many age-related disorders ^7–9^. Despite limited therapeutic efficacy-promoting outcomes that depend on a given context ^10^, the net effect of the SASP in advanced cancers far outweighs its beneficial contributions, ultimately accelerating disease progression ^5, 7^. Specifically, the harmful inflammation imposed by the SASP suggests that eliminating senescent cells or suppressing the SASP can be advantageous in multiple types of pathologies and not just cancer ^8, 9, 11^.

The structure of chromatin is precisely regulated by epigenetic modifications such as DNA methylation and histone posttranslational modification (PTM), which functionally govern gene expression by changing chromatin organization and genome accessibility. The epigenetic landscape is prone to perpetual fluctuations throughout the lifespan and is profoundly altered in aging organisms ^12^. Epigenetic marks potently control heterochromatin segments and the 3-dimensional folding of chromosomes, as well as genetic loci that regulate cell type commitment, proliferation, and cellular senescence, *e.g.*, the *INK4* box ^13^. However, perturbations of histone methylation-associated marks, such as loss of H3K9me3 and H3K27me3, are observed in normal human aging and premature aging diseases, such as Hutchinson-Gilford progeria syndrome (HGPS) or Werner syndrome (WS), suggesting that global decline of heterochromatin may be a common feature of aging ^14^.

Lysine methylation is one of the major histone PTM forms that regulate chromatin structure. KDM4 (JMJD2) refers to a subfamily of demethylases that have a Jumonji domain and target histone H3 on lysines 9 and 36 sites (H3K9 and H3K36), consisting of three 130 KD proteins (KDM4A-C) and KDM4D, which is half the size and lacks double PHD and tandem Tudor domains as epigenome readers ^15^. The role of the KDM4 subfamily is reported in multiple human cancer types, and small molecule inhibitors of KDM4 are being actively pursued to complement the existing arsenal of epigenetic drugs targeting DNA methyltransferases and histone deacetylases ^16^. It was reported that KDM4A can suppress Ras^V12^-induced senescence and collaborate with this oncogene to promote cellular transformation by negatively regulating the p53 pathway ^17^. Further, KDM4A is critical for completion of DNA double strand break (DSB) repair and modulates the relative utilization of homologous recombination (HR) and non-homologous end-joining (NHEJ) repair pathways, but exclusively for heterochromatic DSBs ^18^. Knockdown of KDM4B induces DNA damage *via* ATM and Rad3-related pathway activation, promoting cell cycle arrest, apoptosis, and senescence in both normoxic and hypoxic conditions ^19^. Despite accumulated data addressing the involvement of KDM4 in DNA damage response (DDR) and cancer progression, its implications in cellular senescence and SASP development remain largely underexplored. Understanding the significance of KDM4, a typical epigenetic factor that holds the potential to reprogram the chromatin landscape in senescent cells, is highly warranted in aging biology and geriatric medicine.

## Results

### Histone H3 lysine sites in senescent cells are epigenetically modified

To establish an unbiased epigenetic profile of senescent cells at the protein level, we chose to use stable isotope labeling with amino acids (SILACs), a mass spectrometry (MS)-based technique involving a non-radioactive isotope, to analyze senescent cells (SEN) induced by bleomycin (BLEO) and their proliferating counterparts (CTRL) (Fig. 1A). We examined independent biological replicates for each group and confirmed DNA damage-induced cellular senescence in a primary human prostate stromal cell line (PSC27), which consists predominantly of fibroblasts ^20^. Overall, we detected 732 intracellular proteins with 95% confidence, of which 447 entities are quantifiable (Fig. 1B-1C) (Supplementary Tables 1-2).

**Fig 1.**
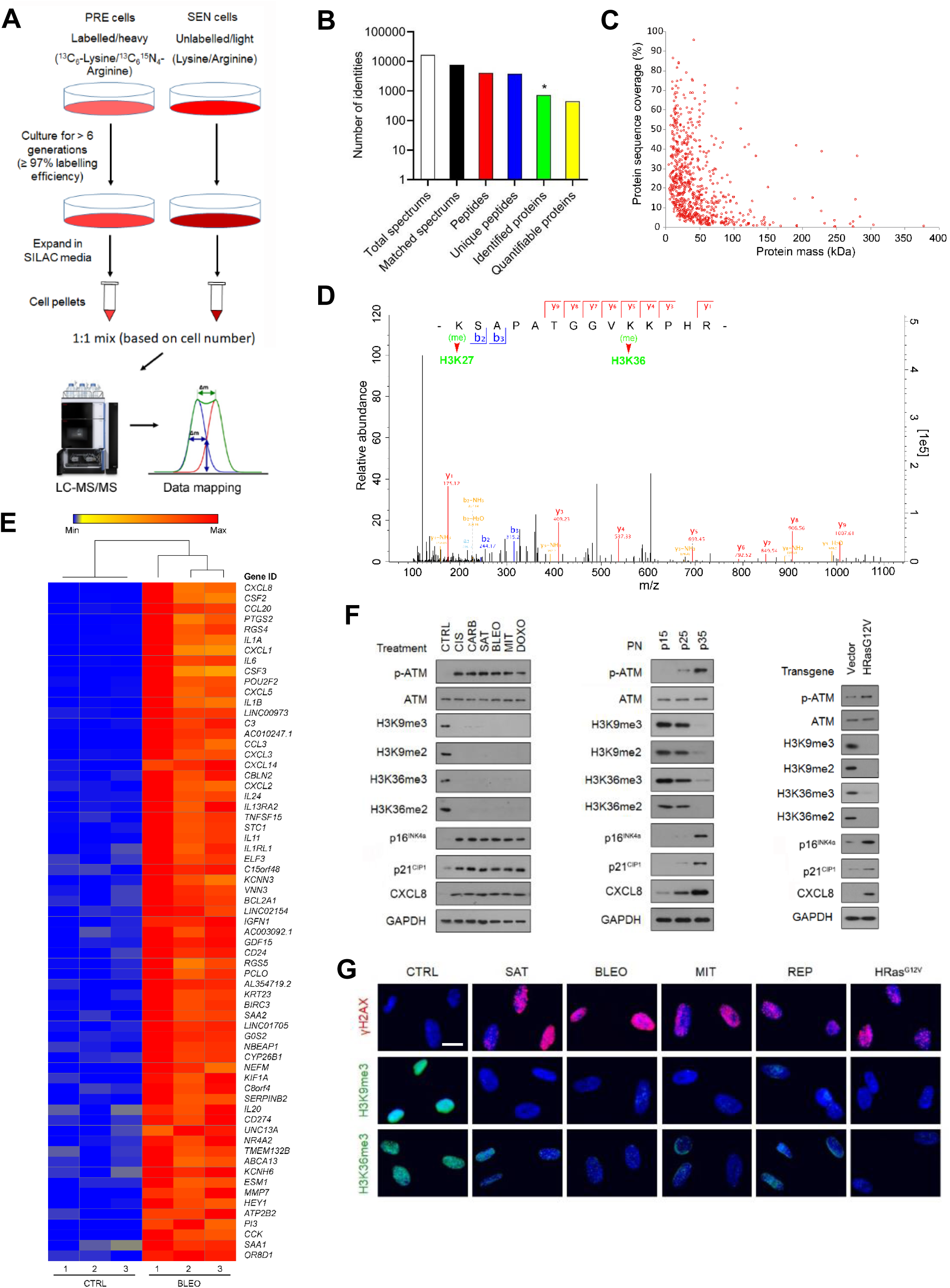
Histone H3 Lysine Sites Are Epigenetically Modified upon Cellular Senescence. A. Technical scheme of SILAC-based identification of intracellular proteins in presenescent *versus* senescent cells of the human stromal line PSC27. PRE cells, presenescent cells. SEN cells, senescent cells. B. Column statistics of different categories of protein molecules in output data after SILAC analysis. *, identified proteins (732). C. Scatterplot of proteins identified by SILAC procedure. Protein sequence coverage was plotted against protein mass (447 quantifiable). D. A representative plot derived from characterization of tandem mass spectrometry (MS/MS) based quantitative proteomics profiling. For MS scans, the m/z scan range was 100 to 1100. Intact peptides were detected in the Orbitrap at a resolution of 70,000. E. Heatmap depicting genes significantly upregulated in SEN cells after bleomycin treatment. CTRL, control. BLEO, bleomycin. Genes are ordered by their expression fold change (highest on top) in PRE *versus* SEN cells after RNA-seq. F. Immunoblot analysis of key molecules in DNA damage repair, cellular senescence and the SASP in PSC27 cells induced to senescence by chemotherapeutic agents (TIS), replicative exhaustion (RS), or oncogene activation (OIS). PN, passage number. p15, p25, p35 representing different passages in culture. Vector, empty control for human HRas^G12V^. H3K9me2/3 and H3K36me2/3, histone H3 methylation markers. G. Immunofluorescence staining of γH2AX, H3K9me3, and H3K36me3 in PSC27 cells after chemotherapeutic treatment (SAT, BLEO, and MIT), replicative exhaustion (REP), or oncogene activation (HRas^G12V^). SAT, satraplatin. BLEO, bleomycin. MIT, mitoxantrone.

Among the differentially expressed proteins we identified, 87 showed a significant increase and 29 displayed a significant decrease in abundance in SEN cells (> 2-fold, *P* < 0.05), respectively (Supplementary Table 1). However, when analyzing PTM sites differentially modified between PRE and SEN cells, we noticed reduction of histone H3.2 signals in the categories of both dimethylated and trimethylated proteins, namely H3K27 and H3K36 (Fig. 1D) (Supplementary Table 3). As histone PTMs can alter the architecture of chromatin and are implicated in epigenetic regulation of senescent cell-associated phenotypes ^21^, we interrogated the possibility of systematic or general changes at histone H3 sites, and whether they are causal for specific consequences during cellular senescence.

To address these questions, we primarily focused on the major forms of PTMs occurring at histone H3.2, including methylation. As it is known that the level of H3K27me3, a repressive histone mark mainly enriched in heterochromatin, tends to diminish in senescent human fibroblasts *via* an autophagy/lysosomal pathway ^22, 23^, here we paid special attention to H3K36, whose demethylation and resulting implications are less reported in the case of cellular senescence ^24^. To substantiate the data from proteomics assays, we performed whole transcriptomic assessment with RNA sequencing (RNA-seq). The data showed a typical SASP expression profile, as evidenced by upregulation of multiple pro-inflammatory factors and activation of senescence-associated signaling pathways (Fig. 1E and Extended Data Fig. 1A), confirming entry into cellular senescence after exposure to agent-delivered genotoxicity. Expression assessment of classic histone-specific methylases and demethylases revealed a distinct upregulation pattern of epigenetic factors, specifically the KDM4 family (including members A, B, C and D), although the significance of only A and B was confirmed by quantitative RT-PCR (Extended Data Fig. 1B-1C). To build on these findings, we used several chemotherapeutic drugs including cisplatin (CIS), carboplatin (CARB), satraplatin (SAT), mitoxantrone (MIT), and doxorubicin (DOXO), in addition to BLEO to treat PSC27 cells (Fig. 1F, left). Additionally, we exposed cells to replicative senescence (RS) and oncogene-induced senescence (OIS, HRas^G12V^) (Fig. 1F, middle and right, respectively). In each case, we observed prominent cellular senescence (Extended Data Fig. 1D-1E). Immunoblot analysis indicated phosphorylation of ATM, enhanced expression of p16^INK4a^, p21^CIP1^, and CXCL8 (IL8, a hallmark SASP factor), but a diminished methylation pattern (including tri- and di-levels) at the H3K36 site (Fig. 1F). As KDM4 selectively demethylates both H3K9 and H3K36, an activity implicated in key cellular processes including DDR, cell cycle regulation, and senescence ^25^, we examined H3K9 methylation in parallel. Not surprisingly, H3K9 exhibited a demethylation tendency largely resembling that of H3K36 throughout all cell-based assays (Fig. 1F).

Upon immunofluorescence (IF) staining, we observed markedly reduced signals of both H3K9me3 and H3K36me3, sharply contrasting with γH2AX, a canonical marker of DDR foci at DSBs, the expression of which was apparently enhanced in DNA-damaged PSC27 cells (Fig. 1G). The data suggested that loss of epigenetic PTM marks at histone H3.2, specifically those imaging H3K9 and H3K36 methylation status, is accompanied by increased expression of KDM4 family members (mainly A/B), and occurs consistently in senescent cells as a general phenomenon, though caused by an unknown mechanism and with unclear biological consequences.

### Stromal expression of KDM4A/B in the damaged TME correlates with adverse clinical outcomes

Given the rising expression of KDM4A/B in senescent cells under *in vitro* conditions, we asked whether the results are reproducible in clinical settings.

We first investigated the biospecimens derived from a cohort of prostate cancer (PCa) patients, and observed remarkable upregulation of KDM4A/B in prostate tumor tissues after chemotherapy, specifically in contrast to those collected before treatment (Fig. 2A). Interestingly, upregulated KDM4A/B were generally localized in the stromal compartments rather than their adjacent cancer epithelium, which appeared to have limited or no signals.

**Fig 2.**
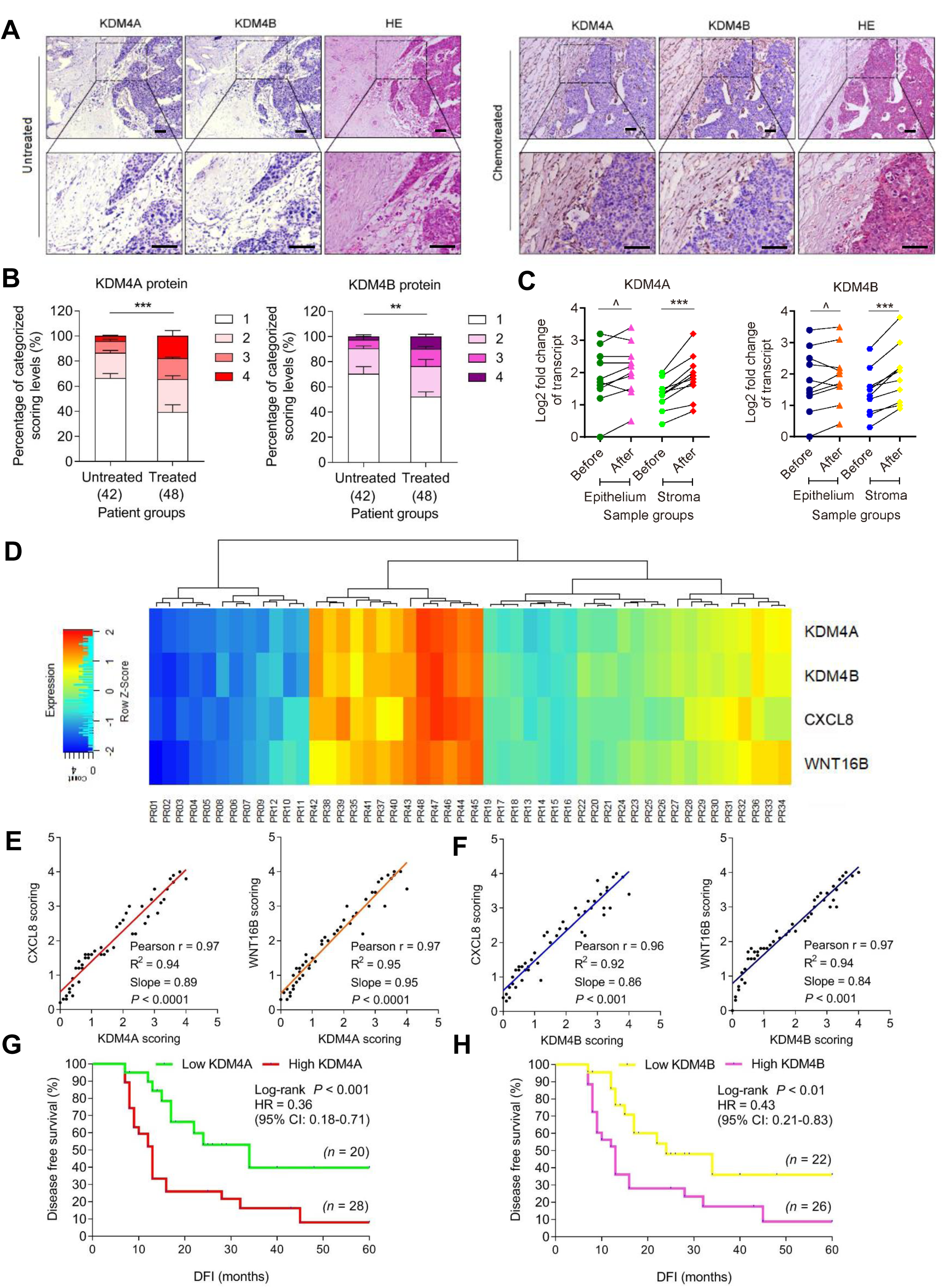
KDM4A and KDM4B Are Expressed in Human Prostate Tumor Stroma and Correlated with Adverse Clinical Survival. A. Histological images of KDM4A/B expression in human prostate cancer (PCa) tissues. Left, before chemotherapy. Right, after chemotherapy. In each panel, images of left and middle columns were from immunohistochemical (IHC) staining, those in the right column from hematoxylin and eosin (HE) staining. Rectangular regions in the upper images amplified into corresponding lower images. Scale bars, 100 μm. B. Pathological assessment of stromal KDM4A/B expression in PCa tissues (42 untreated *versus* 48 treated). Left, KDM4A. Right, KDM4B. In each group, patients were pathologically assigned into 4 categories *per* IHC staining intensity of the antigen in stroma. 1, negative; 2, weak; 3, moderate; 4, strong expression. C. Comparative analysis of KDM4A/B expression at transcription level between before and after chemotherapy. Data are organized for epithelial and stromal cells, respectively, after laser capture microdissection (LCM) of either cell lineage followed by quantitative analysis. Each dot represents an individual patient, with the data of “before” and “after” connected to allow direct profiling of KDM4A/B expression in the same individual patient. D. The landscape of pathological correlation between KDM4A/B, CXCL8, and WNT16B in the stroma of PCa patients after chemotherapy. Scores derived from assessment of molecule-specific IHC staining, with expression levels colored to reflect low (blue) *via* modest (turquoise) and fair (yellow) to high (red) signal intensity. Columns represent individual patients, rows different molecules. Totally 48 patients posttreatment were analyzed, with scores of each patient averaged from 3 independent pathological readings. E. Statistical correlation between pathological scores of KDM4A and CXCL8/WNT16B (Pearson analysis, r = 0.97 for both; *P* < 0.0001 for both) in the 48 tumors with matching protein expression assessment data. F. Statistical correlation between pathological scores of KDM4B and CXCL8/WNT16B (Pearson analysis, r = 0.96, r = 0.97, respectively; *P* < 0.0001 for both) in the 48 tumors described in (E). G. Kaplan-Meier analysis of PCa patients. Disease-free survival (DFS) stratified according to KDM4A expression (low, average score < 2, green line, n = 20; high, average score ≥ 2, red line, n = 28). DFS represents the length (months) of period calculated from the date of PCa diagnosis to the point of first disease relapse. Survival curves generated according to the Kaplan-Meier method, with *P* value calculated using a log-rank (Mantel-Cox) test. H. Kaplan-Meier analysis of PCa patients. Disease free survival (DFS) stratified according to KDM4B expression (low, average score < 2, yellow line, n = 22; high, average score ≥ 2, pink line, n = 26). DFS represents the length (months) of period calculated from the date of PCa diagnosis to the point of first disease relapse. Survival curves generated according to the Kaplan-Meier method, with *P* value calculated using a log-rank (Mantel-Cox) test. Data in all bar plots are shown as mean ± SD and representative of 3 biological replicates. ^, *P* > 0.05. *, *P* < 0.05. **, *P* < 0.01. *** *P* < 0.001.

Expression of KDM4A/B in prostate tissues post-*versus* pre-chemotherapy was quantitatively substantiated by a pre-established pathological evaluation protocol for precise assessment of a target protein expression according to its immunohistochemistry (IHC) staining intensity (*P* < 0.001) (Fig. 2B). To confirm their inducible nature *in vivo*, we analyzed a subset of patients whose pre- and post-chemotherapy samples were both accessible, by performing transcript assays upon laser capture microdissection (LCM) of each cell lineage. The data showed significant upregulation of KDM4A/B in the stroma, but not their nearby cancer epithelium in the posttreatment biospecimens (*P* < 0.001 *versus P* > 0.05) (Fig. 2C). We further noticed the expression dynamics of KDM4A/B in damaged TME essentially parallel those of IL8 and WNT16B, two hallmark SASP factors (Fig. 2D). The correlation between KDM4A/B and IL8/WNT16B expression in the damaged TME was further supported by pathological evaluation of their expression scores in posttreatment patients (Fig. 2E-F). More importantly, Kaplan-Meier analysis of PCa patients stratified according to KDM4A/B expression in tumor stroma suggested a significant, albeit negative correlation between KDM4A/B and disease-free survival (DFS) in the treated cohort (*P* < 0.001 and *P* < 0.01, respectively, log-rank test) (Fig. 2G-2H).

We then analyzed the situations of KDM4C/D, the other members of KDM4 subfamily. Pathological data suggested that neither molecule was upregulated in tumor stroma after chemotherapy, while expression profiling *per* cell lineage showed no changes in stromal or epithelial compartments (Extended Data Fig. 2A-2D). Furthermore, pathological stratification according to KDM4C/D expression in tumor stroma failed to reveal potential correlations between KDM4C/D and patient DFS in the treated cohort (*P* > 0.05, log-rank test) (Extended Data Fig. 2E-2F).

Given the prominent induction of KDM4A/B in tumor stroma of posttreatment cancer patients, we questioned if there were possible changes in methylation levels of H3K9 and H3K36, two major targets of demethylases KDM4A/B in human cells. IHC assessment of patient samples suggested attenuated H3K9me3 and H3K36me3 signals in stromal cells, whereas neighboring cancer cells in the disease foci appeared largely unaffected (Extended Data Fig. 2G-2J). LCM-supported comparative expression assays *per* cell lineage confirmed the declining pattern of H3K9me3 and H3K36me3 in stromal cells, but not their epithelial counterparts in the TME posttreatment (Extended Data Fig. 2K). Of note, there was a largely reverse correlation between KDM4A/B expression and H3K9/H3K36 trimethylation as indicated by pathological evaluation of target signals in posttreatment samples (Extended Data Fig. 2L). Pearson statistical analysis further substantiated these inherent links (Extended Data Fig. 2M). As a special point, we noticed the lower level of H3K9/H3K36 trimethylation in stromal compartments and the shorter survival of PCa patients in the post-chemotherapy stage (Extended Data Fig. 2N-2O), implying a potential mechanism associated with these epigenetic marks and driving adverse consequences, specifically mortality in cancer clinics.

### Upregulated KDM4A/B expression is accompanied by decreased H3K9/H3K36 methylation in senescent cells

We next performed a time course expression assay with stromal cell lysates collected at individual time points after genotoxicity-induced senescence. In contrast to DDR signaling, which appears to be an acute response, KDM4A/B protein levels gradually increased within the first 7 d posttreatment before entering a plateau. KDM4C was slightly enhanced upon DNA damage but seemingly declined after the first 6 h until approaching a stable phase, whereas KDM4D was essentially undetectable at the protein level throughout the assays (Fig. 3A). Transcription analysis of KDM4 largely reproduced the data from immunoblots, with KDM4A/B levels progressively increasing in senescent cells, while KDM4C/D did not (Fig. 3B top). Of note, the expression pattern of KDM4A/B was indeed similar to that of SASP hallmark factors (IL6, CXCL8) and cyclin-dependent kinase inhibitors (p16^INK4a^, p21^CIP1^) (Fig. 3B bottom).

**Fig 3.**
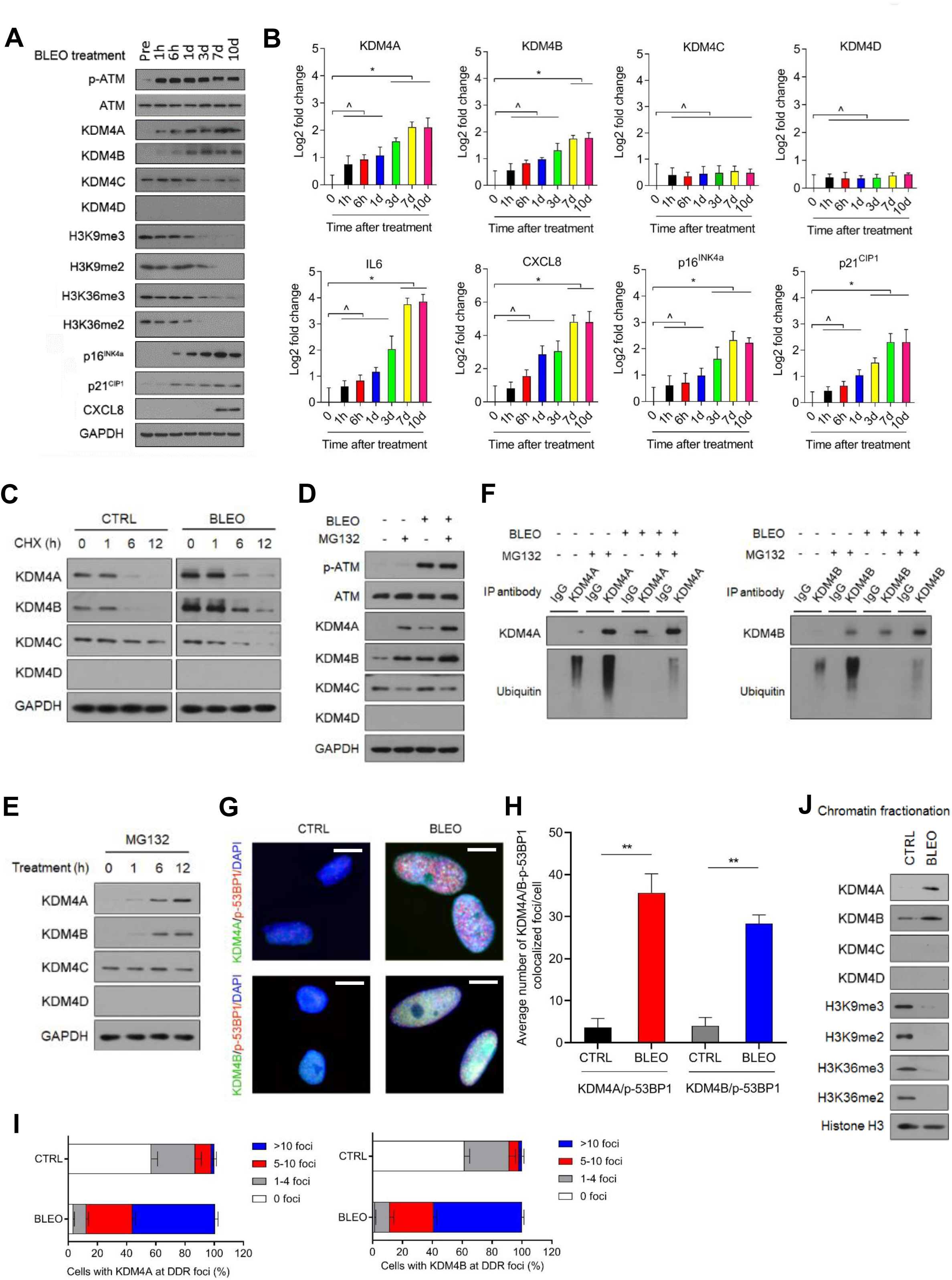
KDM4 Expression Is Regulated Post-Translationally in Senescent Cells. A. Time course analysis of KDM4 expression in PSC27 cells after treatment by bleomycin. Cell lysates were collected at indicated time points posttreatment and subject to immunoblot assays. GAPDH, loading control. B. Time course measurement of transcript expression of KDM4 subfamily members and IL6, CXCL8, p16^INK4a^, and p21^CIP1^ in stromal cells after exposure to genotoxicity. C. Immunoblot analysis of KDM4 expression in stromal cells treated by BLEO and/or cyclohexamide (CHX). Cell lysates were collected at indicated time points. D. Immunoblot assessment of KDM4 expression in stromal cells treated by BLEO and/or MG132. Cell lysates were collected at indicated time points. E. Immunoblot appraisal of KDM4 protein levels in PSC27 cells treated by MG132. Cell lysates were collected at indicated time points. F. Evaluation of KDM4 protein posttranslational modification *via* immunoprecipitation (IP) followed by immunoblot assays. PSC27 cells were treated by BLEO and/or MG132 and lysed for analysis 7 days post *in vitro* damage. Anti-KDM4A/B was used for IP, with precipitates subject to immunoblot assay with KDM4A/B antibody. Anti-ubiquitin was used to probe ubiquitination profile of KDM4A/B. G. Immunofluorescence staining of KDM4A/B and p-53BP1 in stromal cells. Cells were treated by BLEO and subject to immunofluorescence staining 7 days later. KDM4A/B, green. p-53BP1, red. Nuclei (DAPI), blue. Scale bars, 5 μm. H. Comparative statistics of the average number of foci where KDM4A/B and p-53BP1 are colocalized in damaged PSC27 per cell. I. Statistics of stromal cells displaying nuclear co-localization of KDM4A/B and p-53BP1 in control (CTRL) *versus* senescent (SEN) cells. Data represent the percentage (%) after immunofluorescence-based counting. J. Immunoblot assessment of KDM4 and H3K9/H3K36 methylation after chromatin fragmentation. Histone H3, loading control for nuclear lysates. Data in all bar plots are shown as mean ± SD and representative of 3 biological replicates. ^, *P* > 0.05. *, *P* < 0.05.

To investigate the mechanism supporting KDM4 protein levels, we treated cells with cyclohexamide (CHX), a protein synthesis inhibitor. In contrast to control cells, BLEO-induced senescent cells exhibited increased KDM4A/B expression, while their signals in each case diminished in the time course upon CHX treatment, suggesting KDM4A/B are subject to protein turnover in these cells (Fig. 3C). In the presence of MG132, a cell-permeable proteasome inhibitor, protein levels of KDM4A/B were elevated, thus confirming that both factors are indeed vulnerable to proteasome-mediated degradation (Fig. 3D). Distinct from the case of KDM4C/D, which remained largely unchanged, MG132 markedly increased KDM4A/B levels, thus further implying their proteasome-regulated nature (Fig. 3E). Subsequently, immunoprecipitation (IP) from total lysates of senescent cells indicated enhanced KDM4A/B upon MG132 treatment, but with decreased intensities of ubiquitin-mediated PTM, suggesting that the increase of KDM4A/B proteins in senescent cells is partially correlated with a ubiquitin/proteasome-escaping mechanism (Fig. 3F).

Upon IF-based cell staining, we noticed enhanced levels of KDM4A/B in the nuclei of PSC27 (Fig. 3G). Interestingly, both factors are substantially co-localized with p-53BP1, one of the canonical markers of DDR foci, signals of which displayed a significant increase in DNA-damaged cells (Fig. 3G-3H).

Notably, the percentage of cells detected with KDM4A/B at DDR foci was considerably enhanced after BLEO treatment (Fig. 3I). To establish the subcellular localization of KDM4A/B upon cellular senescence, we examined cell lysates with chromatin fractionation. Immunoblot analysis indicated that both factors were significantly elevated in the chromatin fractions of senescent cells relative to their proliferating counterparts (Fig. 3J). Thus, KDM4A/B are regulated by mechanisms involving both PTM-associated and ubiquitin/ proteasome-eluding pathways, with a potential but yet unknown epigenetic role in the nuclear compartment of senescent cells.

### KDM4A/B functionally regulate the SASP without affecting senescence

We next interrogated the relevance of KDM4A/B to SASP development in senescent cells. Data from cell transduction assays showed that KDM4A overexpression did not change the basal level of SASP factors, while after DNA damage expression of the majority of SASP factors was further enhanced by ectopic KDM4A (Fig. 4A). This tendency was validated at protein level by immunoblots, as reflected by enhanced CXCL8 expression in senescent cells, a process accompanied by further reduced H3K9/H3K36 methylation (tri- and di) intensities in the presence of exogenous KDM4A (Fig. 4B).

**Fig 4.**
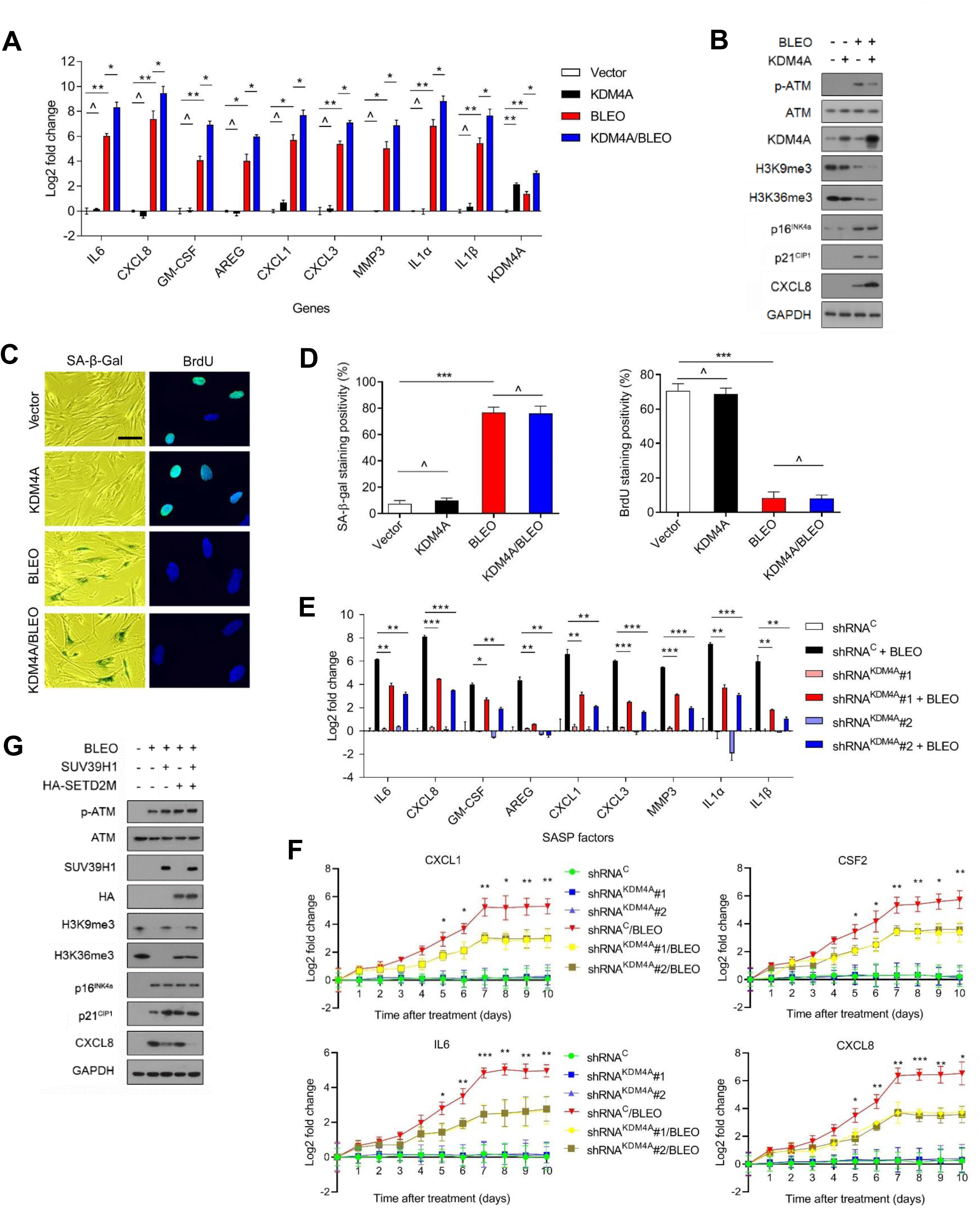
SASP Expression Is Enhanced but H3K9/H3K36 Methylation Is Attenuated by the Histone Demethylase KDM4A. A. Quantitative assessment of SASP expression at the transcription level. PSC27 cells were transduced with a lentiviral construct encoding human KDM4A and/or exposed to BLEO treatment before being collected for expression assays. Signals normalized to CTRL cells (transduced with empty vector and untreated). B. Immunoblot assay of DNA damage repair (DDR) signaling, H3K9/H3K36 methylation, and SASP expression in cells processed in different ways as described in (A). GPADH, loading control. C. Representative images of SA-β-Gal and BrdU staining of PSC27 cells subject to treatment as described in (A). D. Comparative statistics of staining results of stromal cells in the individual conditions of (A). Left, SA-β-Gal staining. Right, BrdU staining. E. Transcript expression of hallmark SASP factors in PSC27 sublines transduced with lentiviral constructs encoding KDM4A-specific shRNAs. Scrambled, shRNA control. Cells were subject to vehicle or BLEO treatment before collected for quantitative analysis. F. Expression curves of hallmark SASP factors in stromal cells treated in conditions as described in (E). Cells were lysed at the indicated time points after BLEO damage. G. Immunoblot assays of stromal cells transduced with lentiviral constructs encoding human SUV39H1 (full length), SETD2M (histone methylase domain SET, tagged with HA) or both. Cells were subject to BLEO treatment after transduction, with cell lysates collected 7-10 days later. GAPDH, loading control.

As KDM4A/B share a similar domain architecture including two PHD and two TUDOR domains, we speculated that the influence of KDM4A on the SASP can be basically phenocopied upon KDM4B transduction. This hypothesis was substantiated in assays performed with KDM4B, as evidenced by a similar set of transcript and protein data (Extended Data Fig. 3A-3B). When cells were exposed to Chaetocin, a histone methyltransferase inhibitor against SUV39H1, which preferentially catalyzes H3K9 methylation (H3K9me2/me3), the data partially resembled those of KDM4A/B expression (Extended Data Fig. 3C). Interestingly, however, expression of p16^INK4a^ and p21^CIP1^, the key CDKIs indicative of cellular senescence, remained largely unaffected upon transgenic expression of KDM4A/B, suggesting that cell cycle arrest, or alternatively, cellular senescence, is likely not regulated by these epigenetic factors. To validate this assumption, cellular senescence and cell cycle arrest assays with SA-β-Gal staining and BrdU incorporation assays were performed, respectively. The resulting data supported that expression of neither KDM4A nor KDM4B was sufficient to influence cellular senescence or replication (Fig. 4C-4D and Extended Data Fig. 3D-3E).

We then selectively eliminated these two factors from stromal cells with shRNA, after which cells were subject to BLEO-induced senescence. Again, removal of KDM4A/B abrogated expression of the SASP (Fig. 4E and Extended Data Fig. 3F). We evaluated the induction pattern of several hallmark SASP factors including CXCL8, CSF2, CXCL1, and IL6 by graphing time-dependent curves, and noticed fold change of these factors significantly restricted by KDM4A/B depletion, starting from cellular senescence induction (Fig. 4F). Interestingly, however, development of cellular senescence *per se* was not affected upon knockdown of KDM4A/B (Extended Data Fig. 3G-3H).

Next, we queried the influence of histone methyltransferases SUV39H1 and SETD2, the latter specifically responsible for trimethylation of the H3K36 site (H3K36me3), on cellular senescence and the SASP. To this end, we transduced the full length human SUV39H1 and the methyltransferase domain of human SETD2 (SETD2M, which encodes aa 946-1738 as the SET domain and mediates SETD2 catalytic function) ^26^, individually or simultaneously before genotoxic treatment. Immunoblots indicated that ectopic expression of these epigenetic factors failed to abrogate cellular senescence, with p21^CIP1^ expression being even slightly enhanced (Fig. 4G). Importantly, however, each of these methyltransferases was able to reduce the SASP expression, as evidenced by diminished CXCL8 signals, although concurrent transduction of SUV39H1 and SETD2M generated the most dramatic effect. Thus, epigenetic modification of certain sites of chromatin histone H3.2, specifically trimethylation of H3K9 and H3K36, hold the potential to restrain the intensity of secretome while sustaining cell cycle arrest of senescent cells.

### Targeting the demethylase activity of KDM4 restrains the SASP

Given the critical role of KDM4A/B in SASP development, we interrogated whether pharmacologically targeting their demethylase activities can effectively control the SASP in senescent cells. To address this, we employed ML324, a small molecule compound selectively inhibiting the activities of KDM4 family ^27^. RNA-seq data suggested that expression of the majority of SASP factors in senescent cells was markedly diminished upon ML324 treatment, including but not limited to CXCL8, CSF2, CCL20, IL1A, CXCL1, and IL6 (Fig. 5A). Among the genes significantly upregulated upon cellular senescence, a considerable portion showed reduced expression when cells were exposed to ML324, although some SASP-unrelated genes were also affected by this agent, such as PTGS2, RGS4, and POU2F2 (Fig. 5A). GO analysis showed that the most suppressed pathways and biological processes by ML324 were correlated with extracellular secretion, NF-κB signaling, receptor tyrosine phosphorylation, and the MAPK cascade, activities generally characteristic of canonical SASP factors (Fig. 5B). Further bioinformatics validated an overlapping zone comprising 129 genes between those upregulated upon cellular senescence and those downregulated upon ML324-mediated KDM4 inhibition, the latter evidenced by rescued methylation of H3K9/H3K36 sites (tri- and di-forms) and accompanied by diminished CXCL8 expression (Fig. 5C and Extended Data Fig. 4A). Gene set enrichment analysis (GSEA) confirmed that restraining KDM4A/B activities effectively dampened the SASP and disturbed NF-κB activity (Fig. 5D and Extended Data Fig. 4B). However, ML324 treatment did not interfere with cell colony formation or cell growth arrest (Fig. 5E and Extended Data Fig. 4C). Interestingly, a number of genes whose expression declined in senescent cells, appeared to be upregulated upon cell exposure to ML324 (Extended Data Fig. 4D), suggesting that KDM4A/B inhibition alters expression of a spectrum of genes, among which are those encoding the SASP factors, and affected genes are not limited to those upregulated in senescent cells. Although the mechanism of ML324 in reversing the expression of many SASP-canonical and a handful of SASP-non-canonical genes remains yet unclear, further work to define the impact of KDM4 deficiency on the transcriptome-wide expression profile of senescent cells is warranted.

**Fig 5.**
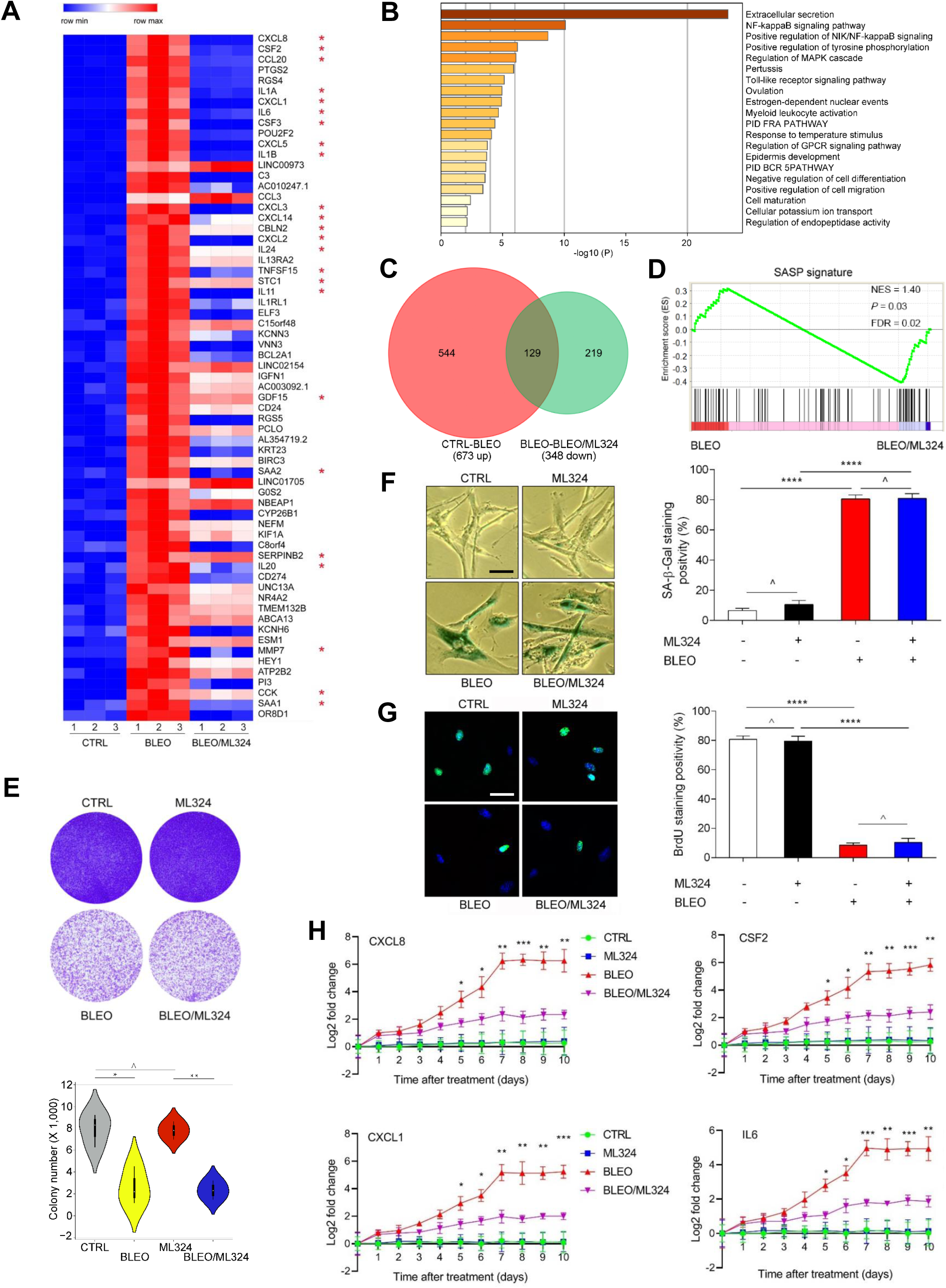
Targeting KDM4 with a Small Molecule Inhibitor Interferes with the SASP but Not Cell Growth or Cell Cycle Arrest. A. Heatmap depicting the influence of DNA damage and ML324, a selective chemical inhibitor of KDM4, on transcriptomic expression profile of PSC27 cells. Genes sorted by expression fold change when comparing between cells treated by CTRL *versus* BLEO (highest on top). Red stars, canonical SASP factors affected by ML324. B. Graphic visualization of pathways by GO profiling, significantly enriched genes were those downregulated and sorted according to their fold change when senescent cells were exposed to ML324. C. Venn diagram presentation of genes upregulated by BLEO (673, in relative to CTRL) and downregulated genes by ML324 (348, in relative to BLEO). D. GSEA profiling of gene expression with significant enrichment scores showing a SASP-specific signature in BLEO/ML324 co-treated cells compared with BLEO only-treated cells. E. *In vitro* colony formation assay of cells exposed to BLEO and/or ML324 treatment. Upper, representative images of crystal violet staining. Lower, comparative statistics. F. SA-β-Gal staining of cells after treatment by BLEO and/or ML324. Left, representative images. Right, statistics. G. BrdU staining of cells treated as described in (E). Left, images. Right, statistics. H. Time course expression of a subset of SASP factors (CXCL8, CSF2, CXCL1, and IL6). Cells were subject to BLEO and/or ML324 treatment.

*In vitro* assays including SA-β-Gal staining and BrdU incorporation suggested that ML324 did not affect cellular senescence or cell cycle arrest, consistent with the findings that KDM4A/B are not essential for the maintenance of cellular senescence *per se* (Fig. 5F-5G). Gene expression curves substantiated that ML324 significantly restricted expression of SASP factors, as exemplified by the cases of CXCL8, CSF2, CXCL1, and IL6, with the efficacy evident throughout the course of cellular senescence induction (Fig. 5H). To define further the ML324-generated SASP-targeting consequence, specifically in a virtual microenvironment, we utilized a co-culture system that involves treatment of cancer cells with stromal cell-derived conditioned media (CM). Notably, the capacity of proliferation, migration, and invasion of PC3, DU145, LNCaP, and M12, which are typical prostate cancer (PCa) cell lines sharing the same organ origin as the PSC27 stromal line, namely human prostate, considerably decreased upon ML324-mediated treatment of stromal cells (Extended Data Fig. 4E-4G). More importantly, resistance of PCa cells to MIT, a chemotherapeutic agent frequently administered to PCa patients in cancer clinics, was significantly reduced when stromal cells were treated by ML324 (Extended Data Fig. 4H). Thus, suppression of the demethylase activity of KDM4 with a small molecule inhibitor can retard SASP development without compromising cellular senescence, resulting in a diminished potential of the SASP to promote cancer progression.

### Chromatin remodeling supports development of the SASP which can be uncoupled from cellular senescence by targeting KDM4

The state of chromatin dictates vital cellular processes such as DNA repair and gene expression, while accessible chromatin marks can decorate regulatory sequences including enhancers, promoters and locus-control regions to cooperatively regulate gene expression ^28^. We next sought to determine the possibility and significance of chromatin structure alterations in supporting SASP development in senescent cells. Since accessible regions can be mapped by assay for transposase-accessible chromatin with high throughput sequencing (ATAC-seq), and co-localize extensively with enhancers and promoters of target genes, we used this epi-strategy to define genome-wide accessible regions in senescent cells.

We first investigated whether the accessible chromatin landscape of senescent cells differs from that of their proliferating counterparts. Surprisingly, we observed unusually strong ATAC-seq signals upstream (∼0.5 kb) of transcription end sites (TESs) of transcriptionally active genes in senescent cells induced by BLEO (Fig. 6A). Although there was a sharp increase in signals at approximately 2.0 kb upstream of transcription start sites (TSSs), the difference between senescent and proliferating control cells seems to be limited at their proximal regions of TSSs. However, in the presence of the KDM4 inhibitor ML324, we observed a remarkable decrease of enrichment signals at both TSSs and TESs of senescent cells (Fig. 6A), suggesting transcriptional activation of these senescence-associated genes was basically reversed. The TES-open chromatin may reflect the binding of factors involved in transcriptional termination, while these sites may alternatively function as enhancers that promote high level transcription of biologically essential genes ^29^. Our data suggest that open chromatin can be found both at promoters and near TES regions of transcriptionally active genes in senescent cells.

**Fig 6.**
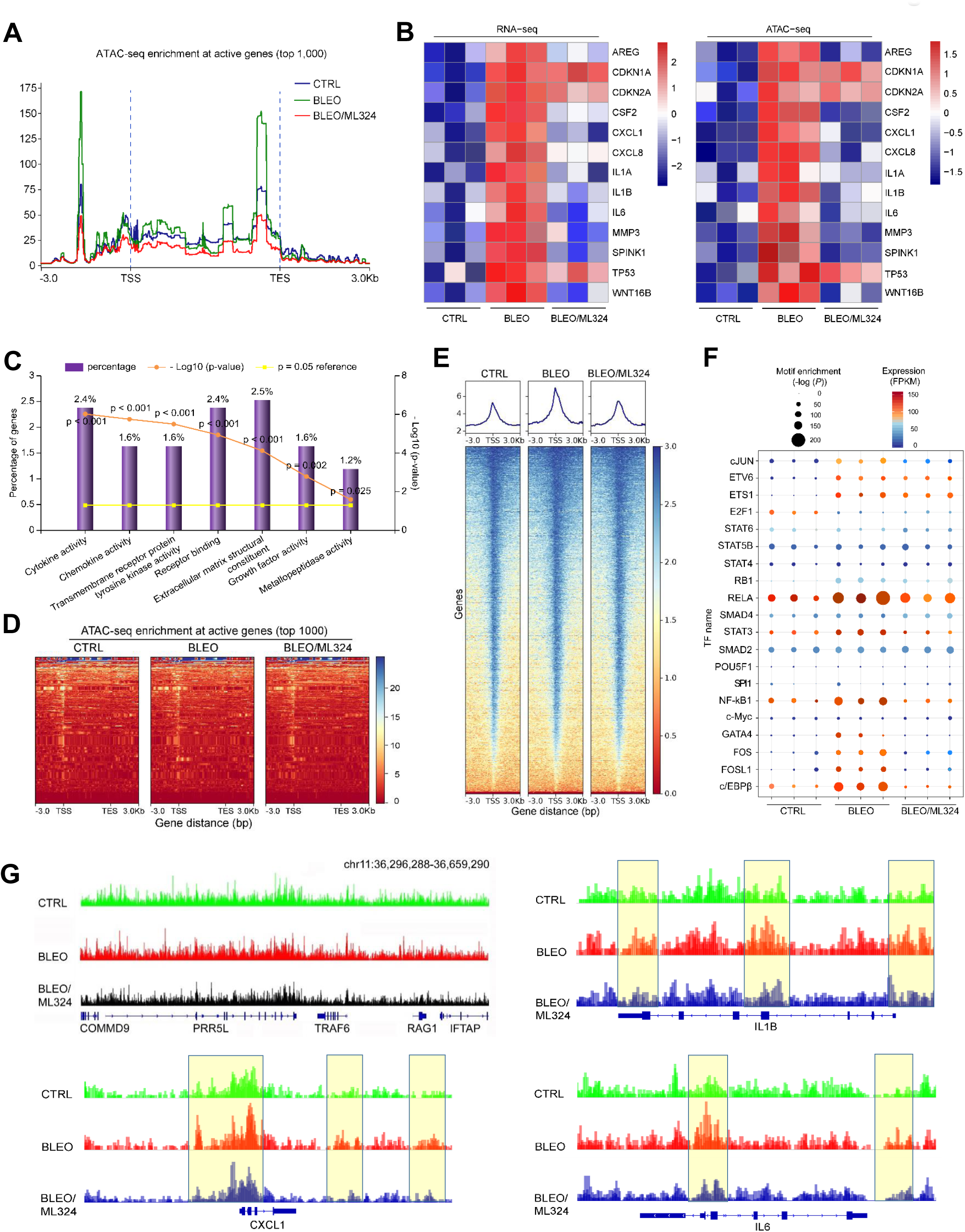
Accessible Chromatin Landscape in Senescent Cells and ML324-Mediated Suppression. A. The average level of ATAC-seq enrichment (normalized) for the top 1,000 most active genes between proliferating cells, senescent cells (bleomycin-induced), and bleomycin/ML324 co-treated cells (marked as CTRL, BLEO, and BLEO/ML324, respectively). B. Heatmaps showing the expression (FPKM) of the SASP hallmark genes (RNA-seq, left) and the assessable chromatin enrichment (RPKM) (ATAC-seq, right) at their promoters (TSS ± 2.5 kb). Example genes are listed alongside each type of heatmap. C. Gene ontology (GO) analysis results for gene classes of significant expression fold change in proliferating *versus* senescent cells, and inhibited substantially upon ML324 treatment. Percentage of gene number among all upregulated genes in senescent cells, log10 of p value *per* class presented, with *P* < 0.05 as significance. D. Heatmap showing ATAC-seq enrichment of peaks near the accessible promoters (3.0 kb upstream of TSS and downstream of TES *per* gene) in each of the assayed samples. Enrichment signals were collected for all active TSSs and TESs, which were assorted by cap analysis of gene expression (CAGE) values, with peaks defined by hierarchical clustering. The top 1,000 genes of enrichment signals were selected for analysis. E. Heatmap depicting ATAC-seq enrichment of peaks near the accessible promoters (3.0 kb upstream TSS and downstream TES *per* gene, respectively) present in each of the assayed samples. The whole genomic range was evaluated *per* sample. F. Transcription factor (TF) motifs identified from distal ATAC-seq peaks in each group’s samples. Only TFs expressed detectable (FPKM ≥ 5) and motif enrichment *P* value < 1 × 10^-10^ in each sample were included. G. The UCSC browser views show enrichment of ATAC-seq signals at the promoters of several SASP-unrelated genes (COMMD9, PRR5L, TRAF6, RAG1, and IFTAP) in contrast to those at the promotes of representative SASP factors (CXCL1, IL-1β, and IL6).

We next questioned whether chromatin accessibility at the enhancers and promoters of SASP factors is generally consistent with their individual expression activity. Results from data mapping with ATAC-seq and RNA-seq outputs indicate that transcriptional levels of a set of typical SASP factors were intimately correlated with intensities of their promoter ATAC-seq signals, though those of CDK inhibitors (e.g. CDKN1A, CDKN2A and TP53) remained essentially unchanged (Fig. 6B). Bioinformatics profiling suggested that the vast majority of genes upregulated upon cellular senescence but downregulated upon ML324 treatment have implications in cytokine, chemokine, transmembrane receptor binding, extracellular matrix structure, growth factor, and metalloproteinase-associated activities (Fig. 6C).

Enrichment-based heatmaps illustrating either the most active 1,000 genes (∼3.0 kb upstream of TSS and downstream of TES *per* gene, respectively), or whole transcriptomics, displayed striking difference between control and senescent cells, as well as between senescent cells exposed to vehicle and exposed to ML324 (Fig. 6D-6E and Extended Data Fig. 5A). As enhancers represent hotspot sites for transcription factor (TF) binding ^30^, we reasoned that distal ATAC-seq peaks may harbor motifs for TFs that regulate cellular senescence and/or related phenotypes. Using the motif analysis program HOMER, we extracted the binding motifs for an array of TFs enriched in distal peaks found in either of the 3 experimental conditions of PSC27 cells (Fig. 6F). Notably, some of these TFs, including AP-1 family members (cJUN, FOS and FOSL1), RELA (p65 of NF-κB), STAT3, NF-κB1 (p50/p105 of NF-κB), GATA4, and c/EBPβ, which have been reported to be SASP-correlated ^31–35^, clearly showed a ‘senescence-up and ML324-down’ pattern. However, certain TFs such as ETV6, ETS1 and RB1, exhibited enhanced binding capacity in senescent cells, a tendency seemingly not affected upon KDM4 suppression. Other TFs such as E2F1, manifested reduced activity in senescent cells and were not rescued by ML324 treatment. Together, senescent cells showed distinct landscapes for distal ATAC-seq peaks, while significant gene expression is correlated with a handful of ‘master TFs’ regulating the senescence-specific circuitry and potently reset upon KDM4 deficiency induced by ML324. In addition, we noticed there were a set of SASP-unrelated genes whose ATAC-seq signals were substantially decreased in senescent cells but apparently reversed upon cell exposure to ML324 (Extended Data Fig. 5B), suggesting the influence of KDM4 inhibition may not be exclusively limited to chromatin accessibility changes underlying SASP expression.

To validate the data further, we performed footprinting analysis of accessible chromatin for specific genes. The reproducibility between biological replicates of our ATAC-seq data was first confirmed with a small group of genes, including COMMD9, PRR5L, TRAF6, RAG1, and IFTAP, which are presumably not correlated with cellular senescence (Extended Data Fig. 5C). Next, we assessed the data from allelic ATAC-seq enrichment assays and uncovered open genomic regions of typical SASP factors, including IL1β, CXCL1, IL6, AREG, SPINK1, MMP3, and WNT16B, in senescent cells, but with a markedly reduced accessibility on the chromatin upon KDM4 suppression (Fig. 6G and Extended Data Fig. 5D). In contrast, the chromatin openness of genes not correlated with senescence remained essentially unaltered, such as COMMD9, PRR5L, and TRAF6 (Fig. 6G). As a special control, CDKN2A and TP53 were used for parallel analysis and showed enhanced accessibility in senescent cells, but were largely sustained when cells were exposed to ML324 treatment (Extended Data Fig. 5E).

Thus, chromatin accessibility and transcriptional expression are intimately correlated for SASP hallmark factors, while active regulatory elements are accessible by the cell’s expression machinery, such as a special set of key TFs. Although the transition of accessible chromatin landscapes during cellular senescence enables site-specific transcription, transposable elements can remodel these chromatin landscapes in the case of KDM4 deficiency in senescent cells.

### Therapeutically targeting KDM4 minimizes chemoresistance and enhances preclinical index

Given the prominent role of the epigenetic factor KDM4 in the development of cellular senescence-associated phenotypes, specifically the SASP, we next reasoned the potential of selectively harnessing this target to improve therapeutic efficacy of age-related disorders. As cancer represents the leading cause of age-standardized incidence and premature mortality in the global range and is adversely correlated with the impact of senescent cells ^36, 37^, we chose the TME as a pathological model for subsequent *in vivo* epigenetic manipulation.

First, we generated tissue recombinants by admixing stromal cells (PSC27) with cancer cells (PC3) at a pre-optimized ratio before subcutaneous implantation into the hind flank of experimental mice with non-obese diabetes and severe combined immunodeficiency (NOD/SCID). To closely simulate clinical conditions, we designed a preclinical regimen that incorporates the genotoxic agent MIT and/or the KDM4-specific inhibitor ML324 (Fig. 7A). Two weeks after implantation, when stable uptake of tumors *in vivo* was observed, a single dose of therapeutic agent or placebo (vehicle) was delivered to animals at the first day of 3^rd^, 5^th^, and 7^th^ week until end of the 8-week regimen (Extended Data Fig. 6A). Upon measurement of tumors at end of the preclinical trial, we found MIT administration alone caused significant reduction in tumor size (42.8%, *P* < 0.001), while addition of ML324 further decreased tumor mass (54.5%, *P* < 0.001), allowing an overall shrinkage of 74.0% (Fig. 7B). Alternatively, bioluminescence imaging (BLI) of xenografts generated with PC3 cells stably expressing luciferase (PC3-luc) and PSC27 cells excluded the potential metastasis of cancer cells from the primary sites, with signal intensities largely in line with tumor growth patterns we observed in PC3/PSC27 animals (Fig. 7C). The data suggest that classic chemotherapy combined with a KDM4-targeting agent can induce tumor regression with an index significantly higher than that of chemotherapy alone.

**Fig 7.**
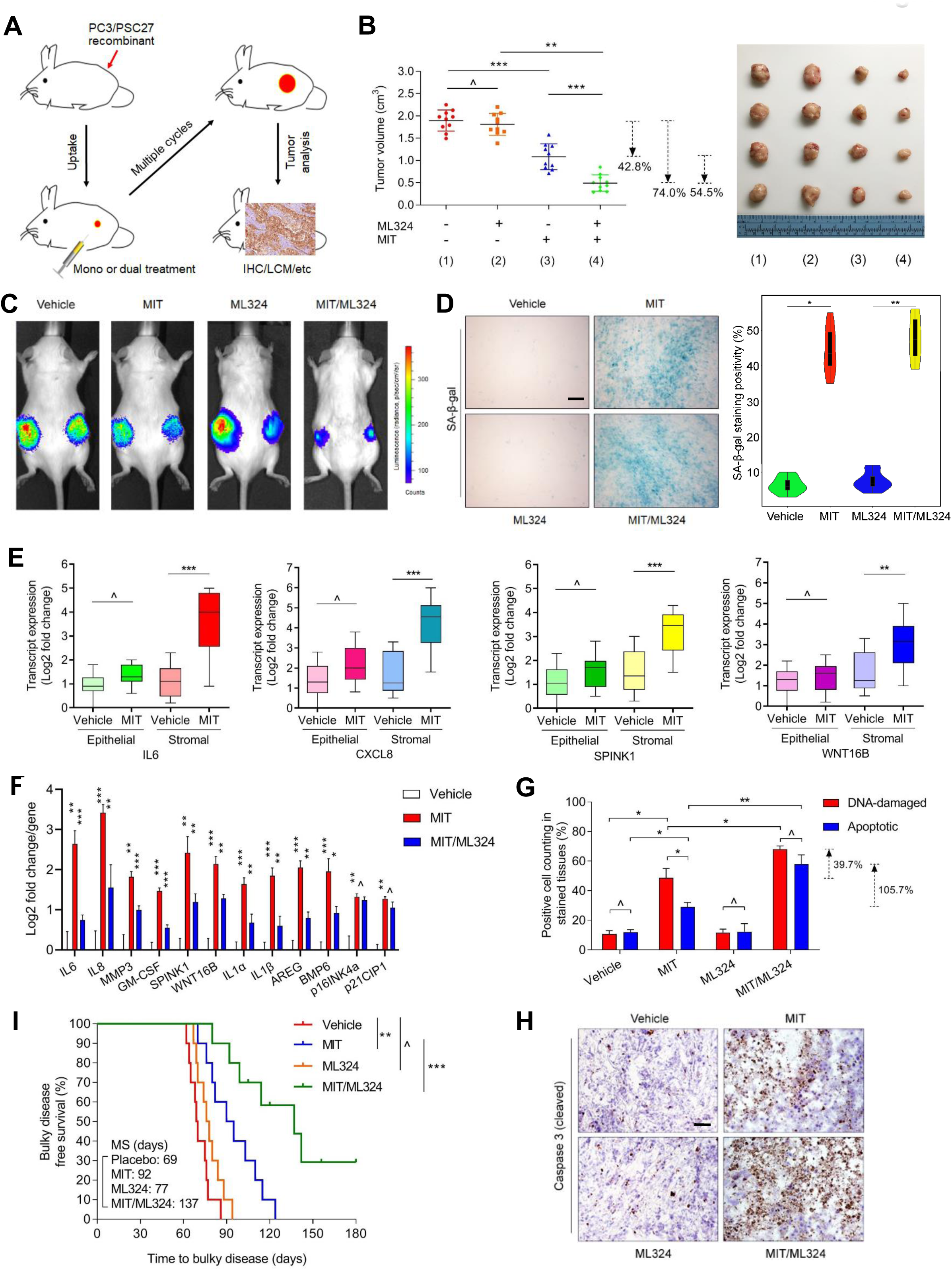
Therapeutically Targeting KDM4 in the Damaged TME Diminishes Cancer Resistance Conferred by Senescent Stroma. A. Illustrative diagram for preclinical treatment of non-obese diabetes and severe combined immunodeficient (NOD/SCID) mice. Two weeks after subcutaneous implantation and *in vivo* uptake of tissue recombinants, animals received either single (mono) or combinational (dual) agents administered as metronomic treatments composed of several cycles. B. Statistical profiling of tumor end volumes. PC3 cells were xenografted alone or together with PSC27 cells to the hind flank of NOD/SCID mice. The chemotherapeutic drug MIT was administered to induce tumor regression, alone or together with the KDM4 inhibitor ML324. Right, representative images of tumors. C. Representative bioluminescence imaging (BLI) of PC3/PSC27 tumor - bearing animals in the preclinical trial. Digital signals were proportional to *in vivo* luciferase activities measured by an IVIS device. D. Comparative imaging of *in vivo* senescence of tumor tissues by SA-β-gal staining. Tumors were freshly dissected upon animal sacrifice and processed as frozen sections before tissue staining. Scale bars, 200 μm. Right, violin plot of positivity statistics. E. Transcript assessment of the *in vivo* expression of several canonical SASP (including IL6/CXCL8/SPINK1/WNT16B) factors in stromal cells isolated from the tumors of NOD/SCID mice. Tissues from animals xenografted with both stromal and cancer cells were subject to LCM isolation, total RNA preparation and expression assays. F. Quantitative appraisal of SASP factor expression in stromal cells isolated from tumor tissues of animals subject to different treatments. Signals *per* factor were normalized to the vehicle-treated group. G. Statistical assessment of DNA-damaged and apoptotic cells in the biospecimens collected as described (B). Values are presented as percentage of cells positively stained by IHC with antibodies against γ-H2AX or caspase 3 (cleaved). H. Representative IHC images of caspase 3 (cleaved) in tumors at the end of therapeutic regimes. Biopsies of vehicle-treated animals served as negative controls for MIT-treated mice. Scale bars, 100 μm. I. Survival appraisal of mice which were sacrificed upon development of advanced bulky diseases. Survival duration was calculated from the time of tissue recombinant injection until the day of death. Data analyzed by log-rank (Mantel-Cox) test.

When examining the potential off-target effects of therapeutic agents on the TME, we noticed a considerable percentage of senescent cells in the foci of mice treated by agents involving MIT, presumably resulting from genotoxicity-induced tissue damage (Fig. 7D). However, ML324 itself did not cause such a consequence, which is basically consistent with its targeting mechanism. Upon LCM-assisted and cell lineage-specific isolation of stromal and cancer cell subpopulations individually from harvested tumor samples, we evaluated transcription levels of specific genes. The resulting data indicated significantly increased expression of SASP factors including but not limited to IL6, IL8, SPINK1, WNT16B, IL1α, MMP3, and GM-CSF, a tendency accompanied by remarkable upregulation of p16^INK4a^ and p21^CIP1^ in tissues dissected from MIT-treated animals (Fig. 7E and Extended Data Fig. 6B). However, these changes seem to be generally limited to stromal cells, rather than their adjacent epithelial cell counterparts, which have repopulated upon development of drug resistance in the course of treatment as previously characterized ^31, 38^. We further assessed the *in situ* inducibility of KDM4A/B in tumor samples, and noticed substantial expression of these factors in animals experiencing MIT-involved treatments (Extended Data Fig. 6C-6D), essentially in line with our *in vitro* findings (Fig. 3A-3B). Together, these data suggest incidence of *in vivo* cellular senescence accompanied by development of a typical SASP and expression of KDM4A/B, a side effect induced by genotoxicity during chemotherapeutic intervention and mainly observed in the benign components of the TME.

We next analyzed the *in vivo* efficacy of ML324 by appraising the level of hallmark SASP factors between these sample groups. Strikingly, expression of the vast majority of SASP components was significantly reduced when comparing animals treated by MIT alone and those by MIT/ML324, although the senescence markers p16^INK4a^ and p21^CIP1^ remained unchanged (Fig. 7F).

These data thus consolidate our *in vitro* findings that KDM4 inhibition retards SASP development, but not cellular senescence, a phenomenon correlated with differential regulation of KDM4-mediated histone H3 demethylation, which enables chromatin reorganization.

To investigate the mechanism directly responsible for MIT-induced cancer resistance, we dissected tumors from animals treated by different agents 7 days after treatment, a time point prior to the emergence of resistant colonies. In contrast to vehicle treatment, MIT administration caused dramatically enhanced indices of DNA damage and apoptosis, a pattern not observed in tumors dissected from ML324-treated animals (Fig. 7G). We noticed that ML324 alone induced neither typical DDR nor substantial cell death. However, upon combinational treatment with MIT and ML324, we observed maximal intensities of both DNA damage and cell apoptosis within tissues, suggesting a therapeutic efficacy superior to those when each agent was used alone (Fig. 7G). Histological appraisal with immunohistochemical (IHC) staining against caspase 3 (cleaved), a typical biomarker of cell apoptosis, largely confirmed the different responses of tumors (Fig. 7H).

Given the remarkable benefit of targeting KDM4 in the treatment-damaged TME, we subsequently evaluated longer time consequences by comparing the survival terms of animals subject to different treatment modalities in a time-extended manner. Development of a bulky disease was considered once the tumor burden was prominent (*e.g.*, when the size exceeds 2000 mm^3^), an approach reported previously ^39, 40^. Mice receiving MIT/ML324 co-treatment exhibited the most prolonged median survival duration, acquiring a minimally 50% longer survival than the group treated by MIT alone (Fig. 7I, green *versus* blue). Notably, however, ML324 alone did not provide significant benefit, conferring only marginal survival elongation (Fig. 7I, orange *versus* red). Thus, targeting KDM4 in the TME neither changes tumor growth nor animal survival, while MIT/ML324 combinational treatment significantly improves both parameters.

We then interrogated the safety of such treatment regimens by assessing individual *in vivo* indices. Experimental data suggest that administration of these therapeutic agents was well tolerated by mice, as no significant perturbations in body weight, renal function (creatinine, urea) or liver integrity (ALP and ALT) were observed (Extended Data Fig. 6E-6F). Further, the chemotherapeutic agents and ML324 provided at the doses used in this study did not significantly interfere with organ metabolism, the immune system or tissue homeostasis, even in immune-competent animals (C57BL/6 strain, Extended Data Fig. 6G-6I). These results together substantiate that ML324, a typical KDM4 inhibitor, combined with conventional chemotherapy holds the potential to promote tumor response without generating severe and systemic cytotoxicity in critical organs.

## Discussion

Senescent cells accumulate in diverse organ types with age and progressively reside in damaged or dysfunctional tissues. Besides exiting cell cycle, senescent cells undergo multiple phenotypic alterations such as morphological abnormality, metabolic reprogramming, and chromatin reorganization ^41^. They actively synthesize and release a large number of soluble factors, collectively termed the SASP, which mediate most of senescence-associated cell-non-autonomous effects ^8, 42^. Senescent cells have attracted increasing attention as a potent therapeutic target for age-related diseases including cancer, one of the leading causes of morbidity and mortality in the elderly ^5^. Considering the beneficial effects of cellular senescence in special circumstances such as those in tissue repair, wound healing, and embryonic development ^6, 7^, however, it may be more advantageous to specifically restrain the SASP, rather than radical elimination of these cells from tissue microenvironments in many situations. In this study, we unraveled an epigenetic mechanism functionally supporting SASP development and defined the key target KDM4 to manipulate senescent cells by dampening the SASP while retaining cell cycle arrest. We further established an effective strategy to improve therapeutic outcomes of age-related pathologies, such as cancer, by integrating classic chemotherapy and KDM4-specific agents as a solution guided by advanced concepts.

Epigenetic patterns of mammalian cells can fluctuate over the organismal lifespan, and are subject to influence of multiple factors including those from interior and/or exterior sources. Epigenetic alterations representing heritable changes and affecting gene expression or cellular phenotype without DNA sequence changes, have emerged as a major type of pathophysiological event ^43^. Covalent modifications of chromatin constituents including DNA and histones, are key regulators of genome-wide expression, while histone methylation, which occurs at lysine and arginine residues, causes either gene activation or silencing in multiple events, depending on methylation residues of the histones ^44, 45^. A global loss or redistribution of heterochromatin may be a common feature of organismal aging and correlated with increased genomic instability and altered gene expression ^12^. However, the biological links between epigenetic changes, cellular senescence, and senescence-specific phenotypes remain poorly understood. We mapped the epigenetic landscape of senescent cells using a proteomics approach (SILAC) to reveal enhancement of KDM4 expression and its relevance in the demethylation of histone residuals specifically H3K9/H3K36 upon cellular senescence. As a genome-wide survey of accessible chromatin in senescent cells remains largely lacking, we were tempted to dissect to what extent gene expression is correlated with epigenomic reprogramming. The data support that gene activation and chromatin openness can occur, at least in part, through pathways associated with epigenetic modification and chromatin remodeling, consistent with data from recent epigenetic studies ^29, 46^. Thus, global transcription can be newly activated, once the open chromatin is established in senescent cells, while broadly accessible chromatin may contribute to the pervasive transcription through regulatory modules, including those driving SASP expression. The study not only unveiled the chromatin landscapes in senescent cells, but also allowed genome-wide identification of regulatory circuitry in the course of epigenomic remodeling, an event that does occur upon cellular senescence (Fig. 8).

**Fig 8.**
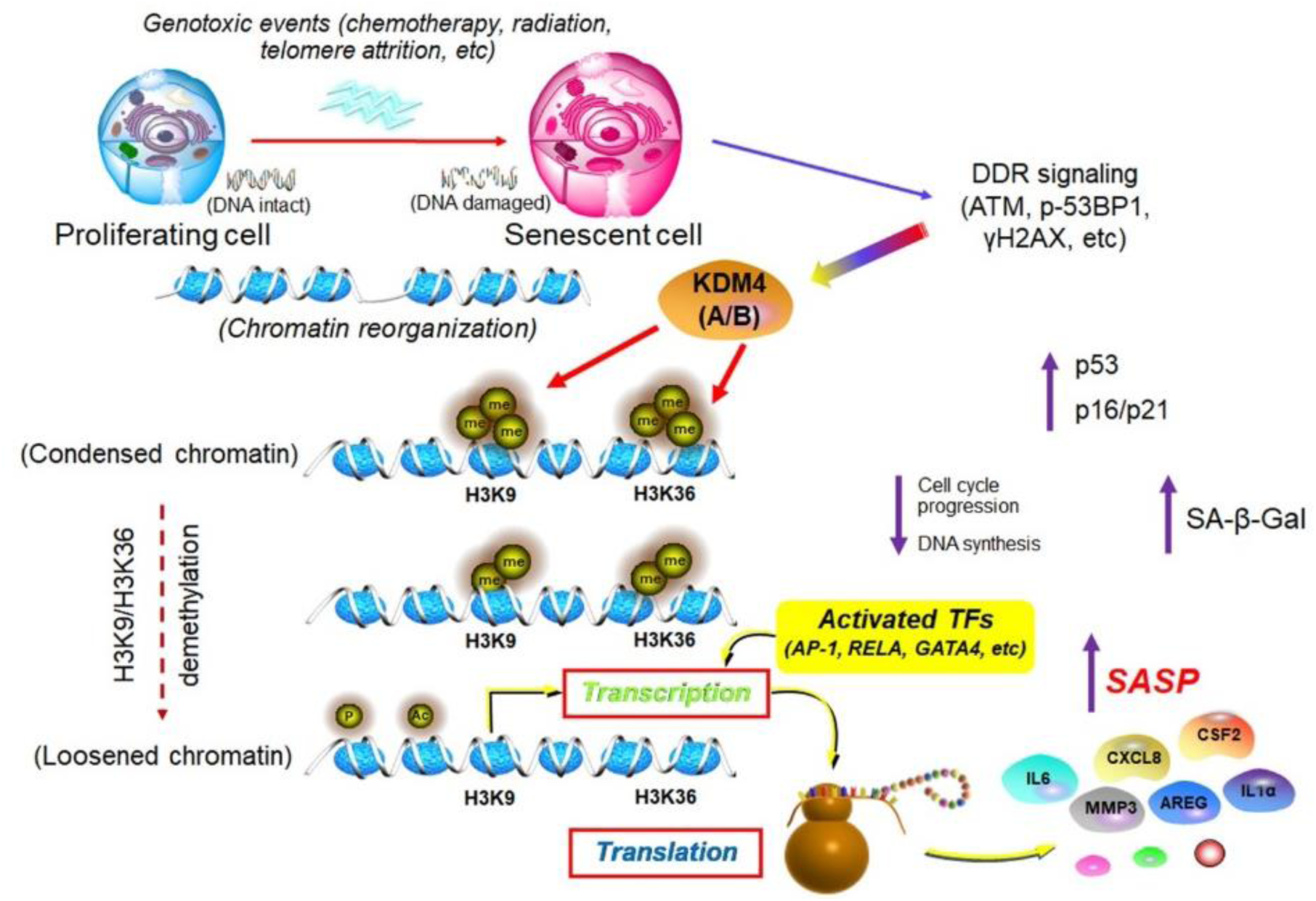
Working model depicting epigenomic reprogramming and KDM4-mediated histone demethylation (H3K9/H3K36), events allowing SASP expression in senescent cells. In genotoxic settings, senescent cells undergo irreversible DNA damage which triggers enhanced DDR signaling. Chromatin accessibility landscape is remodeled, with a set of activated transcription factors physically binding to the enhancers and promoters of senescence-associated genes including those encoding SASP factors. There is a strong concordance of clustering scheme and a close functional linkage between chromatin accessibility and transcriptional output. Future efforts to combine genome and epigenome sequencing, as well as to generate maps of chromosome conformation, will pave the way to tackling the non-coding genome in senescent cells. DDR, DNA damage repair. TF, transcription factor. SASP, senescence-associated secretory phenotype.

Physical access to DNA is a highly dynamic property of chromatin, playing an essential role in development and maintenance of a certain cellular state, albeit frequently displaying a cell type-dependent pattern ^47^. Intricate organization of accessible chromatin across the genome reflects a network of permissible interactions through which enhancers, promoters, insulators, and chromatin-binding factors cooperatively modulate gene expression. This landscape of accessibility changes dynamically during cellular senescence in response to environmental or interior stimuli, while the homeostatic maintenance of accessibility dynamically is regulated through a competitive interplay between chromatin-binding factors and nucleosomes ^28^. Through parallel assessment of signal intensities of canonical SASP factors between ATAC-seq and RNA-seq data, we observed the strongest concordance of the clustering scheme with RNA abundance and enrichment at promoters and enhancers of these genes, consistent with an inherent functional linkage between chromatin accessibility and transcriptional output. Further, the exquisite distal regulatory elements identified from our ATAC-seq data enabled the discovery of formerly unappreciated players in senescent cells. With this data-rich resource, we identified classes of central TFs actively driving the SASP in senescent cells, with expression of these TFs correlated with distinct patterns in TF occupancy and motif binding. Together, integration of RNA-seq and ATAC-seq with paired datasets enables a quantitative model to link the accessibility of a regulatory factor to the expression of predicted target genes, the promoters and enhancers of which tend to be bound by special TFs upon chromatin reorganization. However, despite such a precise workflow, profiling of chromosome conformation in senescent cells has not yet been performed, and future efforts to generate maps of chromosome conformation would further improve our understanding of genome-wide regulatory networks upon cellular senescence, and correspondingly, gain deeper insights into human aging and age-related diseases.

Chemotherapy is one of the mainstream anticancer modalities. However, much like other types of treatments, the effectiveness of chemotherapeutics is limited by drug resistance, which can be divided into two broad categories: intrinsic and acquired. In contrast to intrinsic resistance, which pre-exists in cancer cells prior to initiation of therapy, acquired resistance arises during intervention and is frequently conferred by the tumor adjacent stroma ^48^. Typically, the stroma comprises heterogeneous cell subpopulations of mesenchymal origin, mainly including fibroblasts, smooth muscle cells, endothelial cells, and immune cells, while interactions between cancer cells and stromal cells are essential for cancer progression ^49^. For instance, cancer resistance can be driven by a treatment-damaged stroma, a process pathologically fueled by the SASP ^20, 50, 51^. To date, increasing lines of evidence support that the SASP has broad implications in human cancers. Although highly context-dependent, the SASP has a wealth of consistent functions in the TME, including those involved in cancer initiation, growth, metastasis, and even relapse ^52^. In contrast to studies that indicate the beneficial effects of therapy-induced senescence (TIS) induction in cancer cells, including tumor growth control and the SASP-mediated immune response promoting senescent cancer cell clearance, therapy-triggered off-target effects on the surrounding benign stroma can have undesirable outcomes during and after cancer treatment, particularly when cancer cells develop acquired resistance against subsequent cycles of drug administration.

KDM4 is a family of histone demethylases that regulate chromatin structure and govern gene expression by demethylating special histone H3 sites including H3K9 and H3K36 ^53^. KDM4 members are overexpressed or deregulated in multiple cancer types, cardiovascular diseases and mental retardation as potential broad-spectrum therapeutic targets ^54^. Deregulated KDM is associated with chromatin instability, tumor suppressor silencing, oncogene activation, hormone receptor binding and downstream signaling ^54^.

Despite extensive investments, however, there are few KDM4-selective inhibitors that can be exploited, presumably due to the structural similarity of KDM members and their conserved properties of the active site ^55^. Therefore in this case, precise defining of the pathological functions of KDM4 will provide the foundation for discovery of novel and potent inhibitors, which many be designed to target alternative sites. Our study reveals that expression of KDM4A/B is significantly upregulated in senescent stromal cells, a distinct feature responsible for diminished H3K9/H3K36 methylation. Importantly, this work unmasks the critical role of epigenetic factors specifically KDM4 in supporting SASP development upon cellular senescence and provides an exemplifying framework for targeting the SASP to control age-related pathologies such as cancer, specifically by manipulating senescent cells in an epigenetic manner. A recent study investigating structurally unrelated histone demethylases in melanoma, interesting, reported that KDM1A and KDM2C can disable oncogene (Ras or Braf)-induced senescence by enhancing expression of E2F target genes in mouse embryo fibroblasts (MEFs) and exert oncogenic potential by overcoming the senescence barrier to full-blown tumorigenesis, suggesting the divergent roles of histone demethylases in different settings ^56^. Based on TCGA data compiled from cohorts of human cancer patients (Extended Data Fig. 7A-7D), we further highlight KDM4 as a multifaceted target, and provide proof-of-principle evidence that it can be exploited to restrain the pathological impact of a treatment-damaged TME that harbors myriad senescent cells, a therapeutic approach exemplified by preclinical trials.

## Methods

### Cell lines, *in vitro* culture and lentivirus

Human primary prostate stromal cell line PSC27 was kindly provided by Dr. Peter Nelson (FHCRC) and maintained in PSC complete medium as described previously ^20^. Prostate cancer cell lines PC3, DU145 and LNCaP were from ATCC and routinely cultured in DMEM (10% FBS plus 1% penicillin /streptomycin). M12, a neoplastic and metastatic prostate cancer cell line, was a gift from Dr. Stephen Plymate (UW) ^57^. All cell lines were routinely tested for microbial contamination and authenticated with STR assays.

Lentiviral particles were produced using Lipofectamine 2000 and a packaging kit (Thermal Scientific) based on manufacturer’s instructions. PSC27 infected with viruses encoding the puromycin resistance gene were selected using puromycin (1 μg/ml) for 3 days.

### Reagents, plasmids and antibodies

The following antibodies were purchased from the indicated suppliers and used for immunoblotting at indicated concentrations: rabbit monoclonal anti-p-ATM (Abways cat. no. CY5111), 1:1000; rabbit monoclonal anti-ATM (Abways cat. no. CY5207), 1:1000; rabbit polyclonal anti-H3K36me3 (ABclonal cat. no. A23660), 1:1000; rabbit monoclonal anti-H3K9me3 (abcam cat. no. ab8898), 1:1000; rabbit monoclonal anti-p21 (Abways cat. no. CY5088), 1:1000; rabbit monoclonal anti-p16 (Abways cat. no. CY5357), 1:1000; rabbit polyclonal anti-IL-8 (Proteintech cat. no. 17038-1-AP), 1:1000; rabbit monoclonal anti-GAPDH (Abways cat. no. AB0037), 1:2000; rabbit monoclonal anti-KDM4A (Abways cat. no. CY8322; Abcam cat. no. ab24545), 1:1000; rabbit monoclonal anti-KDM4B (Cell Signaling cat. no. 8639), 1:1000; rabbit polyclonal anti-KDM4C (abcam cat. no. ab85454), 1:1000; rabbit polyclonal anti-KDM4D (abcam cat. no. ab198209), 1:1000; rabbit monoclonal anti-H3 (Cell Signaling cat. no. 4499), 1:1000; rabbit monoclonal anti-SUV39H1 (Cell Signaling cat. no. 8729), 1:1000; rabbit monoclonal anti-HA (Cell Signaling cat. no. 3724), 1:1000.

The pLenti-Puro, pLVX-IRES-Puro and pLKO.1-Puro plasmids were purchased from Addgene. Lentiviral vector pLKO.1-Puro was used to clone small hairpin RNAs (shRNAs): KDM4A shRNA1 (target sequence: GCACCGAGTTTGTCTTGAAAT); KDM4A shRNA2 (target sequence: TTCGAGAGTTCCGCAAGATAG); KDM4A shRNA3 (target sequence: TAGTGAAAGGACGAGCCATTT); KDM4B shRNA1 (target sequence: GCGGCAGACGTATGATGACAT); KDM4B shRNA2 (target sequence: GCGGCATAAGATGACCCTCAT); KDM4B shRNA3 (target sequence: GATGACCTTGAACGCAAATAC). KDM4A and SUV39H1 cDNAs were cloned to pLenti-Puro as expression constructs, while KDM4B and HA-SETD2-M cDNAs were cloned to pLVX-IRES-Puro respectively.

### *In vitro* cell treatments and senescence appraisal

Stromal cells were grown until 60%-80% confluent (CTRL) and treated with bleomycin (50 μg/ml, BLEO, MedChemExpress, cat. no. HY-17565), mitoxantrone (1 uM, MIT, Selleck, cat. no. S2485), doxorubicin (2 uM, DOXO, MedChemExpress, cat. no. HY-15142), cisplatin (10 μM, CIS, MedChemExpress, cat. no. HY-17565), carboplatin (20 μM, CARB, MedChemExpress, cat. no. HY-17393), satraplatin (20 μM, SAT, MedChemExpress, cat. no. HY-17576), docetaxel (100 nM, DTX, TOCRIS, cat. no. 4056), paclitaxel (100 nM, PTX, MedChemExpress, cat. no. HY-B0015), vincristine (50 nM, VCR, Selleck, cat. no. S1241) for 12 h. After treatment, the cells were rinsed twice with PBS and allowed to stay for 7-10 d in media.

Oncogenic HRAS^G12V^-transduced cells were selected by puromycin for 3 d and allowed to stay for 7-10 d in media. Alternatively, the cells were allowed to passage consecutively for replicative exhaustion. SA-β-Gal staining assay was performed with a commercial kit (Beyotime) to examine cellular senescence. For cell cycle arrest, a single pulse of BrdU (10 μM, Yeasen) was performed for 24 h and cells were subject to immunofluorescence staining with an anti-BrdU antibody (1:400 dilution, Cell Signaling, cat no. 5292) before counterstained with DAPI and assessed by fluorescence microscopy.

DNA-damage extent was evaluated by immunostaining for γH2AX or p53-BP1 foci by following a 4-category counting strategy as formerly reported ^20^. Random fields were chosen to show DDR foci, and quantified using CellProfiler (http://www.cellprofiler.org). For clonogenic assays, cells were seeded at 1 × 10^3^ cells/dish in 10mm dishes for 24 h before treated with chemicals. Cells were fixed with 2% paraformaldehyde 7-10 d post treatment, gently washed with PBS and stained instantly with 10% crystal violet prepared in 50% methanol. Excess dye was removed with PBS, with plates photographed. Colony formation were evaluated by quantifying the number of single colonies per dish.

### Cancer patient recruitment and clinical studies

Chemotherapeutic administration involving genotoxic agents was performed for primary prostate cancer patients (Clinical trial no. NCT03258320) by following the CONSORT 2010 Statement (updated guidelines for reporting parallel group randomized trials). Patients with a clinical stage ≥ I subtype A (IA) (T1a, N0, M0) of primary cancer but without manifest distant metastasis were enrolled into the multicentered, randomized, double-blinded and controlled pilot studies. Age between 40-75 years with histologically proven prostate cancer was required for recruitment into the individual clinical cohorts. Data regarding tumor size, histologic type, tumor penetration, lymph node metastasis, and TNM stage were obtained from the pathologic records. Before chemotherapy, tumors were acquired from these patients as ‘Pre’ samples (an ‘Untreated’ cohort). After chemotherapy, remaining tumors in patients were acquired as ‘Post’ samples (a ‘Chemo-treated’ cohort, with most tumors collected within 1-6 months after treatment). For some cases, the ‘Pre’ and ‘Post’ tumor biopsies from the same individual patient were both accessible, and these samples were subject to further evaluation. Tumors were processed as FFPE biospecimens and sectioned for histological assessment, with alternatively prepared OCT-frozen chunks processed via lase capture microdissection (LCM) for gene expression analysis. Specifically, stromal compartments associated with glands and adjacent to cancer epithelium were separately isolated from tumor biopsies before and after chemotherapy using an Arcturus (Veritas Microdissection) laser capture microscope following previously defined criteria ^20^. The immunoreactive scoring (IRS) gives a range of 1–4 qualitative scores according to staining intensity per tissue sample. Categories for the IRS include 0-1 (negative), 1-2 (weak), 2-3 (moderate), 3-4 (strong) ^58^. The diagnosis of prostate cancer tissues was confirmed based on histological evaluation by independent pathologists. Randomized control trial (RCT) protocols and all experimental procedures were approved by the Ethics Committee and Institutional Review Board of Shanghai Jiao Tong University School of Medicine and Zhongshan Hospital of Fudan University, with methods carried out in accordance with the official guidelines. Informed consent was obtained from all subjects and the experiments conformed to the principles set out in the WMA Declaration of Helsinki and the Department of Health and Human Services Belmont Report.

### Chromatin fractionation

To assess the levels of KDM4 family members at the chromatin-bound fraction, CTRL and BLEO-treated PSC27 cells were cross-linked with 1% formaldehyde for 5 min and quenched with 0.125 M glycine; then, cells were washed thrice with 1 × PBS, and incubated with buffer A (10 mM HEPES, pH 7.9, 10 mM KCl, 1.5 mM MgCl2, 10% glycerol, 0.34 M Sucrose, 1 mM DTT, 0.1% Triton, PMSF, and protease inhibitor mixture) for 10 min at 4 °C. The cell lysates were centrifuged at 1,500 × g for 5 min at 4 °C, with supernatants removed. Pellets were resuspended in buffer B (3 mM EDTA, 0.2 mM EGTA, 1 mM DTT, PMSF, and protease inhibitor mixture) and incubated on ice for 10 min before centrifugation at 1,700 × g for 5 min at 4 °C. To prepare the chromatin-bound fraction, the pellets were incubated for 30 min at 37 °C with buffer A containing 1 mM CaCl2 and 0.6 unit of MNase and centrifuged at 20,000 × g for 10 min at 4 °C, and the supernatant was collected and subjected to immunoblot analysis.

### Quantitative RT-PCR

Quantitative RT–PCR (qRT–PCR) was performed after RNA extraction using TRIzol (Thermo Fisher). RNA expression was determined with Universal SYBR Green Two-step kit (Yeasen) on a QuantStudio 7 Real-Time PCR System (Thermo). For normalization of human gene expression, *RPL13A* was used as an internal control, and the fold change was calculated using the 2-ΔΔC(t) method for all analyses (all primer sequences listed in Supplementary Table 4).

### Immunoblotting and immunofluorescence

Proteins were separated using NuPAGE 4 to 12% Bis-Tris gels and transferred onto nitrocellulose membranes (Life Technologies). The blots were blocked with 5% nonfat dry milk at room temperature for 1 h and incubated overnight at 4°C with desired primary antibodies at concentration per the manufacturer’s protocol, followed by incubation with horseradish peroxidase-conjugated secondary antibodies (Santa Cruz) at room temperature for 1 h. The membrane blots were developed with enhanced chemiluminescence (ECL) detection reagent (Millipore) per the manufacturer’s protocol and detected through chemiluminescence using ImageQuant LAS 400 Phospho-Imager (GE Healthcare). As a standard protein marker, we used Thermo Fisher Scientific PageRuler Plus Prestained Protein Ladder (no. 26619).

For immunofluorescence staining, cells were cultured in dishes and pre-seeded for at least 24 h on coverslips. Upon brief washing, cells were fixed with 4% paraformaldehyde in PBS for 8 min, blocked with 5% normal goat serum (NGS, Thermo Fisher) for 30 min and incubated with primary antibodies diluted in PBS containing 5% NGS for 2 h at room temperature. Alexa Fluor 488, 555 or 595-conjugated secondary antibodies (Invitrogen, 1:400) were used. Nuclei were stained with Hoechst 33342, with slides mounted with Vectashield medium. Fluorescence imaging was performed on a fluorescence microscope (Nikon Eclipse Ti S). Captured images were analyzed and processed with the Nikon DS-Ri2 fluorescence workstation and processed with NIS-Elements F4.30.01. Alternatively, a confocal microscope (Zeiss LSM 780) was applied to acquire confocal images (antibodies used for this study listed in Key Resources Table).

### SILAC sample preparation

PSC27 cells were grown to 80% confluence in high glucose (4.5 g/L) DMEM media (with glutamine and sodium pyruvate) containing 10% FBS and 1% penicillin/streptomycin at 37 °C with 95% air and 5% CO2. Cells were either labeled with “heavy isotopic lysine” (^13^C-lysine) or “light isotopic lysine” (^12^C-lysine/arginine) using a SILAC protein quantitation kit (Pierce, Thermo) according to manufacturer’s instructions. Briefly, cells in the control group were labeled with L-^13^C6-Lysine/L-^13^C6^15^N4-Arginine as “heavy” class, while those in the experimental group were labeled with L-Lysine/L-Arginine as “light” class. In each case, cells were cultured for more than 6 generations before being harvested. At least 1 passage prior to the experiment, incorporation efficiency of the heavy labeled amino acids into proteins was assessed in a pilot experiment, where a small aliquot of intracellular protein was lysed, reduced, alkylated, and trypsin digested and subjected to mass spectrometric (MS) analysis as described below. Heavy label incorporation into proteins from control (non-senescent) cells was assessed as > 99%.

For the proteomics, 24 h before harvesting, plated cells (approximately 5 × 10^8^ in fifteen 150 cm2 flasks) were washed thrice with fresh FBS-free medium for 10 min at 37°C each. The cells were then incubated in FBS-free medium (12 ml per flask) for 24 h at 37°C. At the end of the culture period, cells showed no evidence of apoptosis. Samples were prepared for MS analysis including removal of phenol red dye by buffer exchange with 50 mM Tris-Cl (pH8.0) (5 washing steps using spin columns at 5000 × g). Samples were denatured with 6M urea, reduced with 20mM DTT (30 min at 37°C), alkylated with 50 mM iodoacetamide (30 min at RT), and digested overnight at 37°C with 1:50 enzyme:substrate ratio (wt/wt) of sequencing grade trypsin (Promega, Madison, WI) as described ^59^. Following digestion, samples were acidified with formic acid and desalted using HLB Oasis SPE cartridges (Waters, Milford, MA). Protein/peptide recovery was not specifically assessed, however, the above sample preparation and digestion protocol was thoroughly assessed for reproducibility during previous optimization and reproducibility studies. Hydrophilic Interaction Liquid Chromatography (HILIC) peptide fractionation was performed on a Waters 1525 HPLC system equipped with a 4.6 × 25 mm TSK gel Amide-80 HR 5 mm column (Tosoh Bioscience, South San Francisco, CA). Samples were loaded in 80% solvent B (98% acetonitrile, 0.1% TFA) and eluted with the following gradient: 80% B for 5 min followed by 80% B to 60% B in 40 min, 0% B in 5 min at 0.5 ml/min. Solvent A consisted of 98% HPLC grade water and 0.1% TFA collecting nine fractions.

### Mass spectrometry

Samples were analyzed by reverse-phase HPLC-ESI-MS/MS using an Eksigent Ultra Plus nano-LC 2D HPLC system (Dublin, CA) connected to a quadrupole time-of-flight TripleTOF 5600 (QqTOF) mass spectrometer (SCIEX). Typically, mass resolution for MS1 scans and corresponding precursor ions was ∼35,000 (TripleTOF 5600), while resolution for MS/MS scans and resulting fragmentions (MRM-HR transitions) was ∼15,000 (‘high sensitivity’ product ion scan mode) ^60^. Briefly, after injection, peptide mixtures were transferred onto the analytical C18-nanocapillary HPLC column (C18 Acclaim PepMap100, 75 mm I.D. × 15 cm, 3 mm particle size, 100A° pore size, Dionex, Sunnyvale, CA) and eluted at a flow rate of 300 nl/min using stepwise gradients from 5% to 80% solvent B with total runtimes, including mobile phase equilibration, of 90 min. Solvent mobile phase A was 2% acetonitrile/98% of 0.1% formic acid (v/v) in water, and mobile phase B was 98% acetonitrile/2% of 0.1% formic acid (v/v) in water. Data acquisition was performed in data dependent acquisition (DDA) mode on the TripleTOF 5600 to obtain MS/MS spectra for the 30 most abundant precursor ions (50 msec per MS/MS) following each survey MS1 scan (250 msec), yielding a total cycle time of 1.8 s as described ^60, 61^.

### Stromal cell clonogenic assay

PSC27 cells were trypsinized and counted with a hemocytometer. Two milliliters of DMEM full medium containing 500 cells were plated in each well of the six-well plates. The cells were maintained at 37 °C for 7 d to allow colony formation before stained with 0.5% crystal violet (Sigma-Aldrich) in absolute methanol. Colonies per well were counted and numbers were recorded from 3 independent experiments.

### Cancer cell phenotype appraisal

Epithelial cell proliferation was determined following the MTT procedure (Promega). Migration and invasion were assessed in culture using Transwells (Cultrex 24-well Cell Migration Assay plates) containing a porous (8 μm pore size) membrane uncoated (for the migration assay) or coated (for the invasion assay) with a 0.5 × solution of basement membrane extract and the indicated CM in the bottom portion of the well. After 24 h, migrating or invading cells on the bottom side of the porous membranes were stained and quantified by absorbance as recommended by the supplier.

### Histology and immunohistochemistry

Mouse tissue specimens were fixed overnight in 10% neutral-buffered formalin and processed for paraffin embedding. Standard staining with hematoxylin/eosin was performed on sections of 5-8 μm thickness processed from each specimen block. For immunohistochemistry, tissue sections were de-paraffinized and incubated in citrate buffer at 95 °C for 40 min for antigen retrieval before incubated with the indicated antibodies overnight at 4 °C. After 3 washes with PBS, tissue sections were incubated with biotinylated anti-mouse IgG (1:200 dilution, Vector Laboratories) for 1 h at room temperature then washed thrice, after which streptavidin-horseradish peroxidase conjugates (Vector Laboratories, CA, USA) were added and the slides incubated for 45 min. DAB solution (Vector Laboratories) was then added and the slides were counterstained with haematoxylin.

### Bioinformatics database searches and SILAC quantification

MS data were searched using the database search engine ProteinPilot (SCIEX Beta 4.5, revision 1656) with the Paragon algorithm (4.5.0.0, 1654). The search parameters were set as follows: trypsin digestion, cysteine alkylation set to iodoacetamide, SILAC Quantification (Lys-6, Arg-6, no bias correction), and Homo sapiens as species. Trypsin specificity was assumed as C-terminal cleavage at lysine and arginine. Processing parameters were set to ‘‘Biological modification’’ and a thorough ID search effort was used. During the search, Protein Pilot performs an automatic mass recalibration of the datasets based on highly confident peptide spectra. Specifically, a first search iteration was done to select high confidence peptide identifications to recalibrate both the MS and MS/MS data, which was subsequently automatically re-searched. During the iterative steps of re-searching the data, the search parameters were less stringent, e.g., allowing for additional ‘missed cleavages’ (with typically not more than 2).

All data files were searched using the SwissProt (2014-01, released January 22, 2014), with a total of 40,464 human ‘reviewed’ protein sequences searched. A cut-off peptide confidence value of 99 was chosen, and a minimum of 2 identified peptides per protein was required. The Protein Pilot false discovery rate (FDR) analysis tool, the Proteomics System Performance Evaluation Pipeline (PSPEP algorithm) provided a global FDR of 1% and a local FDR at 1% in all cases. For protein quantification comparing proteins from 3 biological replicate experiments between CM from senescent (light) *versus* presenescent (heavy) cells a Protein Pilot significance threshold of < 0.05 was required (all database search results and details for peptide identifications and protein quantification are provided in Supplementary Tables S1-S3).

### RNA-seq and data profiling

Total RNA samples were obtained from stromal cells. Sample quality was validated by Bioanalyzer 2100 (Agilent), and RNA was subjected to sequencing by Illumina HiSeq X10 with gene expression levels quantified by the software package RSEM (https://deweylab.github.io/RSEM/). Briefly, rRNAs in the RNA samples were eliminated using the RiboMinus Eukaryote kit (Qiagen, Valencia, CA, USA), and strand-specific RNA-seq libraries were constructed using the TruSeq Stranded Total RNA preparation kits (Illumina, San Diego, CA, USA) according to the manufacturer’s instructions before deep sequencing.

Paired-end transcriptomic reads were mapped to the reference genome (GRCh38/hg38) with reference annotation from Gencode v27 using the Bowtie tool. Duplicate reads were identified using the picard tools (1.98) script mark duplicates (https://github.com/broadinstitute/picard) and only non-duplicate reads were retained. Reference splice junctions weree provided by a reference transcriptome (Ensembl build 73). FPKM values were calculated using Cufflinks, with differential gene expression called by the Cuffdiff maximum-likelihood estimate function. Genes of significantly changed expression were defined by a false discovery rate (FDR)-corrected *P* value < 0.05. Only ensembl genes 73 of status “known” and biotype “coding” were used for downstream analysis.

Reads were trimmed using Trim Galore (v0.3.0) (http://www.bioinformatics.babraham.ac.uk/projects/trim_galore/) and quality assessed using FastQC (v0.10.0) (http://www.bioinformatics.bbsrc.ac.uk/projects/fastqc/). Differentially expressed genes were subsequently analyzed for enrichment of biological themes using the DAVID bioinformatics platform (https://david.ncifcrf.gov/), the Ingenuity Pathways Analysis (IPA) program (http://www.ingenuity.com/index.html). Raw data were preliminarily analyzed on the free online platform of Majorbio I-Sanger Cloud Platform (www.i-sanger.com), and subsequently deposited in the NCBI Gene Expression Omnibus (GEO) database under the accession code GSE128282.

#### Venn diagrams

Venn diagrams and associated empirical *P*-values were generated using the USeq (v7.1.2) tool IntersectLists ^3^. The t-value used was 22,008, as the total number of genes of status “known” and biotype “coding” in ensembl genes 73. The number of iterations used was 1,000.

#### RNA-seq heatmaps

For each gene, the FPKM value was calculated based on aligned reads, using Cufflinks. Z-scores were generated from FPKMs. Hierarchical clustering was performed using the R package heatmap.2 and the distfun = “pearson” and hclustfun = “average”.

#### Protein–protein interaction network

Protein-protein interaction (PPI) analysis was performed with STRING 3.0. The specific proteins meeting the criteria, were imported to NetworkAnalyst (http://www.networkanalyst.ca). A minimum interaction network was chosen for further hub and module analysis.

### Gene set enrichment analysis (GSEA) of RNA-seq data

For each differential expression analysis comparison, genes were ranked using “wald statistics” obtained from DESeq2 and GSEA was performed on these ranked lists on all curated gene sets available in MSigDB (http://software.broadinstitute.org/gsea/msigdb). DESeq2 independent filtering is based on mean of normalized read counts and filters out genes with very low expression level. The SASP and GSEA signatures were derived as described before ^40^.

### Omni-ATAC protocol

Cells for each of CTRL, BLEO or BLEO/ML324 group (3 replicates per group) were pretreated with 200 U/ml DNase (Worthington) for 30 min at 37 °C to remove free-floating DNA and to digest DNA of dead cells. The media were then washed out, cells were resuspended in cold PBS for counting and processed as previously described ^62^. Briefly, a number of 50,000 cells was resuspended in 1 ml of cold ATAC-seq resuspension buffer (RSB; 10 mM Tris-HCl pH 7.4, 10 mM NaCl, and 3 mM MgCl2 in water). Cells were centrifuged at 500 r.c.f. for 5 min in a pre-chilled (4 °C) fixed-angle centrifuge. After centrifugation, 900 μl of supernatant was aspirated, which left 100 μl of supernatant. This remaining 100 μl of supernatant was carefully aspirated by pipetting with a P200 pipette tip to avoid the cell pellet. Cell pellets were then resuspended in 50 μl of ATAC-seq RSB containing 0.1% NP40, 0.1% Tween-20, and 0.01% digitonin by pipetting up and down thrice. This cell lysis reaction was incubated on ice for 3 min. After lysis, 1 ml of ATAC-seq RSB containing 0.1% Tween-20 (without NP40 or digitonin) was added, and the tubes were inverted to mix. Nuclei were then centrifuged for 10 min at 500 r.c.f. in a pre-chilled (4 °C) fixed-angle centrifuge. Supernatant was removed with two pipetting steps, as described before, and nuclei were resuspended in 50 μl of transposition mix (25 μl 2 × TD buffer, 2.5 μl transposase (100 nM final), 16.5 μl PBS, 0.5 μl 1% digitonin, 0.5 μl 10% Tween-20, and 5 μl H2O) by pipetting up and down six times. The remainder of the ATAC-seq library preparation was performed by following manufacturer’s instructions (Nextera DNA Flex Library Prep kit, Illumina, cat. no. 20018704) and the sample quality was validated by BioAnalyzer 2100 (Agilent) before proceeding to deep sequencing. Briefly, all libraries were amplified with a target concentration of 20 μl at 4 nM, which is equivalent to 80 femtomoles of product. Libraries were purified with the 1.5 × AMPure (Beckman) beads and were subjected to next-generation sequencing. Raw data were deposited in the NCBI Gene Expression Omnibus (GEO) database under the accession code GSE135481.

### ATAC-seq data processing

The single-end ATAC-seq reads were aligned to hg19 reference genome with random chromosome cleaned by Bowtie (version 2.2.2) ^63^ under the parameters -t -q -N 1 -L 25. The paired-end ATAC-seq reads were aligned with the parameters: -t -q -N 1 -L 25 -X 2000 -no-mixed–no-discordant. All unmapped reads, non-uniquely mapped reads and PCR duplicates were removed. For downstream analysis, we normalized the read counts by computing the numbers of reads per kilobase of bin per million of reads sequenced (RPKM). RPKM values were averaged for each bin between replicates. To minimize the batch and cell type variation, the RPKM values were further normalized by *Z*-score transformation. To visualize the ATAC-seq signal in the UCSC genome browser, we extended each read by 250 base pairs (bp) and counted the coverage for each base. The correlation between ATAC-seq replicates was calculated as following: each read was extended 250 bp from the mapped end position and the RPKM value was generated on a 100 bp-window base. The ATAC-seq enrichment was then summed within each 2-kb window for the entire genome and was compared between replicates. Pearson correlation was calculated and was shown. In brief, to assign each read to its parental origins, we examined all SNPs in the read that showed high-quality base calling (Phred score ≥ 30). For paired-end reads, SNP information from both reads in the pair was summed and used. When multiple SNPs were present in a read (or a read pair), the parental origin was determined by votes from all SNPs and the read was assigned to the allele that had at least two thirds of the total votes.

### Identification of ATAC-seq peaks and their genome coverages

All the ATAC-seq peaks were called by MACS v1.4 ^64^ with the parameters-nolambda-nomodel. ATAC-seq peaks at least 2.5 kb away from annotated promoters from ref-Flat were selected as distal ATAC-seq peaks. The genome coverages of peaks from different samples were calculated by genomeCoverageBed ^65^ using hg19 reference genome.

### Comparison between ATAC-seq peaks and known *cis*-regulatory elements

To compare the ATAC-seq peaks identified in senescent cells with the annotated *cis*-regulatory elements, we calculated the overlap between the ATAC-seq peaks of different states and annotated promoters (TSS ± 0.5 kb). Distal peaks were then compared to distal DNase I hypersensitive sites in stromal cells. Random peaks were generated by selecting random regions in the genome with the sizes matching each individual ATAC-seq peak.

### Comparison between ATAC-seq peaks and repetitive elements

To identify the overlap between repetitive elements and promoter or distal ATAC-seq peaks, the ATAC-seq peaks were compared with the locations of annotated repeats (RepeatMasker) downloaded from the UCSC genome browser by intersectBed ^65^ with default parameters. As repeats of different classes vary greatly in numbers, a random set of peaks with identical lengths of ATAC-seq peaks was used for the same analysis as a control. The numbers of observed peaks that overlap with repeats were compared to the numbers of random peaks that overlap with repeats, and a log ratio value (log2) was generated as the ‘observed/expected’ enrichment.

### Prediction of promoter targets of putative enhancers in distal ATAC-seq peaks

To identify the potential targeted genes for stage specific enhancer (distal peaks), we computed the averaged ATAC-seq enrichment (normalized RPKM) for all distal ATAC-seq peaks and annotated promoters (TSS ± 0.5 kb). Among genes assigned to enhancers by GREAT analysis based on distances, we further calculated the correlation between the ATAC-seq enrichment at distal ATAC peaks and these promoters across human stromal cells. The promoter with a Pearson correlation coefficient above 0.8 was selected as the potential target of the enhancer.

### Gene ontology analysis

The DAVID web-tool was used to identify the Gene Ontology (GO) terms using databases including Molecular Functions, Biological Functions and Cellular Components ^66^. Because most lists of GO terms are large and redundant, REVIGO was applied to summarize redundant GO terms based on semantic similarity measures ^67^. Nonredundant GO term sets were visualized by Cytoscape, in which GO terms were set as nodes and 3% of GO term pairwise similarities were set as edges.

### Hierarchical clustering analysis

The hierarchical clustering was performed in R by hclust() function with ATAC-seq RPKM values via Spearman correlation coefficients.

### Motif, enhancer mark and transcription factor-binding sequence analyses for distal ATAC-seq peaks

To find the sequence motifs enriched in distal ATAC-seq peaks, findMotifsGenome.pl from the HOMER program ^68^ was used. Motifs with known match in HOMER database were selected. The ChIP–seq data for human stromal cell H3K9me3 and H3K36me3 marks ^69^ and the collection of transcription factor (TF)-binding sites ^70^ were downloaded from the UCSC genome browser. The average RPKM values at the distal peaks and their nearby regions were calculated. The number of transcription factor-binding sites was first binned for each 100-bp window, and the average enrichment at the distal peaks and their nearby regions was calculated.

To connect TFs to genes at a particular cell state, we first identified TF motifs present in distal ATAC-seq peaks and selected those highly enriched (*P* value < 1 × 10−10). We then assigned these TF motifs to genes by the distal-promoter peak pairs established in this work. If multiple putative enhancers were assigned to the same promoter, the numbers of TF motifs within these enhancers assigned to a gene were then summed. If a gene receives no assignment of motifs for a TF from its linked enhancers, the number of TF motif for this gene is 0.

### Functional analysis, protein ontology, pathway enrichment analysis

For functional analysis and protein ontology analysis, ‘The Database for Annotation, Visualization and Integrated Discovery’ (DAVID v.6.8) was used ^71, 72^. For pathway enrichment analysis, we used the Benjamini correction as adjustment to P values to determine statistical significance.

### Genomics mapping of human KDM4 gene alterations across TCGA datasets

The TCGA multi-omics data were collected and subject to clinical characterization for human prostate and breast cancer patients at cBioPortal, an open-access resource for interactive exploration of multidimensional cancer genomics data sets ^73^. A propensity score matching (PSM) analysis was applied to identify significant molecular features, as proposed formerly ^74^.

Unlike pair matching, this method improves balance and estimates efficiency and employs all subjects by weighting them so that every subject potentially contributes to the estimation ^75^.

### Generation of density heatmaps and profiles

For heatmaps and profiles, we used deepTools (v2.5.0) ^76^ to generate read abundance from all datasets around peak center (± 2.5 kb/ 2.0 kb), using ‘computeMatrix’. These matrices were then used to create heatmaps and profiles, using deep-Tools commands ‘plotHeatmap’ or ‘plotProfile’ respectively.

### Preclinical trials with experimental animals

Nod-obese diabetic and severe combined immune-deficiency (NOD-SCID) male mice (NanJing Model Animal Centry, China) of 6-8 weeks old were housed and maintained in accordance with animal guidelines of Shanghai Institutes for Biological Sciences. For mouse tumor xenograft establishment, human prostate stromal cells (PSC27) and cancer epithelial cells (PC3 or PC3-Luc) were mixed at a ratio of 1:4, with each *in vitro* recombinant comprising 1.25 × 10^6^ total cells prior to subcutaneous implantation. Two weeks later, mice were randomized into groups and subject to preclinical treatments. Animals were treated by MIT (0.2 mg/kg) alone or MIT plus ML324 (4 mg/kg). Agents were delivered intraperitoneally (i.p.) once *per* biweekly starting from the beginning of the 3rd week, with totally 3 biweek cycles throughout the whole regimen.

Mice were sacrificed at end of an 8^th^ week post tumor xenograft regimen. Primary tumor sizes were measured upon animal dissection, with approximate ellipsoid tumor volume (V) measured and calculated according to the tumor length (l) and width (w) by the formula: V = (π/6) × ((l + w)/2)^3^ used previously ^40^. Excised tumors were either freshly snap-frozen, or fixed in 10% buffered formalin and subsequently processed as formalin-fixed and paraffin-embedded sections (FFPE) for IHC staining. Tumor growth and metastasis in mice was evaluated using the bioluminescence emitted by PC3-luc cells which stably express the firefly luciferase. The Xenogen IVIS Imager (Caliper Lifesciences) was applied to document BLI across the visible spectrum, with the substrate D-Luciferin (150 mg/kg, BioVision) injected subcutaneously each time for real time tumor surveillance.

### *In vivo* cytotoxicity assessment *via* blood test

As routine blood examination, 100 μl fresh blood was acquired from each animal and mixed with EDTA immediately. The blood samples were analyzed by Celltac Alpha MEK-6400 series hematology analyzers (Nihon Kohden, MEK-6400). For serum biochemical analysis, blood samples were collected, clotted for 2 h at room temperature or overnight at 4 °C. Samples were then centrifuged (1000 × g, 10 min) to obtain serum. An aliquot of approximately 50 μl serum was subject to analysis for creatinine, urea, alkaline phosphatase (ALP) and alanine transaminase (ALT) by Chemistry Analyzer (Mindray, BS-350E). Evaluation of circulating levels of hemoglobin, white blood cells, lymphocytes and platelets were performed with dry-slide technology on VetTest 8008 chemistry analyzer (IDEXX) as reported previously ^31^.

### Study approval

All animal experiments were conducted in compliance with NIH Guide for the Care and Use of Laboratory Animals (National Academies Press, 2011) and the ARRIVE guidelines, and were approved by the Institutional Animal Care and Use Committee (IACUC) of the University of Washington or Shanghai Institutes for Biological Sciences, Chinese Academy of Sciences. Mice were maintained under specific pathogen-free (SPF) conditions, with food and water provided ad libitum.

### Statistical methods

Unless otherwise specified, data in the figures are presented as mean ± SD, and statistical significance was determined by unpaired two-tailed Student’s *t*-test (in the case of comparing two groups), one-way ANOVA or two-way ANOVA (comparing more than two groups), Pearson’s correlation coefficients test, Kruskal–Wallis, log-rank test, Wilcoxon–Mann–Whitney test, or Fisher’s exact test with GraphPad Prism 8.0 primed with customized parameters. Cox proportional hazards regression model and multivariate Cox proportional hazards model analysis were performed with statistical software SPSS.

To determine sample size, we chose to set the values of type I error (α) and power (1−β) to be statistically adequate: 0.05 and 0.80, respectively ^77^. We then determined n on the basis of the smallest effect we wish to measure. If the required sample size is too large, we chose to reassess the objectives or to more tightly control the experimental conditions to reduce the variance. For all statistical tests, a p value < 0.05 was considered significant, with p values presented as follows throughout the article: ^, not significant; *, *P* < 0.05; **, *P* < 0.01; ***, *P* < 0.001. For dose response curve, p values were calculated by non-linear regression with extra sum-of-squares F test. The usage of all statistical approaches was examined by biostatistics experts. All bioinformatics analysis and comparisons are described in details below.

### Data availability

Requests for further information and reagents should be directed to and will be fulfilled by the corresponding author. Any plasmid, cell line and associated reagent generated in this study would be available upon reasonable request, and with a completed Materials Transfer Agreement if there is potential for commercial application. The RNA-seq data have been deposited in GEO with an accession number GSE128282. ATAC-seq data have been deposited in GEO with an accession number GSE135481.

## Acknowledgments

This work was supported by grants from National Key Research and Development Program of China (2016YFC1302400, 2020YFC2002801), National Natural Science Foundation of China (NSFC) (81472709, 31671425, 31871380) to Y.S., the Strategic Priority Research Program of Chinese Academy of Sciences (XDB39010500) to Y.S., the fund of Key Laboratory of Tissue Microenvironment and Tumor of Chinese Academy of Sciences (201506, 201706) to Y.S.; Anti-Aging Collaborative Program of SIBS and BY-HEALTH (C01201911260006) to Y.S., the University and Locality Collaborative Development Program of Yantai (2019XDRHXMRC08) Y.S., and the U.S. DoD PCRP (Idea Development Award PC111703) to Y.S.; U.S. NIH grants R37-AG013925 and P01 AG062413, the Connor Fund, Robert J. and Theresa W. Ryan, and the Noaber Foundation to J.L.K.; Breast Cancer Now (2012MayPR070; 2012NovPhD016), the Medical Research Council of the United Kingdom (MR/N012097/1), Cancer Research UK Imperial Centre, Imperial ECMC and NIHR Imperial BRC to E.W-F.L.; National Natural Science Foundation of China (NSFC) (31870927, 81671552) to X.C.; and National Key Research and Development Program of China (2020YFC2002800) to X.Z.

## Author contributions

Y.S. and B.Z. conceived this study, designed the experiments and supervised the project. B.Z. performed most of the biological experiments. Q.L. acquired and analyzed clinical samples from prostate cancer patients, and managed subject information. L.H. and Y.S. performed data mining and bioinformatics of gene expression and signaling pathways. S.W., S.S., Q.X., L.H., M.Q., X.R., J.J., and Q.F. helped *in vitro* culture and phenotypic characterization of cancer cells. J.G., X.Z., X.C., E.W-F.L., J.C. and J.L.K. provided conceptual inputs or supervised a specific subset of experiments. Q.L. and Y.S. performed preclinical studies. Y.S. orchestrated data integration and prepared the manuscript. All authors critically read and commented on the final manuscript.

## Additional information

**Extended data** is available for this paper online.

**Supplementary information** is available for this paper online.

**Correspondence and requests for materials** should be addressed to Y. Sun.

