## Supplementary Information for "KDM4 Orchestrates Epigenomic Remodeling of Senescent Cells and Potentiates the Senescence-Associated Secretory Phenotype"

### Extended Data Fig. 1

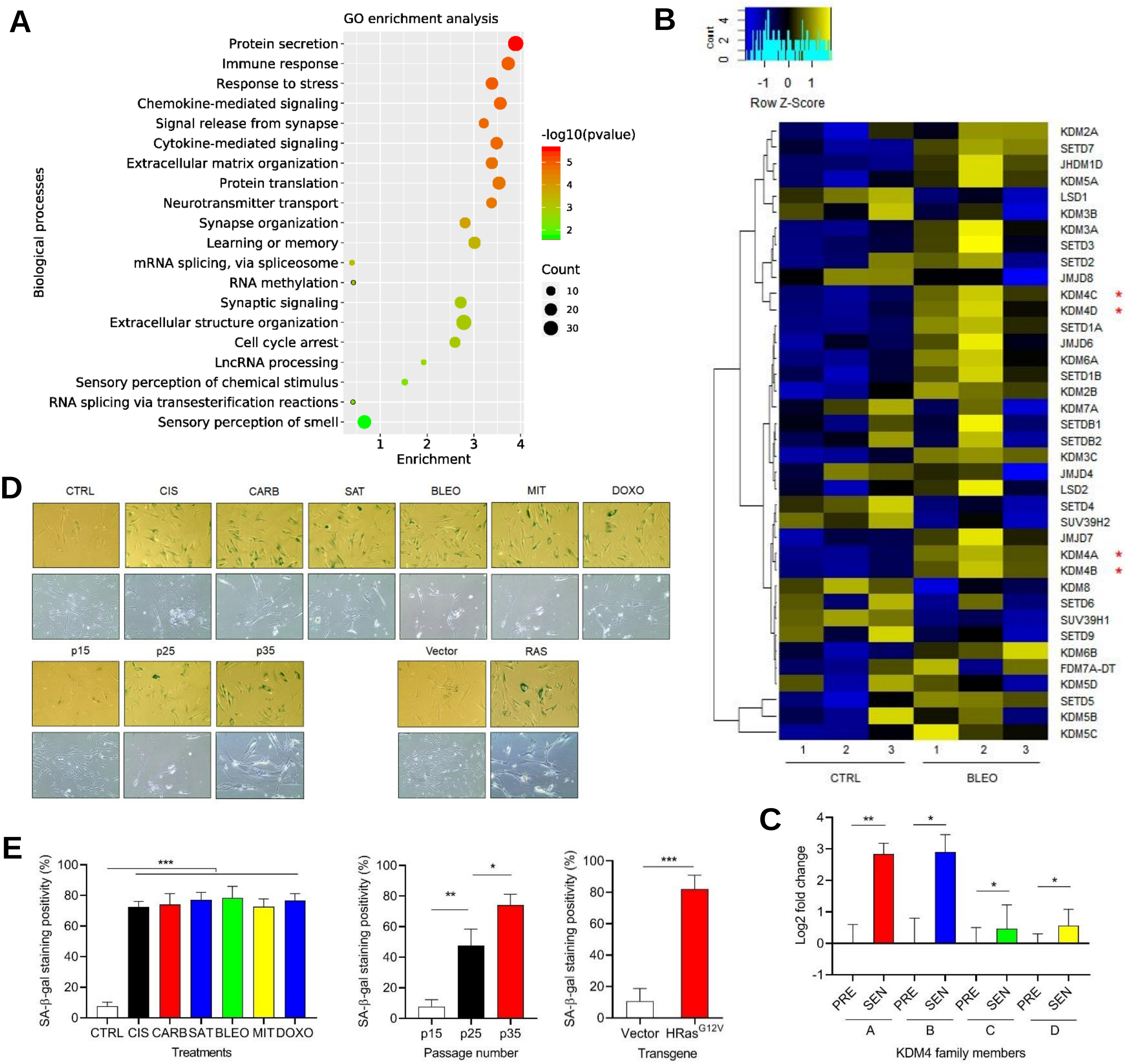

Extended Data Fig. 2

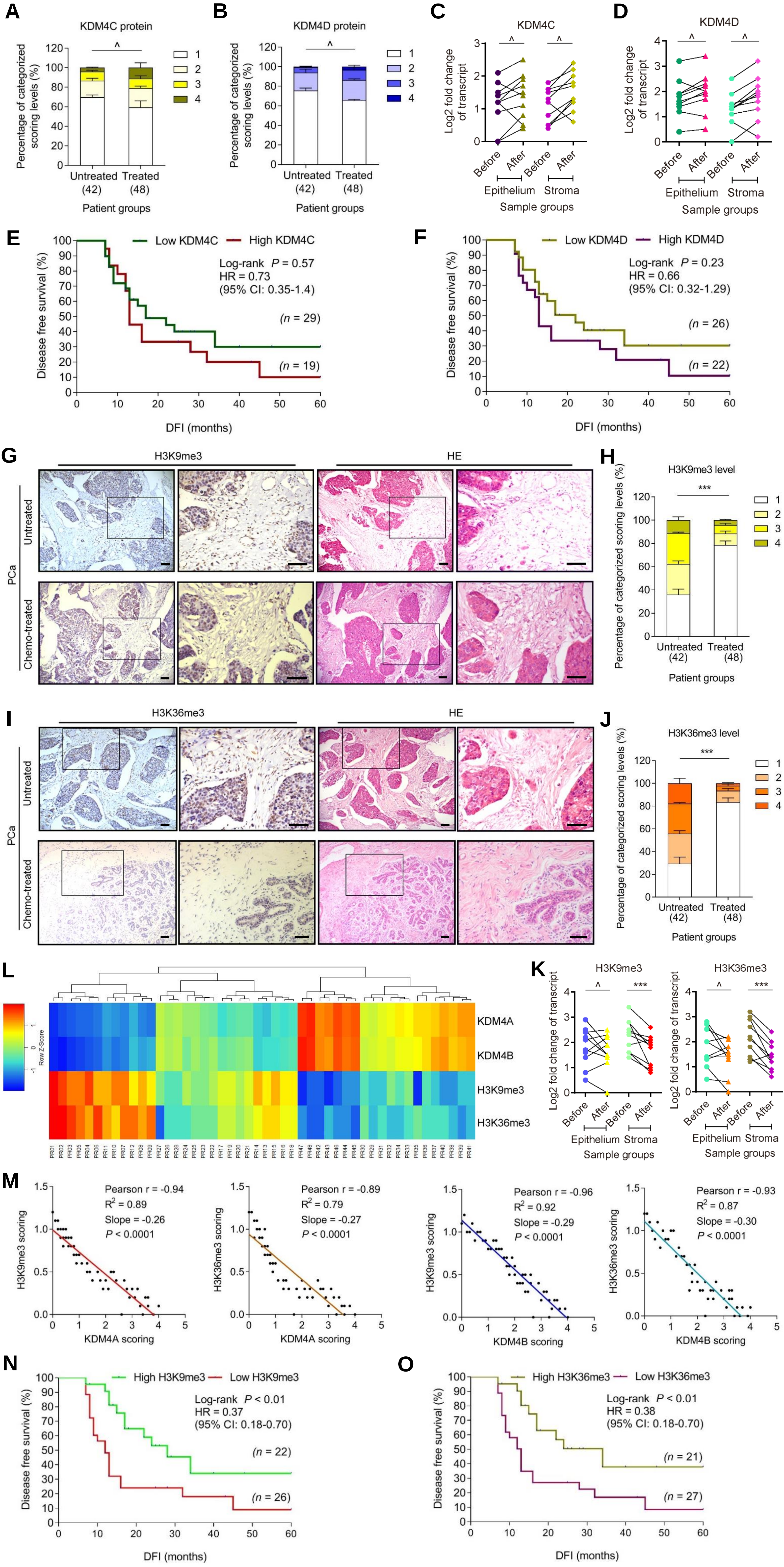

### Extended Data Fig. 3

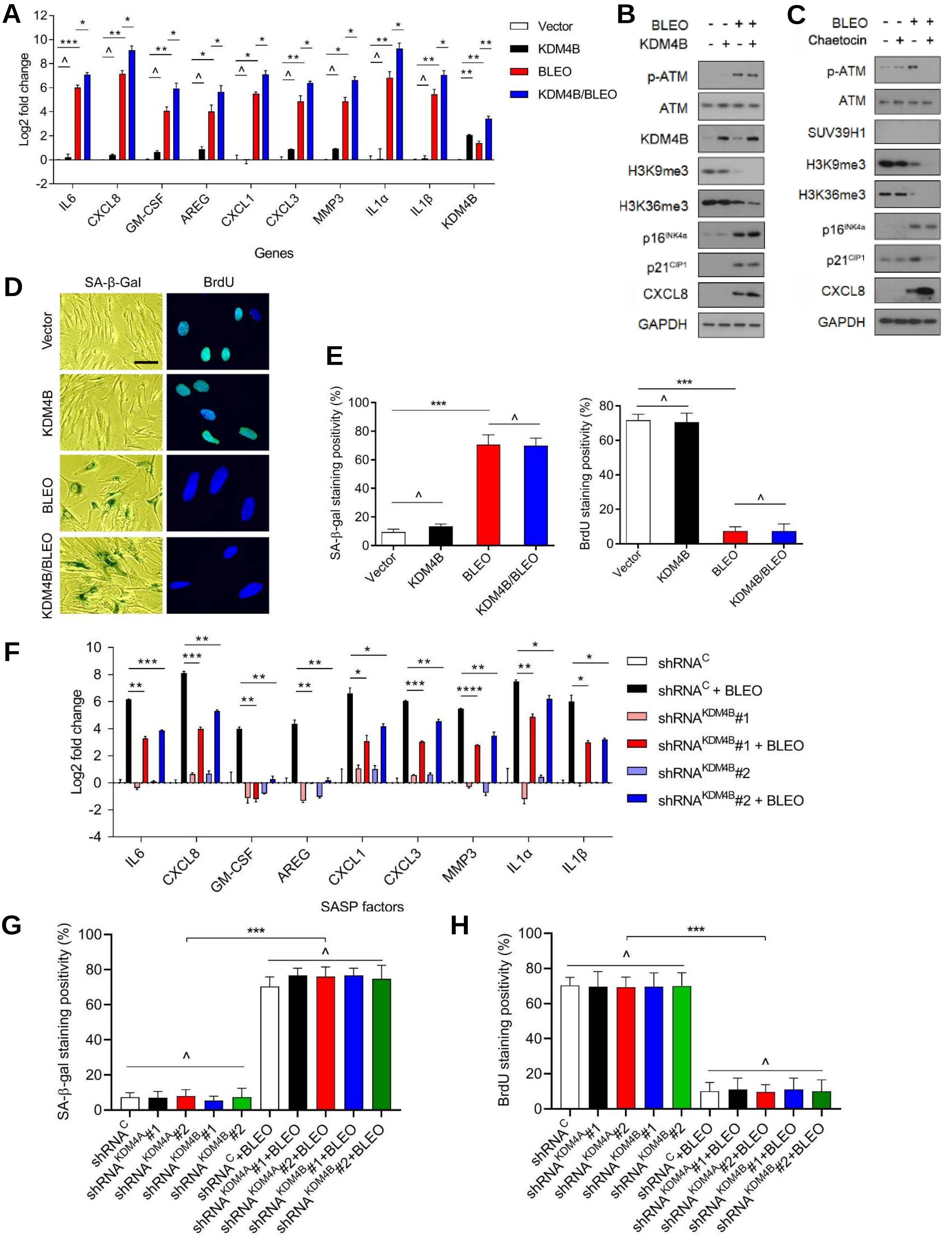

### Extended Data Fig. 4

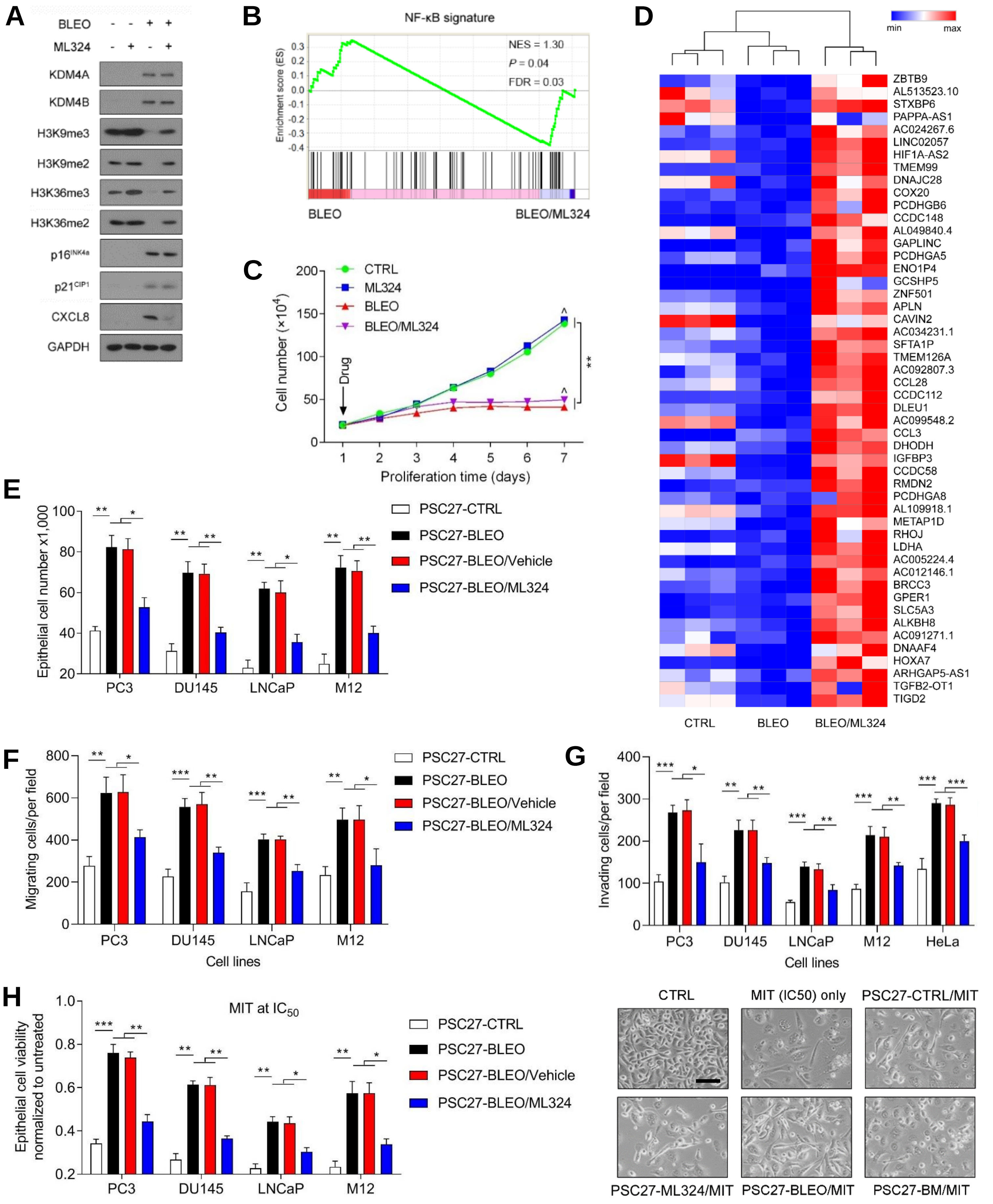

**Extended Data Fig. 5**

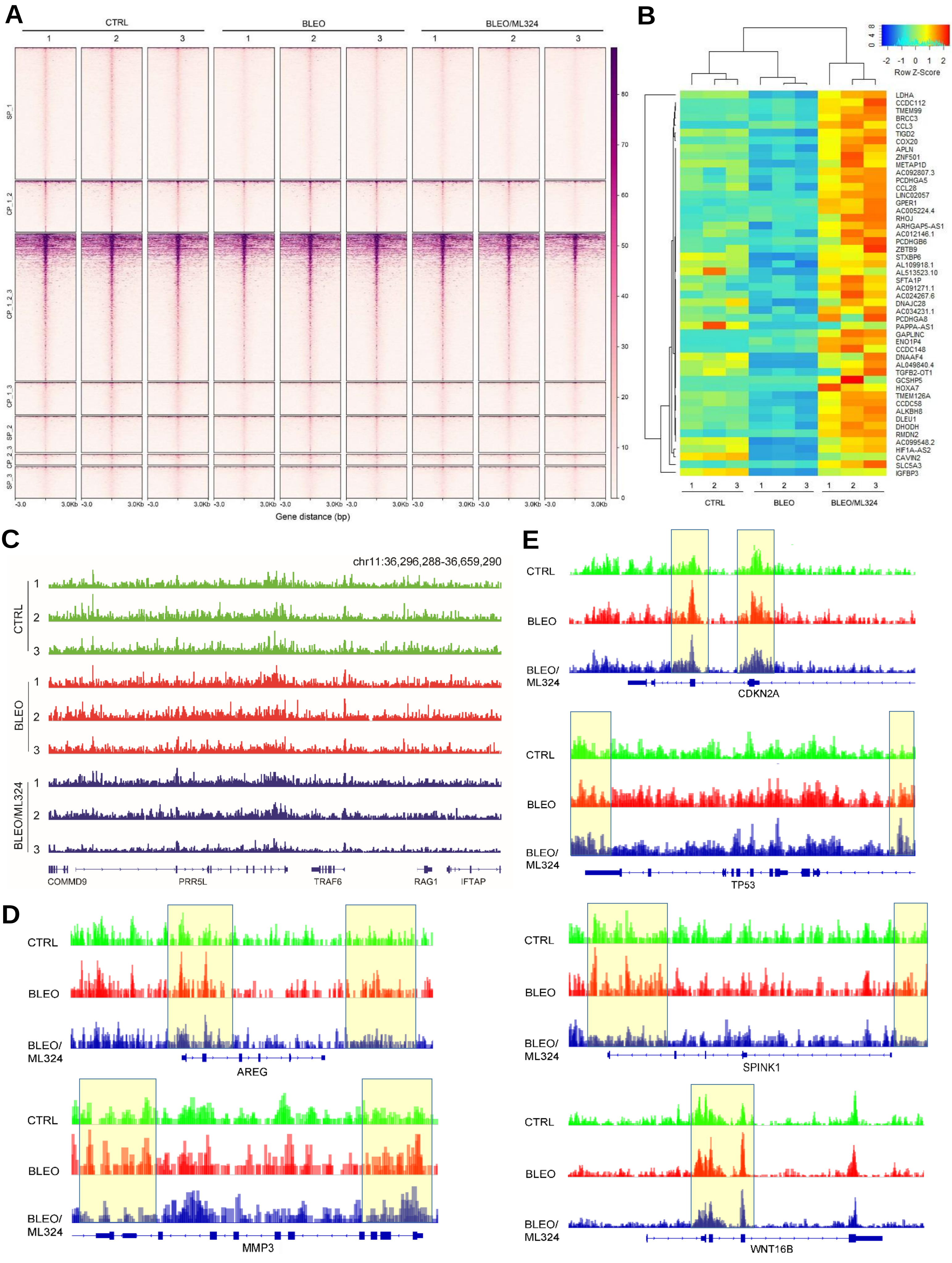

Extended Data Fig. 6

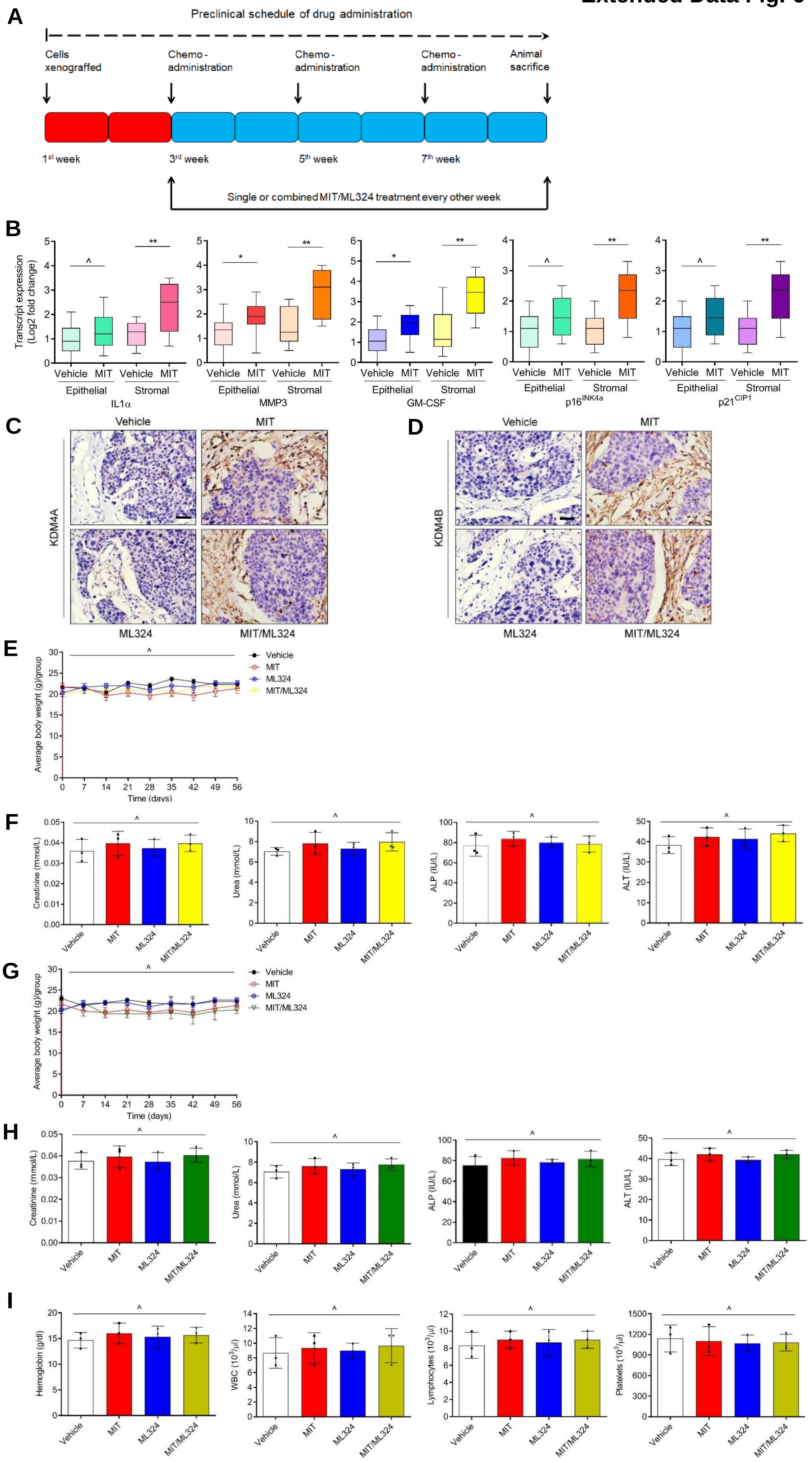

### Extended Data Fig. 7

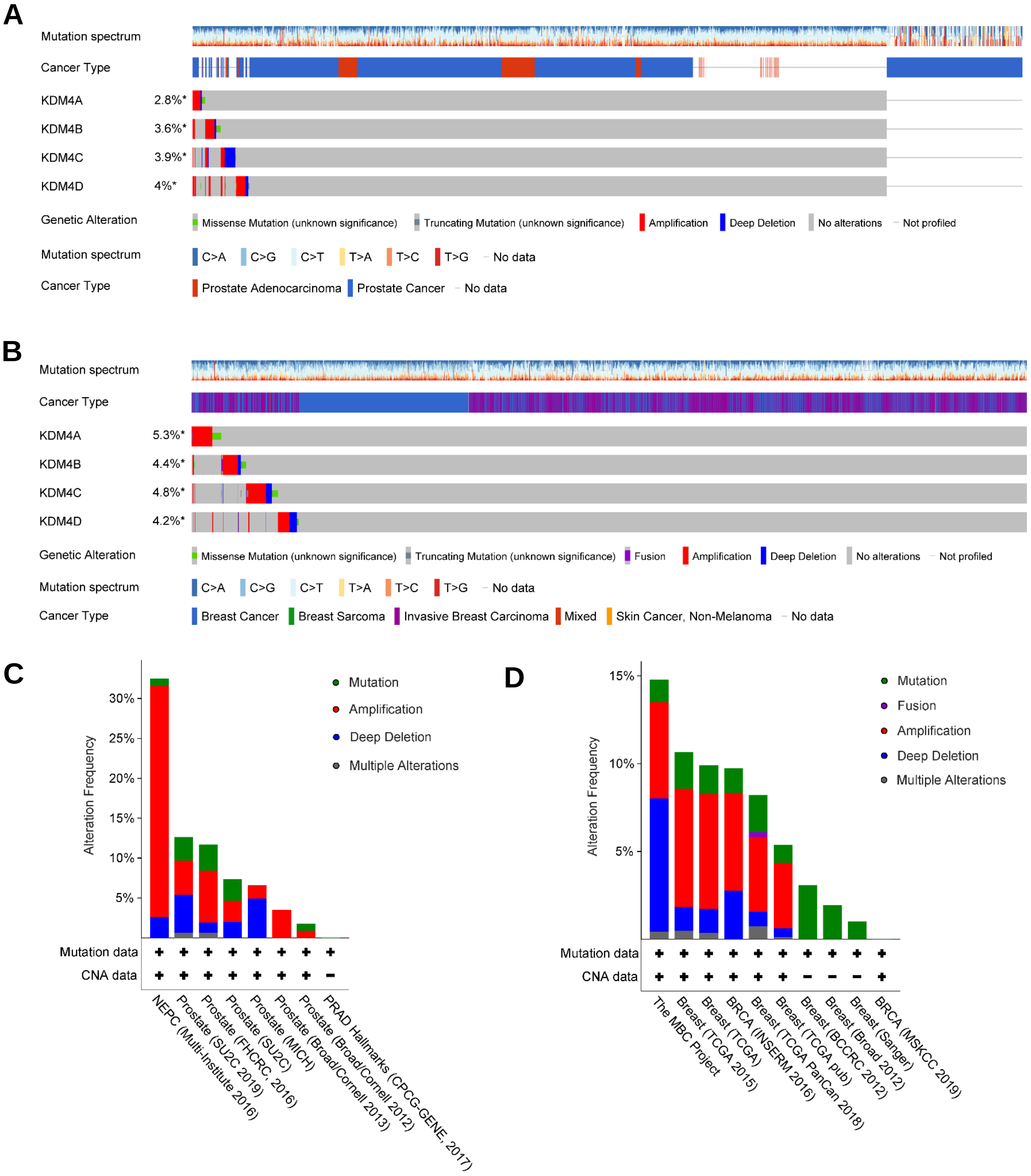

**Extended Data Fig. 1. Expression of KDM4 family is significantly upregulated in senescent cells in parallel to SASP development. Related to Fig. 1.**

A. GO enrichment profiling of upregulated genes in PSC27 cells upon genotoxicity-induced senescence.
B. Heatmap displaying genes correlated with histone methylation and significantly upregulated in senescent cells. CTRL, control. BLEO, bleomycin. Genes are listed by their expression fold change in CTRL *versus* BLEO cells.
C. Comparative transcript assay of KDM4 subfamily members (A/B/C/D) in presenescent (PRE) *versus* senescent (SEN) stromal cells. Signals were normalized to the PRE samples *per* gene.
D. Representative SA-β-Gal staining images of PSC27 cells upon therapy-induced senescence (TIS), replicative senescence (RS) or oncogene-induced senescence (OIS).
E. Comparative statistics of SA-β-Gal staining-based quantification of senescent PSC27 cells. Data organized for cases of TIS, RS or OIS.

**Extended Data Fig. 2. H3K9me3 and H3K36me3 levels decline in human prostate tumor stroma and is correlated with adverse clinical survival. Related to Fig. 2.**

A. Pathological assessment of stromal KDM4C expression in PCa samples (42 untreated *versus* 48 treated). In each group, patients were pathologically assigned into 4 categories per IHC staining intensity of KDM4C in tumor stroma. 1, negative; 2, weak; 3, moderate; 4, strong expression.
B. Pathological assessment of stromal KDM4D expression in PCa samples (42 untreated *versus* 48 treated). In each group, patients were pathologically assigned into 4 categories per IHC staining intensity of KDM4D in tumor stroma. 1, negative; 2, weak; 3, moderate; 4, strong expression.
C. Comparative analysis of KDM4C expression at transcription level between before and after chemotherapy. Data were organized for epithelial and stromal cells, respectively, after laser capture microdissection (LCM) of either cell lineage followed by quantitative analysis. Each dot represents an individual patient, with the data of “before” and “after” connected to allow direct profiling of KDM4C expression in the same individual patient.
D. Comparative analysis of KDM4D expression at transcription level between before and after chemotherapy. Data were organized for epithelial and stromal cells, respectively, after LCM of either cell lineage followed by quantitative analysis. Each dot represents an individual patient, with the data of “before” and “after” connected to allow direct profiling of KDM4D expression in the same individual patient.
E. Kaplan-Meier analysis of PCa patients. Disease-free survival (DFS) stratified according to KDM4C expression (low, average score $< 2$, dark green line, $n = 29$; high, average score $\geq 2$, dark red line, $n = 19$). DFS

represents the length (months) of period calculated from the date of PCa diagnosis to the point of first disease relapse. Survival curves generated according to the Kaplan–Meier method, with *P* value calculated using a log-rank (Mantel-Cox) test.

F. Kaplan-Meier analysis of PCa patients. Disease-free survival (DFS) stratified according to KDM4D expression (low, average score < 2, kelly line, n = 26; high, average score ≥ 2, purple line, n = 22). DFS represents the length (months) of period calculated from the date of PCa diagnosis to the point of first disease relapse. Survival curves generated according to the Kaplan–Meier method, with *P* value calculated by a log-rank (Mantel-Cox) test.

G. Histological images of H3K9me3 staining in the stroma of human prostate cancer (PCa) tissues. Upper, before chemotherapy; lower, after chemotherapy. Left panel, immunohistochemical (IHC) staining; right panel, hematoxylin and eosin (HE) staining. Rectangular regions *per* left images are amplified into corresponding right images. Scale bars, 100 μm.

H. Pathological assessment of stromal H3K9me3 signal intensity in PCa samples (42 untreated *versus* 48 treated). In each group, patients were pathologically assigned into 4 categories *per* IHC staining intensity of H3K9me3 in tumor stroma. 1, negative; 2, weak; 3, moderate; 4, strong expression.

I. Histological images of H3K36me3 staining in the stroma of human PCa tissues. Upper, before chemotherapy; lower, after chemotherapy. Left panel, IHC staining; right panel, HE staining. Rectangular regions *per* left images are amplified into corresponding right images. Scale bars, 100 μm.

J. Pathological assessment of stromal H3K36me3 signal intensity in PCa samples (42 untreated *versus* 48 treated). In each group, patients were pathologically assigned into 4 categories *per* IHC staining intensity of H3K9me3 in tumor stroma. 1, negative; 2, weak; 3, moderate; 4, strong expression.

K. Comparative analysis of H3K9me3/H3K36me3 signal intensity between before and after chemotherapy. Data were presented for epithelial and stromal cells, respectively, after laser capture microdissection (LCM) of either cell lineage followed by quantitative analysis. Each dot represents an individual patient, with the data of “before” and “after” connected to allow direct profiling of H3K9me3/H3K36me3 expression in the same individual patient.

L. Landscape mapping of pathological correlation between KDM4A/B and H3K9me3/H3K36me3 in the stroma of PCa patients after chemotherapy. Scores derived from assessment of molecule-specific IHC staining, with expression levels colored to reflect low (blue) *via* modest (turquoise) and fair (yellow) to high (red) signal intensity. Columns represent individual patients, rows different molecules. Totally 48 patients posttreatment were analyzed, with scores of each patient averaged from 3 independent pathological readings.

M. Statistical correlation between pathological scores of KDM4A/B and H3K9me3/H3K36me3 (Pearson analysis, r = 0.97; $P < 0.0001$ for both) in the 48 tumors with matching protein expression assessment data.
N. Kaplan-Meier analysis of PCa patients. Disease-free survival (DFS) stratified according to H3K9me3 intensity (low, average score < 2, green line, n = 26; high, average score ≥ 2, red line, n = 22). DFS represents the length (months) of period calculated from the date of PCa diagnosis to the point of first disease relapse. Survival curves generated according to the Kaplan–Meier method, with *P* value calculated using a log-rank (Mantel-Cox) test.
O. Kaplan-Meier analysis of PCa patients. Disease-free survival (DFS) stratified according to H3K36me3 expression (low, average score < 2, green line, n = 27; high, average score ≥ 2, red line, n = 21). DFS represents the length (months) of period calculated from the date of PCa diagnosis to the point of first disease relapse. Survival curves generated according to the Kaplan–Meier method, with *P* value calculated using a log-rank (Mantel-Cox) test.

Data in all bar plots are shown as mean ± SD and representative of 3 biological replicates. ^, $P > 0.05$. *, $P < 0.05$. **, $P < 0.01$. *** $P < 0.001$.

**Extended Data Fig. 3. Enhanced SASP expression and decreased H3K9/H3K36 methylation is regulated by histone demethylase KDM4B. Related to Fig. 4.**

A. Quantitative measurement of the SASP expression at transcription level. Stromal cells were transduced by a lentiviral construct encoding human KDM4B and/or exposed to BLEO treatment before analyzed. Signals normalized to CTRL cells (transduced with empty vector and untreated).
B. Immunoblot assay of DNA damage repair (DDR) signaling, H3K9/H3K36 methylation, and CXCL8 expression in stromal cells treated differently as described in (A). GPADH, loading control.
C. Immunoblot assay of DDR signaling, H3K9/H3K36 methylation, and CXCL8 expression in cells treated with BLEO and/or Chaetocin. GPADH, loading control.
D. Representative images of SA-β-Gal and BrdU staining of PSC27 treated in the ways described in (A).
E. Comparative statistics of SA-β-Gal and BrdU staining results of stromal cells in the individual conditions of (A). Left, SA-β-Gal staining. Right, BrdU staining.
F. Transcript expression of hallmark SASP factors in PSC27 sublines transduced with lentiviral constructs encoding shRNAs specific for KDM4B. Scrambled, transduction control. Cells subject to vehicle or BLEO treatment before analyzed.

G. Comparative statistics of SA-β-Gal staining results of stromal cells transduced with a lentiviral construct encoding human KDM4A or 4B and/or exposed to BLEO treatment before processed.
H. Comparative statistics of BrdU staining results of stromal cells transduced with a lentiviral construct encoding human KDM4A or 4B and/or exposed to BLEO treatment before processed.

**Extended Data Fig. 4. Targeting KDM4 changes expression profile and limits cancer malignancy conferred by senescent stromal cells. Related to Fig. 5.**

A. Immunoblot analysis of KDM4A/B expression, H3K9/H3K36 methylation, cell cycle arrest and CXCL8 expression in PSC27 treated by BLEO and/or ML324. GAPDH, loading control.
B. GSEA profiling of gene expression with significant enrichment scores exhibiting a NF-κB-specific signature in BLEO/ML324 co-treated cells compared with BLEO only-treated cells.
C. Growth curve assessment of PSC27 cells upon exposure to BLEO, ML324 or both. Cells were counted at the indicated time points after initiation of assays.
D. Heatmap depicting the influence of DNA damage and ML324, a small molecule inhibitor of KDM4, on transcriptomic expression profile of PSC27 cells. Genes displayed are downregulated upon BLEO-induced cellular senescence, and sorted by their expression fold change in response to ML324 treatment (in descending order).
E. Measurement of *in vitro* proliferation of prostate cancer (PCa) cells after exposure to the conditioned media (CM) of stromal cells treated by BLEO, ML324 or both.
F. Assessment of *in vitro* migration of PCa cells after exposure to the CM of stromal cells treated as described in (D).
G. Evaluation of *in vitro* invasion of PCa cells after exposure to the CM of stromal cells treated as described in (D).
H. Chemoresistance of PCa cells to the cytotoxic agent mitoxantrone (MIT) given at the IC50 value of individual PCa cell lines. Bottom, representative images of PC3 cells upon treatment with the CM as described in (D).

**Extended Data Fig. 5. The chromatin accessibility declines in senescent cells but is prevented upon ML324 treatment. Related to Fig. 6.**

A. Heatmap showing ATAC-seq enrichment of peaks near the accessible promoters (3.0 kb upstream and downstream TSS per gene) present in each of the assayed stromal samples (totally 9). Enrichment signals were collected for all active TSSs which were assorted by cap analysis of gene expression (CAGE) values, with peaks defined by hierarchical clustering.
B. Heatmap displaying representative genes whose accessibility decreased upon cellular senescence but increased when cells were treated by ML324.

C. The UCSC browser view shows enrichment of ATAC-seq signals near the promoters of CMMD9, PRR5L, TRAF6, RAG1 and IFTAP, genes not correlated with the SASP.
D. The UCSC browser views depict enrichment of ATAC-seq signals near the promoters of several SASP factors including AREG, SPINK1, MMP3, and WNT16B.
E. The UCSC browser view shows enrichment of ATAC-seq signals near the promoters of CDKN2A ($p16^{INK4a}$) and TP53 (p53), genes correlated with cellular senescence.

**Extended Data Fig. 6. Schematic design of preclinical trial, expression analysis of the SASP and KDM4A/B, and appraisal of treatment effects on body weight and biochemistry of animals. Related to Fig. 7.**

A. Schematic illustration of drug administration and tumor surveillance for preclinical trial. Cancer cells (PC3) alone or alongside stromal cells (PSC27) were inoculated subcutaneously to NOC/SCID mice 2 weeks prior to chemotherapy. MIT was provided on the 1st day of each week starting from the 3rd week, then given every other week with a total number of 3 doses. ML324 was provided 6 h before each time of MIT delivery. At the end of 8 weeks mice were sacrificed, tumor volume measured and tissues histologically assessed.
B. Transcript analysis of several canonical SASP factors including IL1α, MMP3, and GM-CSF expressed in the epithelial and stromal cells, respectively. Individual cell types were isolated from tumor tissues via LCM. Expression of $p16^{INK4a}$ and $p21^{CIP1}$ was measured to determine *in vivo* cellular senescence.
C. Comparative histological staining to evaluate KDM4A expression in tumor tissues. At the end of therapeutic regimen, animals were sacrificed, with tissues processed for immunohistochemical (IHC) staining.
D. Comparative histological staining to evaluate KDM4B expression in tumor tissues. Animals were subject to processes as described in (C).
E. Mouse body weight determination performed on a weekly basis for immunodeficient mice.
F. Serum measurement of creatinine, urea, alkaline phosphatase (ALP), and alanine aminotransferase (ALT) with terminal bleeds (cardiac punctures) taken at the end of therapeutic regimens.
G. Mouse body weight determination performed on a weekly basis for immunocompetent mice (C57BL/6 strain).
H. Serum measurement of creatinine, urea, alkaline phosphatase (ALP), and alanine aminotransferase (ALT) with terminal bleeds (cardiac punctures) taken at the end of therapeutic regimens for immunocompetent mice.
I. Routine analysis of peripheral blood. The circulating levels of hemoglobin, white blood cells, lymphocytes and platelets at the end of each therapeutic regimen were assessed.

Data are shown as mean ± SD and representative of 3 independent experiments. MIT, mitoxantrone. WBC, white blood count. N = 5 per treatment arm. ^*P* > 0.05.

**Extended Data Fig. 7. Schematic diagram, molecular profiles and clinical attributes of human KDM4 alterations with datasets extracted from large-scale cancer genomics. Related to Fig. 8.**

A. OncoPrint view of human KDM4 alterations. Multidimensional cancer genomics datasets of KDM4 mutations in human prostate cancer (PCa) patients were mapped. Data from 3317 patients (3480 biospecimens) out of 10 individual clinical studies were subject to analysis and visualization, with source data from the Cancer Genome Atlas (TCGA), a landmark cancer genomics program. Top columns, individual patients affected by particular alterations.
B. OncoPrint view of similar genomics mapping presentation of human KDM4 alterations in breast cancer (BCa) patients as described in (B). Data from 4559 patients (4625 biospecimens) out of 10 individual clinical studies were incorporated. Source data derived from the TCGA portal.
C. Cancer type-specific summary of human KDM4 mutations in PCa patients. Data were pooled from 8 separate studies that reported single-site mutations, amplifications, deep deletions and multiple alterations of KDM4.
D. Cancer type-specific summary of human KDM4 mutations in BCa patients. Data were pooled from 10 separate studies that reported single-site mutations, fusions, amplifications, deep deletions and multiple alterations of KDM4.

**Supplementary Table 1. Mass spectrometric profiling of senescent cell proteomics. Related to Fig. 1.**

**A. MS/MS spectrum database search analysis summary**

| Total spectrums | Matched spectrums | Peptides | Unique peptides | Identified proteins | Quantifiable proteins |
|---|---|---|---|---|---|
| 16495 | 7694 (46.6%) | 4110 | 3803 | 732 | 447 |

**B. Differentially expressed protein summary**
**(Filtered with threshold value of expression fold change and *P* value < 0.05)**

| Compare group | Regulated type | fold change >1.2 | fold change >1.3 | fold change >1.5 | fold change >2 |
|---|---|---|---|---|---|
| SEN/CTRL | up-regulated | 218 | 194 | 156 | 87 |
| | down-regulated | 125 | 106 | 79 | 29 |

#### Supplementary Table 2. Intracellular proteins identified by mass spectrometry. Related to Fig. 1.

| Protein accession | Gene name | MW [kDa] | Score | Coverage [%] | #Peptides | #PSMs | #Unique peptide | CTRL | SEN | SEN/CTRL Ratio | Subcellular localization | KEGG KO No. | KOG category | KOG NO. | KOG description |
|---|---|---|---|---|---|---|---|---|---|---|---|---|---|---|---|
| P04179 | SOD2 | 24.75 | 39.431 | 31.1 | 5 | 8 | 5 | 8451300 | 3.25E+08 | 13.124 | mitochondria | K04564 | P | KOG0876 | Manganese superoxide dismutase |
| P53007 | SLC25A1 | 34.012 | 17.126 | 11.6 | 3 | 3 | 3 | 6521900 | 1.06E+08 | 7.788 | plasma membrane | K15100 | C | KOG0756 | Mitochondrial tricarboxylate/dicarboxylate carrier proteins |
| P24844 | MYL9 | 19.827 | 38.661 | 56.4 | 11 | 30 | 3 | 19644000 | 7.47E+08 | 7.272 | mitochondria | K12755 | Z | KOG0031 | Myosin regulatory light chain, EF-Hand protein superfamily |
| O00479 | HMGN4 | 9.5388 | 13.977 | 16.7 | 2 | 3 | 2 | 54708000 | 2.29E+08 | 5.207 | nucleus | K11302 | | | |
| O00159 | MYO1C | 121.68 | 293.12 | 42.2 | 34 | 43 | 34 | 2.29E+08 | 3.93E+09 | 4.574 | cytoplasm | K10356 | Z | KOG0164 | Myosin class I heavy chain |
| Q9H4G4 | GLIPR2 | 17.218 | 20.284 | 37 | 3 | 3 | 3 | 22738000 | 1.3E+08 | 4.032 | cytoplasm | | S | KOG3017 | Defense-related protein containing SCP domain |
| P25705 | ATP5F1A | 59.75 | 152.64 | 38 | 15 | 19 | 15 | 1.33E+08 | 1.5E+09 | 3.877 | mitochondria | K02132 | C | KOG1353 | F0F1-type ATP synthase, alpha subunit |
| P80723 | BASP1 | 22.693 | 88.605 | 75.3 | 11 | 15 | 11 | 1.93E+08 | 1.47E+09 | 3.766 | nucleus | K17272 | | | |
| P08582 | MELTF | 80.214 | 138.98 | 40.4 | 16 | 22 | 16 | 54881000 | 1.22E+09 | 3.725 | extracellular | K06569 | | | |
| Q8NBQ5 | HSD17B11 | 32.935 | 93.938 | 45 | 10 | 14 | 10 | 57261000 | 5.51E+08 | 3.637 | endoplasmic reticulum | | Q | KOG1201 | Hydroxysteroid 17-beta dehydrogenase 11 |
| P48047 | ATP5O | 23.277 | 22.969 | 17.4 | 3 | 4 | 3 | 19552000 | 1.1E+08 | 3.591 | mitochondria | K02137 | C | KOG1662 | Mitochondrial F1F0-ATP synthase, subunit OSCP/ATP5 |
| P21589 | NT5E | 63.367 | 103.23 | 38.7 | 14 | 19 | 14 | 99343000 | 8.05E+08 | 3.55 | peroxisome | K19970 | F | KOG4419 | 5' nucleotidase |
| P27824 | CANX | 67.567 | 24.663 | 12.3 | 4 | 5 | 4 | 15641000 | 1.22E+08 | 3.534 | endoplasmic reticulu | K08054 | O | KOG0675 | Calnexin |
| P02545 | LMNA | 74.139 | 176.93 | 45.2 | 20 | 23 | 20 | 88713000 | 9.26E+08 | 3.529 | nucleus | K12641 | DY | KOG0977 | Nuclear envelope protein lamin, intermediate filament superfamily |
| P52926 | HMGA2 | 11.832 | 31.678 | 40.4 | 3 | 5 | 3 | 16460000 | 1.07E+08 | 3.524 | nucleus | K09283 | | | |
| Q56VL3 | OCIAD2 | 16.953 | 36.745 | 43.5 | 5 | 5 | 5 | 19218000 | 2.44E+08 | 3.498 | extracellular | | | | |
| P06576 | ATP5F1B | 56.559 | 248.4 | 68.8 | 22 | 33 | 22 | 4.87E+08 | 4.34E+09 | 3.378 | mitochondria | K02133 | C | KOG1350 | F0F1-type ATP synthase, beta subunit |
| P30838 | ALDH3A1 | 50.394 | 58.004 | 24.3 | 7 | 10 | 7 | 1.01E+08 | 3.29E+08 | 3.296 | mitochondria | K00129 | C | KOG2456 | Aldehyde dehydrogenase |
| P56385 | ATP5ME | 7.9331 | 23.651 | 52.2 | 4 | 4 | 4 | 9839500 | 1.13E+08 | 3.254 | mitochondria | K02129 | C | KOG4326 | Mitochondrial F1F0-ATP synthase, subunit e |
| Q96N66 | MBOAT7 | 52.764 | 28.204 | 10.8 | 3 | 3 | 3 | 12169000 | 1.16E+08 | 3.244 | plasma membrane | K13516 | S | KOG2706 | Predicted membrane protein |
| P09493 | TPM1 | 32.708 | 323.31 | 53.2 | 27 | 37 | 9 | 9.32E+08 | 9.19E+09 | 3.19 | cytoplasm | K10373 | | | |
| P09525 | ANXA4 | 35.882 | 72.36 | 28.8 | 7 | 7 | 7 | 54065000 | 4.89E+08 | 3.18 | cytoplasm | K17093 | U | KOG0819 | Annexin |
| P47755 | CAPZA2 | 32.949 | 35.751 | 30.4 | 5 | 6 | 4 | 40361000 | 2.26E+08 | 3.176 | cytoplasm | K10364 | Z | KOG0836 | F-actin capping protein, alpha subunit |
| P07858 | CTSB | 37.821 | 45.797 | 25.1 | 5 | 6 | 5 | 16233000 | 1.97E+08 | 3.153 | extracellular | K01363 | O | KOG1543 | Cysteine proteinase Cathepsin L |
| P67936 | TPM4 | 28.521 | 119.29 | 66.1 | 21 | 19 | 10 | 1.74E+08 | 1.72E+09 | 3.068 | cytoplasm | K10375 | | | |
| P04792 | HSPB1 | 22.782 | 56.634 | 57.6 | 8 | 12 | 8 | 76335000 | 7.93E+08 | 3.047 | nucleus | K04455 | O | KOG3591 | Alpha crystallins |
| Q16795 | NDUFA9 | 42.509 | 31.695 | 17 | 5 | 5 | 5 | 18465000 | 1.28E+08 | 3.003 | mitochondria | K03953 | C | KOG2865 | NADH:ubiquinone oxidoreductase, NDUFA9/39kDa subunit |
| Q9UM54 | MYO6 | 149.69 | 116.21 | 20.4 | 15 | 18 | 15 | 36694000 | 8.46E+08 | 2.984 | cytoplasm | K10358 | Z | KOG0163 | Myosin class VI heavy chain |
| P60903 | S100A10 | 11.203 | 17.953 | 35.1 | 2 | 5 | 2 | 61356000 | 5.37E+08 | 2.952 | cytoplasm | K17274 | | | |
| Q9Y608 | LRRFIP2 | 82.17 | 18.271 | 6.8 | 2 | 3 | 2 | 10794000 | 1.41E+08 | 2.924 | nucleus | | R | KOG2010 | Double stranded RNA binding protein |
| Q6NZI2 | CAVIN1 | 43.476 | 68.981 | 24.4 | 6 | 8 | 6 | 20151000 | 1.7E+08 | 2.922 | nucleus | K19387 | | | |
| P06753 | TPM3 | 32.95 | 77.878 | 46.3 | 24 | 28 | 7 | 1.5E+08 | 1.41E+09 | 2.875 | cytoplasm | K09290 | Z | KOG1003 | Actin filament-coating protein tropomyosin |
| Q9Y4I1 | MYO5A | 215.4 | 149 | 15 | 19 | 22 | 19 | 86320000 | 7.27E+08 | 2.853 | cytoplasm | K10357 | Z | KOG0161 | Myosin class II heavy chain |
| Q9P0K7 | RAI14 | 110.04 | 323.31 | 50.3 | 34 | 45 | 34 | 3.51E+08 | 2.35E+09 | 2.767 | nucleus | | | | |
| P39656 | DDOST | 50.8 | 46.385 | 20.6 | 6 | 6 | 6 | 24268000 | 2.9E+08 | 2.724 | plasma membrane | K12670 | O | KOG2754 | Oligosaccharyltransferase, beta subunit |
| P56134 | ATP5J2 | 10.918 | 16.065 | 25.5 | 2 | 2 | 2 | 9911200 | 1.17E+08 | 2.709 | cytoplasm | K02130 | C | KOG4092 | Mitochondrial F1F0-ATP synthase, subunit f |
| Q14257 | RCN2 | 36.876 | 11.758 | 11 | 2 | 2 | 2 | 10436000 | 44429000 | 2.703 | extracellular | | TU | KOG4223 | Reticulocalbin, calumenin, DNA supercoiling factor, and related Ca2+-binding proteins of the CREC family (EF-Hand protein superfamily) |
| Q15165 | PON2 | 39.38 | 42.55 | 35 | 5 | 5 | 5 | 12239000 | 1.7E+08 | 2.7 | extracellular | K01045 | | | |
| P13987 | CD59 | 14.177 | 38.046 | 25.8 | 5 | 7 | 5 | 31798000 | 2.08E+08 | 2.678 | extracellular | K04008 | | | |
| P51608 | MECP2 | 52.44 | 67.069 | 24.7 | 9 | 10 | 9 | 40833000 | 3.08E+08 | 2.665 | nucleus | K11588 | KB | KOG4161 | Methyl-CpG binding transcription regulators |
| P23141 | CES1 | 62.52 | 35.558 | 12.3 | 5 | 7 | 5 | 13822000 | 1.78E+08 | 2.661 | extracellular | K01044 | R | KOG1516 | Carboxylesterase and related proteins |
| Q6WCQ1 | MPRIP | 116.53 | 323.31 | 40.7 | 32 | 40 | 32 | 5.72E+08 | 2.44E+09 | 2.643 | nucleus | | Z | KOG4807 | F-actin binding protein, regulates actin cytoskeletal organization |
| P36957 | DLST | 48.755 | 39.504 | 18.1 | 5 | 5 | 5 | 20952000 | 1.17E+08 | 2.621 | mitochondria | K00658 | C | KOG0559 | Dihydrolipoamide succinyltransferase (2-oxoglutarate dehydrogenase, E2 subunit) |
| P04843 | RPN1 | 68.569 | 41.954 | 14.7 | 6 | 6 | 6 | 18973000 | 1.15E+08 | 2.613 | endoplasmic reticulu | K12666 | O | KOG2291 | Oligosaccharyltransferase, alpha subunit (ribophorin I) |
| P10301 | RRAS | 23.48 | 29.383 | 28.4 | 4 | 4 | 4 | 16909000 | 1.31E+08 | 2.584 | nucleus | K07829 | R | KOG0395 | Ras-related GTPase |
| P18206 | VCL | 123.8 | 32.739 | 8.9 | 5 | 5 | 5 | 11979000 | 1.14E+08 | 2.561 | cytoplasm | K05700 | W | KOG3681 | Alpha-catenin |

| | | | | | | | | | | | | | | | |
|---|---|---|---|---|---|---|---|---|---|---|---|---|---|---|---|
| P11233 | RALA | 23.567 | 28.725 | 25.2 | 4 | 4 | 4 | 6799500 | 87280000 | 2.553 | cytoplasm | K07834 | R | KOG0395 | Ras-related GTPase |
| Q13045 | FLII | 144.75 | 32.266 | 5.1 | 5 | 5 | 5 | 12253000 | 86182000 | 2.544 | cytoplasm | | Z | KOG0444 | Cytoskeletal regulator Flightless-I (contains leucine-rich and gelsolin repeats) |
| O60506 | SYNCRIP | 69.602 | 53.512 | 22.2 | 8 | 9 | 6 | 82070000 | 2.82E+08 | 2.535 | nucleus | K13160 | A | KOG0117 | Heterogeneous nuclear ribonucleoprotein R (RRM superfamily) |
| P21333 | FLNA | 280.74 | 323.31 | 36.4 | 56 | 75 | 54 | 7.97E+08 | 4.37E+09 | 2.534 | cytoplasm | K04437 | Z | KOG0518 | Actin-binding cytoskeleton protein, filamin |
| O75368 | SH3BGRL | 12.774 | 47.672 | 60.5 | 5 | 7 | 5 | 44555000 | 2.6E+08 | 2.512 | mitochondria | | S | KOG4023 | Uncharacterized conserved protein |
| P09936 | UCHL1 | 24.824 | 47.314 | 32.3 | 4 | 5 | 4 | 37857000 | 2.45E+08 | 2.473 | cytoplasm | K05611 | O | KOG1415 | Ubiquitin C-terminal hydrolase UCHL1 |
| Q9P0M6 | H2AFY2 | 40.058 | 92.939 | 52.2 | 13 | 16 | 11 | 75946000 | 3.21E+08 | 2.443 | mitochondria | K11251 | BK | KOG2633 | Hismacro and SEC14 domain-containing proteins |
| Q9H9B4 | SFXN1 | 35.619 | 7.1511 | 12.1 | 2 | 3 | 1 | 13476000 | 49510000 | 2.398 | cytoplasm | | R | KOG3767 | Sideroflexin |
| O15231 | ZNF185 | 73.525 | 33.738 | 10.4 | 5 | 5 | 5 | 15110000 | 1.46E+08 | 2.389 | cytoplasm,nucleus | | | | |
| O00151 | PDLIM1 | 36.071 | 53.529 | 46.5 | 7 | 7 | 7 | 4.29E+08 | 2.26E+08 | 2.379 | cytoplasm | | TZ | KOG1703 | Adaptor protein Enigma and related PDZ-LIM proteins |
| P21980 | TGM2 | 77.328 | 35.084 | 13.4 | 5 | 5 | 5 | 18150000 | 91747000 | 2.352 | cytoplasm | K05625 | | | |
| O75947 | ATP5H | 18.491 | 25.399 | 37.3 | 4 | 5 | 4 | 20152000 | 2.08E+08 | 2.344 | cytoplasm | K02138 | C | KOG3366 | Mitochondrial F1F0-ATP synthase, subunit d/ATP7 |
| P46821 | MAP1B | 270.63 | 20.901 | 1.7 | 3 | 3 | 3 | 10930000 | 62882000 | 2.328 | nucleus | K10429 | Z | KOG3592 | Microtubule-associated proteins |
| P42330 | AKR1C3 | 36.853 | 71.259 | 45.8 | 9 | 9 | 6 | 42453000 | 2.33E+08 | 2.322 | cytoplasm | K04119 | R | KOG1577 | Aldo/keto reductase family proteins |
| P63000 | RAC1 | 21.45 | 29.252 | 25 | 4 | 7 | 4 | 38019000 | 2.81E+08 | 2.303 | cytoplasm | K04392 | | | |
| O43795 | MYO1B | 131.98 | 299.66 | 36.4 | 32 | 46 | 32 | 3.26E+09 | 3.91E+09 | 2.3 | cytoplasm | K10356 | Z | KOG0164 | Myosin class I heavy chain |
| O75369 | FLNB | 278.16 | 153.87 | 14.3 | 23 | 23 | 21 | 1.7E+08 | 4.45E+08 | 2.289 | cytoplasm,nucleus | K04437 | Z | KOG0518 | Actin-binding cytoskeleton protein, filamin |
| Q13838 | DDX39B | 48.991 | 61.277 | 29.9 | 9 | 11 | 9 | 57869000 | 4.2E+08 | 2.279 | cytoplasm,nucleus | K12812 | A | KOG0329 | ATP-dependent RNA helicase |
| P27105 | STOM | 31.73 | 85.013 | 42 | 7 | 11 | 7 | 1.52E+08 | 1.17E+09 | 2.268 | cytoplasm | K17286 | C | KOG2621 | Prohibitins and stomatins of the PID superfamily |
| P04899 | GNAI2 | 40.45 | 54.268 | 27.6 | 7 | 8 | 4 | 99363000 | 4.2E+08 | 2.261 | cytoplasm | K04630 | DT | KOG0082 | G-protein alpha subunit (small G protein superfamily) |
| Q9BWM7 | SFXN3 | 35.503 | 19.391 | 12.8 | 3 | 3 | 2 | 23716000 | 1.33E+08 | 2.253 | cytoplasm | | R | KOG3767 | Sideroflexin |
| Q16881 | TXNRD1 | 70.905 | 19.137 | 5.9 | 2 | 2 | 2 | 12954000 | 70308000 | 2.238 | cytoplasm | K22182 | O | KOG4716 | Thioredoxin reductase |
| P14923 | JUP | 81.744 | 56.36 | 17.7 | 7 | 8 | 7 | 33753000 | 2.4E+08 | 2.229 | cytoplasm | K10056 | TZ | KOG4203 | Armadillo/beta-Catenin/plakoglobin |
| P14866 | HNRNPL | 64.132 | 73.749 | 28.5 | 8 | 9 | 8 | 72239000 | 4.14E+08 | 2.2 | nucleus | K13159 | A | KOG1456 | Heterogeneous nuclear ribonucleoprotein L (contains RRM repeats) |
| P62873 | GNB1 | 37.377 | 68.028 | 32.9 | 7 | 9 | 4 | 51229000 | 3.13E+08 | 2.189 | cytoplasm | K04536 | R | KOG0286 | G-protein beta subunit |
| O75489 | NDUFS3 | 30.241 | 50.015 | 31.1 | 6 | 6 | 6 | 27136000 | 90619000 | 2.178 | mitochondria | K03936 | C | KOG1713 | NADH-ubiquinone oxidoreductase, NDUFS3/30 kDa subunit |
| P15121 | AKR1B1 | 35.853 | 96.443 | 55.1 | 11 | 15 | 11 | 2.89E+08 | 1.63E+09 | 2.175 | cytoplasm | K00011 | R | KOG1577 | Aldo/keto reductase family proteins |
| Q9UBI6 | GNG12 | 8.0061 | 21.612 | 56.9 | 3 | 3 | 3 | 52453000 | 3.5E+08 | 2.155 | nucleus | K04347 | T | KOG4119 | G protein gamma subunit |
| P10620 | MGST1 | 17.598 | 38.185 | 20 | 4 | 6 | 4 | 68874000 | 4.38E+08 | 2.145 | cytoplasm,nucleus | K00799 | | | |
| P35580 | MYH10 | 229 | 323.31 | 38.8 | 72 | 109 | 40 | 8.41E+08 | 5.69E+09 | 2.132 | cytoplasm | K10352 | Z | KOG0161 | Myosin class II heavy chain |
| P37802 | TAGLN2 | 22.391 | 71.639 | 52.3 | 9 | 10 | 9 | 1.18E+08 | 6.42E+08 | 2.127 | cytoplasm,nucleus | K20526 | Z | KOG2046 | Calponin |
| P40199 | CEACAM6 | 37.194 | 17.137 | 5.8 | 2 | 3 | 2 | 12938000 | 40194000 | 2.11 | extracellular | K06499 | | | |
| Q09666 | AHNAK | 629.09 | 120.14 | 12.9 | 18 | 21 | 18 | 2.08E+08 | 6.63E+08 | 2.109 | nucleus | | | | |
| Q13813 | SPTAN1 | 284.54 | 62.961 | 5.4 | 7 | 7 | 7 | 46211000 | 1.39E+08 | 2.081 | nucleus | K06114 | Z | KOG0035 | Ca2+-binding actin-bundling protein (actinin), alpha chain |
| Q6NUK1 | SLC25A24 | 53.354 | 37.688 | 16.4 | 6 | 6 | 6 | 26640000 | 1.54E+08 | 2.06 | cytoplasm | K14684 | F | KOG0036 | Predicted mitochondrial carrier protein |
| P26038 | MSN | 67.819 | 32.027 | 10.2 | 5 | 5 | 5 | 13531000 | 1.09E+08 | 2.052 | cytoplasm | K05763 | R | KOG3529 | Radixin, moesin and related proteins of the ERM family |
| P51970 | NDUFA8 | 20.105 | 24.495 | 30.2 | 3 | 4 | 3 | 70760000 | 1.43E+08 | 2.052 | extracellular | K03952 | C | KOG3458 | NADH:ubiquinone oxidoreductase, NDUFA8/PGIV/19 kDa subunit |
| Q9NYL9 | TMOD3 | 39.594 | 177.16 | 75 | 20 | 30 | 19 | 5.01E+08 | 3.33E+09 | 2.04 | cytoplasm | K10370 | Z | KOG3735 | Tropomodulin and leiomodulin |
| Q07065 | CKAP4 | 66.022 | 80.037 | 23.8 | 10 | 11 | 10 | 2.63E+08 | 3.76E+08 | 2.019 | plasma membrane | K13999 | | | |
| Q969G5 | CAVIN3 | 27.701 | 14.168 | 7.7 | 2 | 2 | 2 | 9055800 | 40254000 | 2.011 | extracellular | | | | |
| O60675 | MAFK | 17.523 | 21.419 | 24.4 | 3 | 3 | 3 | 23843000 | 97765000 | 2.009 | nucleus | K09037 | | | |
| P0DMV9 | HSPA1B | 70.051 | 66.554 | 27.3 | 10 | 14 | 6 | 1.27E+08 | 4.33E+08 | 1.996 | cytoplasm | K03283 | | | |
| O43390 | HNRNPR | 70.942 | 35.432 | 15.8 | 7 | 9 | 5 | 45012000 | 2.25E+08 | 1.992 | nucleus | K13161 | A | KOG0117 | Heterogeneous nuclear ribonucleoprotein R (RRM superfamily) |
| Q16719 | KYNU | 52.351 | 38.241 | 19.1 | 6 | 8 | 6 | 34369000 | 1.44E+08 | 1.982 | cytoplasm | K01556 | E | KOG3846 | L-kynurenine hydrolase |
| P49411 | TUFM | 49.541 | 32.492 | 15.5 | 5 | 5 | 5 | 14538000 | 1.06E+08 | 1.97 | mitochondria | K02358 | J | KOG0460 | Mitochondrial translation elongation factor Tu |
| P14649 | MYL6B | 22.764 | 42.857 | 49 | 7 | 11 | 5 | 57876000 | 2.48E+08 | 1.951 | cytoplasm | K12751 | Z | KOG0030 | Myosin essential light chain, EF-Hand protein superfamily |
| Q13435 | SF3B2 | 100.23 | 58.085 | 15.5 | 9 | 9 | 9 | 42298000 | 2.72E+08 | 1.944 | nucleus | K12829 | A | KOG2330 | Splicing factor 3b, subunit 2 |
| Q9Y5S9 | RBM8A | 19.889 | 12.701 | 33.3 | 2 | 2 | 2 | 9105600 | 84319000 | 1.944 | nucleus | K12876 | R | KOG0130 | RNA-binding protein RBM8/Tsunagi (RRM superfamily) |
| P07099 | EPHX1 | 52.948 | 15.98 | 8.4 | 2 | 2 | 2 | 20502000 | 1.22E+08 | 1.922 | endoplasmic reticulu | K01253 | R | KOG2565 | Predicted hydrolases or acyltransferases (alpha/beta hydrolase superfamily) |
| P11413 | G6PD | 59.256 | 91.487 | 31.5 | 11 | 13 | 11 | 82504000 | 3.68E+08 | 1.909 | cytoplasm | K00036 | G | KOG0563 | Glucose-6-phosphate 1-dehydrogenase |
| Q9UHB6 | LIMA1 | 85.225 | 24.407 | 10 | 4 | 4 | 4 | 17440000 | 86467000 | 1.905 | nucleus | | | | |

| | | | | | | | | | | | | | | |
|---|---|---|---|---|---|---|---|---|---|---|---|---|---|---|
| Q7Z7K6 | CENPV | 29.946 | 29.17 | 20.4 | 4 | 6 | 4 | 84610000 | 3.33E+08 | 1.902 | mitochondria | | S | KOG4192 | Uncharacterized conserved protein |
| P13804 | ETFA | 35.079 | 11.974 | 6.9 | 2 | 2 | 2 | 14341000 | 31836000 | 1.894 | mitochondria | K03522 | C | KOG3954 | Electron transfer flavoprotein, alpha subunit |
| P61160 | ACTR2 | 44.76 | 28.994 | 13.7 | 4 | 4 | 4 | 12663000 | 60574000 | 1.891 | cytoplasm | K17260 | Z | KOG0677 | Actin-related protein Arp2/3 complex, subunit Arp2 |
| P52895 | AKR1C2 | 36.735 | 24.26 | 26.6 | 5 | 7 | 2 | 48545000 | 2.78E+08 | 1.884 | cytoplasm | K00089 | R | KOG1577 | Aldo/keto reductase family proteins |
| P09211 | GSTP1 | 23.356 | 28.424 | 26.2 | 3 | 4 | 3 | 46123000 | 2.34E+08 | 1.882 | mitochondria | K00799 | O | KOG1695 | Glutathione S-transferase |
| P08754 | GNAI3 | 40.532 | 15.629 | 21.5 | 5 | 6 | 2 | 11410000 | 76918000 | 1.88 | cytoplasm | K04630 | DT | KOG0082 | G-protein alpha subunit (small G protein superfamily) |
| Q15459 | SF3A1 | 88.885 | 39.2 | 12.6 | 5 | 6 | 5 | 83465000 | 2.88E+08 | 1.863 | cytoplasm | K12825 | A | KOG0007 | Splicing factor 3a, subunit 1 |
| P63010 | AP2B1 | 104.55 | 37.675 | 12 | 6 | 6 | 6 | 28607000 | 1.82E+08 | 1.859 | plasma membrane | K11825 | U | KOG1061 | Vesicle coat complex AP-1/AP-2/AP-4, beta subunit |
| P30044 | PRDX5 | 22.086 | 26.284 | 23.8 | 4 | 5 | 4 | 28717000 | 1.03E+08 | 1.858 | cytoplasm | K11187 | O | KOG0541 | Alkyl hydroperoxide reductase/peroxiredoxin |
| Q16718 | NDUFA5 | 13.459 | 19.338 | 31 | 2 | 2 | 2 | 17446000 | 97836000 | 1.857 | cytoplasm | K03949 | C | KOG3365 | NADH:ubiquinone oxidoreductase, NDUFA5/B13 subunit |
| P12814 | ACTN1 | 103.06 | 255.21 | 63.3 | 42 | 61 | 27 | 5.24E+08 | 2.37E+09 | 1.849 | cytoplasm,nucleus | K05699 | Z | KOG0035 | Ca2+-binding actin-bundling protein (actinin), alpha chain |
| O60832 | DKC1 | 57.673 | 17.235 | 8.8 | 2 | 2 | 2 | 11602000 | 65577000 | 1.841 | cytoplasm | K11131 | J | KOG2529 | Pseudouridine synthase |
| Q9UL46 | PSME2 | 27.401 | 24.073 | 21.3 | 3 | 3 | 3 | 22139000 | 1.14E+08 | 1.84 | cytoplasm | K06697 | O | KOG4470 | Proteasome activator subunit |
| Q96IZ0 | PAWR | 36.567 | 36.379 | 27.1 | 4 | 5 | 4 | 17198000 | 1.5E+08 | 1.836 | nucleus | | | | |
| Q00325 | SLC25A3 | 40.094 | 45.28 | 16.6 | 6 | 6 | 6 | 54770000 | 1.55E+08 | 1.83 | nucleus | K15102 | C | KOG0767 | Mitochondrial phosphate carrier protein |
| P35232 | PHB | 29.804 | 48.582 | 34.9 | 6 | 7 | 6 | 3.08E+08 | 4.96E+08 | 1.818 | cytoplasm | K17080 | O | KOG3083 | Prohibitin |
| P50454 | SERPINH1 | 46.44 | 29.471 | 12.7 | 3 | 3 | 3 | 12508000 | 89840000 | 1.807 | extracellular | K09501 | V | KOG2392 | Serpin |
| Q03135 | CAV1 | 20.471 | 61.532 | 46.6 | 7 | 10 | 7 | 2.45E+08 | 1.48E+09 | 1.806 | cytoplasm | K06278 | | | |
| O43920 | NDUFS5 | 12.517 | 33.37 | 47.2 | 5 | 5 | 5 | 34445000 | 1.07E+08 | 1.804 | extracellular | K03938 | C | KOG4110 | NADH:ubiquinone oxidoreductase, NDUFS5/15kDa |
| P09669 | COX6C | 8.7813 | 22.73 | 36 | 3 | 4 | 3 | 35359000 | 1.27E+08 | 1.804 | mitochondria | K02268 | | | |
| Q02978 | SLC25A11 | 34.061 | 46.042 | 23.9 | 5 | 6 | 5 | 28894000 | 1.49E+08 | 1.789 | cytoplasm | K15104 | C | KOG0759 | Mitochondrial oxoglutarate/malate carrier proteins |
| Q969G3 | SMARCE1 | 46.649 | 20.577 | 6.8 | 2 | 2 | 2 | 9131100 | 25623000 | 1.761 | nucleus | K11651 | B | KOG4715 | SWI/SNF-related matrix-associated actin-dependent regulator of chromatin |
| P00352 | ALDH1A1 | 54.861 | 208.82 | 57.9 | 21 | 30 | 21 | 5.13E+08 | 2.95E+09 | 1.759 | cytoplasm | K07249 | C | KOG2450 | Aldehyde dehydrogenase |
| P62491 | RAB11A | 24.393 | 15.107 | 10.6 | 2 | 2 | 2 | 13014000 | 39901000 | 1.758 | cytoskeleton | K07904 | U | KOG0087 | GTPase Rab11/YPT3, small G protein superfamily |
| P47985 | UQCRFS1 | 29.668 | 41.547 | 35.4 | 5 | 6 | 5 | 41846000 | 2.31E+08 | 1.743 | mitochondria | K00411 | C | KOG1671 | Ubiquinol cytochrome c reductase, subunit RIP1 |
| P51991 | HNRNPA3 | 39.594 | 51.01 | 24.6 | 8 | 10 | 8 | 2.03E+08 | 7.15E+08 | 1.736 | nucleus | K12741 | R | KOG0118 | FOG: RRM domain |
| Q9H299 | SH3BGRL3 | 10.438 | 16.384 | 20.4 | 1 | 2 | 1 | 31614000 | 1.53E+08 | 1.735 | mitochondria | | S | KOG4023 | Uncharacterized conserved protein |
| Q15717 | ELAVL1 | 36.091 | 57.444 | 32.2 | 7 | 8 | 7 | 72728000 | 2.79E+08 | 1.732 | nucleus | K13088 | A | KOG0145 | RNA-binding protein ELAV/HU (RRM superfamily) |
| P30101 | PDIA3 | 56.782 | 27.848 | 14.1 | 4 | 4 | 4 | 14117000 | 70941000 | 1.731 | endoplasmic reticulu | K08056 | O | KOG0190 | Protein disulfide isomerase (prolyl 4-hydroxylase beta subunit) |
| P46940 | IQGAP1 | 189.25 | 109.82 | 14.5 | 14 | 15 | 14 | 87606000 | 4.34E+08 | 1.729 | cytoplasm | K16848 | T | KOG2128 | Ras GTPase-activating protein family - IQGAP |
| P51149 | RAB7A | 23.489 | 18.513 | 14 | 2 | 2 | 2 | 10459000 | 32514000 | 1.719 | cytoplasm | K07897 | R | KOG0394 | Ras-related GTPase |
| O60218 | AKR1B10 | 36.019 | 117.22 | 58.5 | 13 | 18 | 13 | 2.58E+08 | 9.52E+08 | 1.704 | cytoplasm | K00011 | R | KOG1577 | Aldo/keto reductase family proteins |
| Q14103 | HNRNPD | 38.434 | 61.25 | 24.5 | 8 | 10 | 7 | 1.46E+08 | 5.22E+08 | 1.701 | nucleus | K13044 | R | KOG0118 | FOG: RRM domain |
| Q16643 | DBN1 | 71.428 | 74.211 | 17.9 | 8 | 9 | 8 | 76215000 | 3.73E+08 | 1.688 | nucleus | | | | |
| Q9Y2W2 | WBP11 | 69.997 | 18.613 | 9.4 | 3 | 3 | 3 | 42675000 | 2.73E+08 | 1.667 | nucleus | K12866 | S | KOG4672 | Uncharacterized conserved low complexity protein |
| Q13151 | HNRNPA0 | 30.84 | 15.14 | 7.5 | 2 | 2 | 2 | 17107000 | 57176000 | 1.652 | nucleus | K12894 | | | |
| Q00610 | CLTC | 191.61 | 323.31 | 41.7 | 47 | 61 | 47 | 9.43E+08 | 3.22E+09 | 1.651 | cytoplasm | K04646 | U | KOG0985 | Vesicle coat protein clathrin, heavy chain |
| O95777 | LSM8 | 10.403 | 8.5156 | 16.7 | 1 | 3 | 1 | 20164000 | 73889000 | 1.65 | cytoplasm | K12627 | A | KOG1784 | Small Nuclear ribonucleoprotein splicing factor |
| Q99623 | PHB2 | 33.296 | 43.561 | 22.7 | 5 | 6 | 5 | 36891000 | 2.03E+08 | 1.65 | cytoplasm | K17081 | O | KOG3090 | Prohibitin-like protein |
| P10606 | COX5B | 13.696 | 22.098 | 24.8 | 3 | 3 | 3 | 22296000 | 83363000 | 1.646 | mitochondria | K02265 | C | KOG3352 | Cytochrome c oxidase, subunit Vb/COX4 |
| Q9P0L0 | VAPA | 27.893 | 11.718 | 13.7 | 2 | 2 | 2 | 20983000 | 78310000 | 1.63 | cytoplasm | K06096 | | | |
| Q9NX63 | CHCHD3 | 26.152 | 23.415 | 14.5 | 3 | 4 | 3 | 25626000 | 93713000 | 1.629 | extracellular | K17563 | K | KOG4083 | Head-elevated expression protein |
| P60842 | EIF4A1 | 46.153 | 12.492 | 9.6 | 3 | 3 | 2 | 11136000 | 24842000 | 1.626 | nucleus | K03257 | J | KOG0327 | Translation initiation factor 4F, helicase subunit (eIF-4A) and related helicases |
| P00558 | PGK1 | 44.614 | 82.972 | 44.1 | 10 | 13 | 10 | 1.41E+08 | 4.67E+08 | 1.607 | cytoplasm | K00927 | G | KOG1367 | 3-phosphoglycerate kinase |
| P47756 | CAPZB | 31.35 | 68.002 | 27.4 | 6 | 10 | 6 | 3.27E+08 | 3.15E+08 | 1.589 | cytoplasm,nucleus | K10365 | Z | KOG3174 | F-actin capping protein, beta subunit |
| P61978 | HNRNPK | 50.976 | 121.18 | 38.7 | 13 | 16 | 13 | 3.14E+08 | 1.62E+09 | 1.581 | nucleus | K12886 | AR | KOG2192 | PolyC-binding hnRNP-K protein HRB57A/hnRNP, contains KH domain |
| P08758 | ANXA5 | 35.936 | 53.781 | 22.5 | 6 | 7 | 6 | 77955000 | 2.5E+08 | 1.577 | cytoplasm | K16646 | U | KOG0819 | Annexin |
| P61158 | ACTR3 | 47.371 | 53.274 | 30.9 | 8 | 10 | 8 | 59337000 | 2.99E+08 | 1.576 | cytoplasm | K18584 | Z | KOG0678 | Actin-related protein Arp2/3 complex, subunit Arp3 |
| P43490 | NAMPT | 55.52 | 104.47 | 38.5 | 11 | 14 | 11 | 2.54E+08 | 9.12E+08 | 1.567 | cytoplasm | K03462 | | | |
| Q9Y3E1 | HDGFL3 | 22.619 | 22.515 | 20.7 | 3 | 3 | 2 | 35758000 | 2.2E+08 | 1.56 | nucleus | | | | |
| P38117 | ETFB | 27.843 | 21.789 | 17.6 | 4 | 4 | 4 | 37310000 | 53308000 | 1.559 | cytoplasm | K03521 | C | KOG3180 | Electron transfer flavoprotein, beta subunit |

| | | | | | | | | | | | | | | | |
|---|---|---|---|---|---|---|---|---|---|---|---|---|---|---|---|
| P52272 | HNRNPM | 77.515 | 48.747 | 12.3 | 6 | 6 | 6 | 63160000 | 1.89E+08 | 1.556 | cytoplasm | K12887 | A | KOG4212 | RNA-binding protein hnRNP-M |
| Q06830 | PRDX1 | 22.11 | 85.37 | 60.3 | 10 | 12 | 10 | 3.35E+08 | 9.59E+08 | 1.555 | cytoplasm | K13279 | O | KOG0852 | Alkyl hydroperoxide reductase, thiol specific antioxidant and related enzymes |
| O15511 | ARPC5 | 16.32 | 21.356 | 43.7 | 3 | 5 | 3 | 63159000 | 2.99E+08 | 1.547 | cytoplasm | K05754 | Z | KOG3380 | Actin-related protein Arp2/3 complex, subunit ARPC5 |
| P60981 | DSTN | 18.506 | 11.586 | 17 | 2 | 3 | 2 | 27428000 | 88172000 | 1.54 | mitochondria | K10363 | Z | KOG1735 | Actin depolymerizing factor |
| P07355 | ANXA2 | 38.604 | 283.3 | 75.5 | 27 | 40 | 27 | 1.66E+09 | 9.76E+09 | 1.519 | cytoplasm | K17092 | U | KOG0819 | Annexin |
| P62306 | SNRPF | 9.7251 | 22.789 | 48.8 | 3 | 4 | 3 | 19331000 | 47349000 | 1.516 | cytoplasm | K11098 | A | KOG3482 | Small nuclear ribonucleoprotein (snRNP) SMF |
| P05114 | HMGN1 | 10.659 | 30.051 | 45 | 4 | 4 | 4 | 5544900 | 12703000 | 1.515 | nucleus | K11299 | | | |
| Q96QD9 | FYTTD1 | 35.818 | 19.718 | 12.6 | 3 | 3 | 3 | 16487000 | 78063000 | 1.511 | mitochondria | | | | |
| O60701 | UGDH | 55.023 | 46.08 | 19.6 | 7 | 9 | 7 | 1.13E+08 | 2.81E+08 | 1.498 | cytoplasm | K00012 | GT | KOG2666 | UDP-glucose/GDP-mannose dehydrogenase |
| P63104 | YWHAZ | 27.745 | 108.55 | 44.5 | 11 | 13 | 10 | 2.87E+08 | 1.1E+09 | 1.496 | cytoplasm | K16197 | O | KOG0841 | Multifunctional chaperone (14-3-3 family) |
| P08574 | CYC1 | 35.422 | 13.149 | 8.6 | 2 | 2 | 2 | 13849000 | 41576000 | 1.474 | mitochondria | K00413 | C | KOG3052 | Cytochrome c1 |
| P09651 | HNRNPA1 | 38.746 | 127.64 | 34.7 | 11 | 17 | 11 | 2.72E+08 | 9.67E+08 | 1.47 | nucleus | K12741 | R | KOG0118 | FOG: RRM domain |
| Q8IY81 | FTSJ3 | 96.557 | 22.156 | 7.6 | 3 | 3 | 3 | 22601000 | 1.14E+08 | 1.469 | nucleus | K14857 | AR | KOG1098 | Putative SAM-dependent rRNA methyltransferase SPB1 |
| P07339 | CTSD | 44.552 | 37.804 | 15 | 5 | 5 | 5 | 1.06E+08 | 2.59E+08 | 1.462 | extracellular | K01379 | O | KOG1339 | Aspartyl protease |
| P26641 | EEF1G | 50.118 | 11.457 | 5.5 | 2 | 2 | 2 | 10197000 | 23763000 | 1.456 | cytoplasm | K03233 | J | KOG1627 | Translation elongation factor EF-1 gamma |
| Q00839 | HNRNPU | 90.583 | 107.12 | 26.5 | 14 | 15 | 14 | 2.3E+08 | 8.32E+08 | 1.456 | nucleus | K12888 | | | |
| P68371 | TUBB4B | 49.83 | 135.95 | 44.5 | 14 | 23 | 2 | 4.94E+08 | 1.8E+09 | 1.453 | cytoplasm,nucleus | K07375 | Z | KOG1375 | Beta tubulin |
| P18859 | ATP5J | 12.587 | 19.801 | 44.4 | 3 | 3 | 3 | 24042000 | 94681000 | 1.45 | mitochondria | K02131 | C | KOG4634 | Mitochondrial F1F0-ATP synthase, subunit Cf6 (coupling factor 6) |
| Q9ULV4 | CORO1C | 53.248 | 45.91 | 20.9 | 6 | 7 | 6 | 1.12E+08 | 4.5E+08 | 1.443 | mitochondria | K13886 | Z | KOG0303 | Actin-binding protein Coronin, contains WD40 repeats |
| P61421 | ATP6V0D1 | 40.329 | 20.734 | 17.7 | 3 | 4 | 3 | 16904000 | 1.32E+08 | 1.438 | cytoskeleton | K02146 | C | KOG2957 | Vacuolar H+-ATPase V0 sector, subunit d |
| P22695 | UQCRC2 | 48.442 | 105.97 | 34.4 | 11 | 14 | 11 | 1.46E+08 | 3.94E+08 | 1.434 | mitochondria | K00415 | C | KOG2583 | Ubiquinol cytochrome c reductase, subunit QCR2 |
| P63096 | GNAI1 | 40.361 | 32.294 | 24.3 | 6 | 9 | 3 | 25292000 | 1.71E+08 | 1.434 | cytoplasm | K04630 | DT | KOG0082 | G-protein alpha subunit (small G protein superfamily) |
| P40926 | MDH2 | 35.503 | 65.044 | 37.6 | 8 | 8 | 8 | 1.01E+08 | 3.26E+08 | 1.431 | mitochondria | K00026 | C | KOG1494 | NAD-dependent malate dehydrogenase |
| Q9UJS0 | SLC25A13 | 74.175 | 43.396 | 12.4 | 6 | 7 | 6 | 36183000 | 1.48E+08 | 1.424 | cytoplasm | K15105 | C | KOG0751 | Mitochondrial aspartate/glutamate carrier protein Aralar/Citrin (contains EF-hand Ca2+-binding domains) |
| P0DP25 | CALM3 | 16.837 | 169.78 | 74.5 | 14 | 32 | 14 | 1.91E+09 | 9.01E+09 | 1.423 | cytoplasm,nucleus | K02183 | | | |
| O95758 | PTBP3 | 59.689 | 43.225 | 17.6 | 7 | 7 | 4 | 13066000 | 84000000 | 1.419 | cytoskeleton | K17844 | A | KOG1190 | Polypyrimidine tract-binding protein |
| P29401 | TKT | 67.877 | 137.46 | 46.4 | 15 | 20 | 15 | 4.37E+08 | 1.27E+09 | 1.412 | cytoplasm | K00615 | G | KOG0523 | Transketolase |
| Q15050 | RRS1 | 41.193 | 19.557 | 12.1 | 3 | 4 | 3 | 30183000 | 98726000 | 1.411 | nucleus | K14852 | J | KOG1765 | Regulator of ribosome synthesis |
| P82979 | SARNP | 23.671 | 38.586 | 22.4 | 4 | 4 | 4 | 1.22E+08 | 2E+08 | 1.409 | nucleus | K18732 | D | KOG4259 | Putative nucleic acid-binding protein Hcc-1/proliferation associated |
| P05141 | SLC25A5 | 32.852 | 96.017 | 39.9 | 13 | 16 | 6 | 2.59E+08 | 8.99E+08 | 1.396 | cytoplasm | K05863 | C | KOG0749 | Mitochondrial ADP/ATP carrier proteins |
| P62241 | RPS8 | 24.205 | 57.806 | 42.3 | 7 | 8 | 7 | 1.05E+08 | 2.9E+08 | 1.396 | nucleus | K02995 | J | KOG3283 | 40S ribosomal protein S8 |
| P10599 | TXN | 11.737 | 29.269 | 41 | 4 | 5 | 4 | 90494000 | 4.21E+08 | 1.38 | extracellular | K03671 | O | KOG0907 | Thioredoxin |
| Q10588 | BST1 | 35.724 | 16.501 | 6.9 | 2 | 2 | 2 | 17279000 | 61227000 | 1.378 | extracellular | K18152 | | | |
| P08579 | SNRPB2 | 25.486 | 37.056 | 28 | 5 | 5 | 3 | 81171000 | 1.05E+08 | 1.377 | nucleus | K11094 | A | KOG4206 | Spliceosomal protein snRNP-U1A/U2B |
| Q9Y4Z0 | LSM4 | 15.35 | 14.511 | 11.5 | 2 | 2 | 2 | 23048000 | 50035000 | 1.375 | cytoplasm,nucleus | K12623 | A | KOG3293 | Small nuclear ribonucleoprotein (snRNP) |
| P16070 | CD44 | 81.537 | 27.406 | 5.1 | 3 | 3 | 3 | 65026000 | 1.7E+08 | 1.368 | extracellular,plasma r | K06256 | | | |
| P37837 | TALDO1 | 37.54 | 11.011 | 7.4 | 2 | 2 | 2 | 10443000 | 34823000 | 1.367 | mitochondria | K00616 | G | KOG2772 | Transaldolase |
| P62308 | SNRPG | 8.496 | 13.777 | 25 | 2 | 2 | 2 | 27941000 | 78527000 | 1.355 | cytoplasm | K11099 | A | KOG1780 | Small Nuclear ribonucleoprotein G |
| O95831 | AIFM1 | 66.9 | 20.785 | 8.8 | 3 | 3 | 3 | 11579000 | 46268000 | 1.338 | mitochondria | K04727 | T | KOG1346 | Programmed cell death 8 (apoptosis-inducing factor) |
| Q15365 | PCBP1 | 37.497 | 15.178 | 10.7 | 3 | 4 | 2 | 27243000 | 63739000 | 1.337 | cytoskeleton | K12889 | AR | KOG2190 | PolyC-binding proteins alphaCP-1 and related KH domain proteins |
| P15144 | ANPEP | 109.54 | 22.262 | 7.1 | 3 | 3 | 3 | 27947000 | 82277000 | 1.328 | endoplasmic reticulu | K11140 | EO | KOG1046 | Puromycin-sensitive aminopeptidase and related aminopeptidases |
| Q9Y3Y2 | CHTOP | 26.396 | 60.395 | 29.8 | 6 | 7 | 6 | 1.61E+08 | 4.72E+08 | 1.327 | mitochondria | | | | |
| P12236 | SLC25A6 | 32.866 | 41.928 | 43.6 | 12 | 16 | 4 | 1.33E+08 | 4.68E+08 | 1.326 | cytoplasm | K05863 | C | KOG0749 | Mitochondrial ADP/ATP carrier proteins |
| Q12965 | MYO1E | 127.06 | 75.718 | 13.2 | 10 | 10 | 10 | 1.04E+08 | 3.35E+08 | 1.325 | cytoplasm | K10356 | Z | KOG0164 | Myosin class I heavy chain |
| P60953 | CDC42 | 21.258 | 33.308 | 34.6 | 4 | 4 | 4 | 44736000 | 2.53E+08 | 1.319 | cytoplasm | K04393 | R | KOG0393 | Ras-related small GTPase, Rho type |
| P23284 | PPIB | 23.742 | 59.35 | 34.3 | 7 | 10 | 7 | 99955000 | 3.67E+08 | 1.315 | extracellular | K03768 | O | KOG0880 | Peptidyl-prolyl cis-trans isomerase |
| P83916 | CBX1 | 21.418 | 24.182 | 24.9 | 4 | 5 | 3 | 21373000 | 46752000 | 1.298 | nucleus | K11585 | | | |
| P05556 | ITGB1 | 88.414 | 51.146 | 15.3 | 7 | 8 | 7 | 82078000 | 2.87E+08 | 1.295 | extracellular | K05719 | TW | KOG1226 | Integrin beta subunit (N-terminal portion of extracellular region) |
| P00441 | SOD1 | 15.936 | 43.508 | 53.9 | 3 | 4 | 3 | 1.69E+08 | 6.67E+08 | 1.292 | cytoplasm | K04565 | P | KOG0441 | Cu2+/Zn2+ superoxide dismutase SOD1 |
| P11021 | HSPA5 | 72.332 | 100.64 | 29.8 | 13 | 14 | 13 | 1.41E+08 | 3.36E+08 | 1.291 | endoplasmic reticulu | K09490 | O | KOG0101 | Molecular chaperones HSP70/HSC70, HSP70 superfamily |
| P59998 | ARPC4 | 19.667 | 49.465 | 39.9 | 6 | 7 | 6 | 86038000 | 2.39E+08 | 1.286 | mitochondria | K05755 | Z | KOG1876 | Actin-related protein Arp2/3 complex, subunit ARPC4 |

| | | | | | | | | | | | | | | |
|---|---|---|---|---|---|---|---|---|---|---|---|---|---|---|
| P62820 | RAB1A | 22.677 | 21.346 | 22.9 | 3 | 3 | 2 | 49001000 | 1.12E+08 | 1.285 | cytoplasm | K07874 | TU | KOG0084 GTPase Rab1/YPT1, small G protein superfamily |
| P30043 | BLVRB | 22.119 | 22.664 | 25.7 | 3 | 3 | 3 | 19638000 | 72070000 | 1.284 | cytoplasm | K05901 | | |
| P60174 | TPI1 | 30.791 | 82.503 | 48.3 | 9 | 12 | 9 | 2.63E+08 | 7.77E+08 | 1.284 | cytoplasm | K01803 | G | KOG1643 Triosephosphate isomerase |
| P49755 | TMED10 | 24.976 | 17.095 | 11.9 | 2 | 3 | 2 | 65658000 | 1.34E+08 | 1.278 | extracellular | K20352 | U | KOG1691 emp24/gp25L/p24 family of membrane trafficking proteins |
| Q04837 | SSBP1 | 17.259 | 32.785 | 32.4 | 3 | 4 | 3 | 36110000 | 1.42E+08 | 1.278 | nucleus | K03111 | L | KOG1653 Single-stranded DNA-binding protein |
| P27348 | YWHAQ | 27.764 | 45.7 | 30.6 | 6 | 7 | 5 | 68082000 | 2.87E+08 | 1.277 | cytoplasm | K16197 | O | KOG0841 Multifunctional chaperone (14-3-3 family) |
| P20674 | COX5A | 16.762 | 30.699 | 26 | 4 | 9 | 4 | 1.34E+08 | 4.27E+08 | 1.267 | mitochondria | K02264 | C | KOG4077 Cytochrome c oxidase, subunit Va/COX6 |
| Q9NZI8 | IGF2BP1 | 63.48 | 34.163 | 10.6 | 4 | 5 | 3 | 46592000 | 1.15E+08 | 1.26 | cytoplasm | K17391 | AR | KOG2193 IGF-II mRNA-binding protein IMP, contains RRM and KH domains |
| O15145 | ARPC3 | 20.546 | 14.718 | 20.8 | 2 | 2 | 2 | 31039000 | 86242000 | 1.254 | cytoplasm | K05756 | Z | KOG3155 Actin-related protein Arp2/3 complex, subunit ARPC3 |
| Q99536 | VAT1 | 41.92 | 29.445 | 16 | 4 | 4 | 4 | 12017000 | 73587000 | 1.249 | cytoplasm | | CR | KOG1198 Zinc-binding oxidoreductase |
| Q9Y3A2 | UTP11 | 30.446 | 15.097 | 13.4 | 2 | 2 | 2 | 19442000 | 60785000 | 1.249 | nucleus | K14769 | S | KOG3237 Uncharacterized conserved protein |
| P04075 | ALDOA | 39.42 | 82.281 | 47.8 | 10 | 13 | 8 | 1.99E+08 | 4.83E+08 | 1.239 | cytoplasm | K01623 | G | KOG1557 Fructose-biphosphate aldolase |
| P04844 | RPN2 | 69.283 | 71.392 | 29 | 8 | 9 | 8 | 58636000 | 5.4E+08 | 1.234 | extracellular | K12667 | O | KOG2447 Oligosaccharyltransferase, delta subunit (ribophorin II) |
| P22392 | NME2 | 17.298 | 48.122 | 48 | 5 | 7 | 3 | 74551000 | 1.54E+08 | 1.234 | cytoplasm | K00940 | F | KOG0888 Nucleoside diphosphate kinase |
| P52907 | CAPZA1 | 32.922 | 50.431 | 28 | 5 | 6 | 4 | 73449000 | 3.38E+08 | 1.234 | cytoplasm | K10364 | Z | KOG0836 F-actin capping protein, alpha subunit |
| P61981 | YWHAG | 28.302 | 49.691 | 42.9 | 8 | 10 | 6 | 72331000 | 5.13E+08 | 1.225 | cytoplasm | K16198 | O | KOG0841 Multifunctional chaperone (14-3-3 family) |
| Q15084 | PDIA6 | 48.121 | 24.755 | 14.1 | 3 | 3 | 3 | 12260000 | 38116000 | 1.225 | extracellular | K09584 | O | KOG0191 Thioredoxin/protein disulfide isomerase |
| P68363 | TUBA1B | 50.151 | 175.57 | 62.7 | 19 | 29 | 0 | 8.13E+08 | 2.85E+09 | 1.221 | cytoskeleton | K07374 | Z | KOG1376 Alpha tubulin |
| P31943 | HNRNPH1 | 49.229 | 32.803 | 14.9 | 4 | 4 | 4 | 28193000 | 78899000 | 1.209 | nucleus | K12898 | A | KOG4211 Splicing factor hnRNP-F and related RNA-binding proteins |
| P68032 | ACTC1 | 42.019 | 43.304 | 44.8 | 19 | 50 | 3 | 99415000 | 4.12E+08 | 1.194 | cytoskeleton | K12314 | Z | KOG0676 Actin and related proteins |
| P38159 | RBMX | 42.331 | 37.335 | 15.9 | 5 | 6 | 5 | 65070000 | 2.34E+08 | 1.182 | nucleus | K12885 | | |
| Q15428 | SF3A2 | 49.255 | 28.014 | 9.5 | 3 | 4 | 3 | 46945000 | 1.42E+08 | 1.175 | cytoplasm | K12826 | A | KOG0227 Splicing factor 3a, subunit 2 |
| Q14978 | NOLC1 | 73.602 | 81.159 | 15.5 | 10 | 18 | 10 | 5.2E+08 | 1.16E+09 | 1.173 | nucleus | | Y | KOG2992 Nucleolar GTPase/ATPase p130 |
| P62937 | PPIA | 18.012 | 86.457 | 58.2 | 9 | 14 | 9 | 3.04E+08 | 8.89E+08 | 1.169 | cytoplasm | K03767 | O | KOG0865 Cyclophilin type peptidyl-prolyl cis-trans isomerase |
| O00567 | NOP56 | 66.049 | 77.556 | 26.1 | 10 | 12 | 10 | 89590000 | 2.55E+08 | 1.166 | nucleus | K14564 | AJ | KOG2572 Ribosome biogenesis protein - Nop58p/Nop5p |
| P61604 | HSPE1 | 10.932 | 37.551 | 52 | 5 | 6 | 5 | 1.35E+08 | 3.03E+08 | 1.164 | mitochondria | K04078 | O | KOG1641 Mitochondrial chaperonin |
| Q14764 | MVP | 99.326 | 51.658 | 15.5 | 7 | 7 | 7 | 27365000 | 1.75E+08 | 1.16 | cytoplasm | K17266 | | |
| P07919 | UQCRH | 10.739 | 23.867 | 19.8 | 1 | 3 | 1 | 46412000 | 82857000 | 1.159 | extracellular | K00416 | C | KOG4763 Ubiquinol-cytochrome c reductase hinge protein |
| Q9NX24 | NHP2 | 17.201 | 49.201 | 68.6 | 5 | 7 | 5 | 2.37E+08 | 8.13E+08 | 1.155 | cytoplasm | K11129 | A | KOG3167 Box H/ACA snoRNP component, involved in ribosomal RNA pseudouridinylation |
| P62314 | SNRPD1 | 13.281 | 30.049 | 54.6 | 4 | 5 | 4 | 1.11E+08 | 2.86E+08 | 1.149 | nucleus | K11087 | A | KOG3428 Small nuclear ribonucleoprotein SMD1 and related snRNPs |
| Q9Y6C9 | MTCH2 | 33.331 | 47.901 | 32 | 6 | 8 | 6 | 1.71E+08 | 6.16E+08 | 1.147 | extracellular | K17885 | R | KOG2745 Mitochondrial carrier protein |
| P43243 | MATR3 | 94.622 | 55.917 | 12.9 | 6 | 6 | 6 | 1.63E+08 | 4.5E+08 | 1.144 | nucleus | K13213 | | |
| P38646 | HSPA9 | 73.68 | 131.03 | 29.5 | 15 | 16 | 15 | 2.34E+08 | 5.83E+08 | 1.136 | mitochondria | K04043 | O | KOG0102 Molecular chaperones mortalin/PBP74/GRP75, HSP70 superfamily |
| P62316 | SNRPD2 | 13.527 | 51.649 | 67.8 | 8 | 8 | 8 | 1.03E+08 | 2.66E+08 | 1.136 | cytoplasm | K11096 | A | KOG3459 Small nuclear ribonucleoprotein (snRNP) Sm core protein |
| P22626 | HNRNPA2B | 37.429 | 143.87 | 55.5 | 17 | 25 | 17 | 4.85E+08 | 1.54E+09 | 1.131 | nucleus | K13158 | R | KOG0118 FOG: RRM domain |
| P54289 | CACNA2D1 | 124.57 | 37.792 | 7.9 | 6 | 6 | 6 | 22916000 | 1.12E+08 | 1.121 | plasma membrane | K04859 | PT | KOG2353 L-type voltage-dependent Ca2+ channel, alpha2/delta subunit |
| P62879 | GNB2 | 37.331 | 11.35 | 19.1 | 4 | 6 | 1 | 21775000 | 68770000 | 1.121 | cytoplasm | K04537 | R | KOG0286 G-protein beta subunit |
| P53999 | SUB1 | 14.395 | 29.265 | 37 | 4 | 4 | 4 | 64254000 | 1.45E+08 | 1.12 | nucleus | | K | KOG2712 Transcriptional coactivator |
| P38919 | EIF4A3 | 46.871 | 33.937 | 18.2 | 5 | 5 | 4 | 57083000 | 1.68E+08 | 1.116 | nucleus | K13025 | J | KOG0327 Translation initiation factor 4F, helicase subunit (eIF-4A) and related helicases |
| P31949 | S100A11 | 11.74 | 19.958 | 45.7 | 2 | 3 | 2 | 1.51E+08 | 8.2E+08 | 1.114 | cytoplasm | | | |
| P22087 | FBL | 33.784 | 157.07 | 58.6 | 14 | 28 | 13 | 1.68E+09 | 6.15E+09 | 1.109 | nucleus | K14563 | A | KOG1596 Fibrillarin and related nucleolar RNA-binding proteins |
| O75955 | FLOT1 | 47.355 | 100.79 | 33.7 | 10 | 14 | 10 | 2.01E+08 | 4.76E+08 | 1.107 | cytoplasm | K07192 | UZ | KOG2668 Flotillins |
| P62753 | RPS6 | 28.68 | 40.198 | 22.9 | 5 | 5 | 5 | 61555000 | 1.08E+08 | 1.107 | cytoplasm | K02991 | J | KOG1646 40S ribosomal protein S6 |
| Q14254 | FLOT2 | 47.064 | 104.2 | 34.8 | 11 | 14 | 11 | 2.27E+08 | 6.18E+08 | 1.105 | cytoplasm | K07192 | UZ | KOG2668 Flotillins |
| Q9NS69 | TOMM22 | 15.521 | 42.084 | 66.2 | 5 | 6 | 5 | 88148000 | 1.97E+08 | 1.096 | cytoplasm | K17769 | U | KOG4111 Translocase of outer mitochondrial membrane complex, subunit TOM22 |
| Q5SSJ5 | HP1BP3 | 61.206 | 25.639 | 7.6 | 4 | 4 | 4 | 11907000 | 54544000 | 1.095 | nucleus | | B | KOG4012 Histone H1 |
| P04083 | ANXA1 | 38.714 | 175.13 | 60.4 | 17 | 24 | 17 | 6.82E+08 | 1.99E+09 | 1.091 | cytoplasm | K17091 | U | KOG0819 Annexin |
| P46087 | NOP2 | 89.301 | 23.223 | 4.7 | 3 | 4 | 3 | 26268000 | 59835000 | 1.09 | nucleus | K14835 | A | KOG1122 tRNA and rRNA cytosine-C5-methylase (nucleolar protein NOL1/NOP2) |
| Q02543 | RPL18A | 20.762 | 22.969 | 16.5 | 3 | 3 | 3 | 26262000 | 50653000 | 1.089 | nucleus | K02882 | J | KOG0829 60S ribosomal protein L18A |
| Q9H1E3 | NUCKS1 | 27.296 | 21.183 | 17.7 | 3 | 5 | 3 | 16198000 | 37669000 | 1.088 | nucleus | | | |
| P55769 | SNU13 | 14.173 | 78.825 | 68.8 | 9 | 12 | 9 | 5.35E+08 | 2.3E+09 | 1.084 | cytoplasm | K12845 | AJ | KOG3387 60S ribosomal protein 15.5kD/SNU13, NHP2/L7A family |

| | | | | | | | | | | | | | | | |
|---|---|---|---|---|---|---|---|---|---|---|---|---|---|---|---|
| P14406 | COX7A2 | 9.3959 | 16.479 | 27.7 | 2 | 2 | 2 | 67689000 | 2.08E+08 | 1.08 | mitochondria | K02270 | | | (includes ribonuclease P subunit p38), involved in splicing |
| P10809 | HSPD1 | 61.054 | 116.04 | 33 | 15 | 18 | 15 | 3.57E+08 | 7.7E+08 | 1.077 | mitochondria | K04077 | O | KOG0356 | Mitochondrial chaperonin, Cpn60/Hsp60p |
| P07195 | LDHB | 36.638 | 75.239 | 38.6 | 10 | 13 | 9 | 2.05E+08 | 5.79E+08 | 1.076 | cytoplasm | K00016 | C | KOG1495 | Lactate dehydrogenase |
| P07437 | TUBB | 49.67 | 42.713 | 41.2 | 13 | 21 | 3 | 1.08E+08 | 4.59E+08 | 1.072 | cytoplasm,nucleus | K07375 | Z | KOG1375 | Beta tubulin |
| P14625 | HSP90B1 | 92.468 | 50.801 | 16.4 | 8 | 8 | 7 | 53330000 | 2.29E+08 | 1.07 | endoplasmic reticulu | K09487 | O | KOG0020 | Endoplasmic reticulum glucose-regulated protein (GRP94/endoplasmin) |
| Q96AE4 | FUBP1 | 67.56 | 23.382 | 5.3 | 3 | 4 | 3 | 16929000 | 32690000 | 1.069 | cytoplasm | K13210 | A | KOG1676 | K-homology type RNA binding proteins |
| Q12906 | ILF3 | 95.337 | 104.35 | 23.2 | 12 | 15 | 12 | 3.75E+08 | 9.15E+08 | 1.066 | nucleus | K13090 | R | KOG3792 | Transcription factor NFAT, subunit NF90 |
| P14854 | COX6B1 | 10.192 | 21.178 | 47.7 | 3 | 4 | 3 | 70423000 | 2.04E+08 | 1.058 | extracellular | K02267 | C | KOG3057 | Cytochrome c oxidase, subunit VIb/COX12 |
| Q05639 | EEF1A2 | 50.47 | 15.094 | 41 | 11 | 16 | 2 | 56349000 | 1.9E+08 | 1.048 | cytoplasm | K03231 | J | KOG0052 | Translation elongation factor EF-1 alpha/Tu |
| P24534 | EEF1B2 | 24.763 | 12.633 | 22.2 | 2 | 2 | 2 | 15912000 | 39326000 | 1.046 | nucleus | K03232 | K | KOG1668 | Elongation factor 1 beta/delta chain |
| P84090 | ERH | 12.259 | 32.627 | 64.4 | 4 | 4 | 4 | 1.13E+08 | 3.98E+08 | 1.043 | cytoplasm | | R | KOG1766 | Enhancer of rudimentary |
| P62826 | RAN | 24.423 | 79.21 | 43.1 | 9 | 12 | 9 | 2.2E+08 | 6.14E+08 | 1.041 | cytoplasm | K07936 | U | KOG0096 | GTPase Ran/TC4/GSP1 (nuclear protein transport pathway), small G protein |
| Q92522 | H1FX | 22.487 | 5.5407 | 5.6 | 1 | 2 | 1 | 1.28E+08 | 2.47E+09 | 1.034 | nucleus | K11275 | B | KOG4012 | Histone H1 |
| P62258 | YWHAE | 29.174 | 56.839 | 35.3 | 8 | 8 | 7 | 1.07E+08 | 3.27E+08 | 1.033 | cytoplasm | K06630 | O | KOG0841 | Multifunctional chaperone (14-3-3 family) |
| P11142 | HSPA8 | 70.897 | 177.54 | 42.1 | 19 | 25 | 14 | 9.59E+08 | 1.8E+09 | 1.031 | cytoplasm | K03283 | O | KOG0101 | Molecular chaperones HSP70/HSC70, HSP70 superfamily |
| Q9Y2R4 | DDX52 | 67.497 | 12.764 | 4.5 | 2 | 4 | 2 | 18669000 | 30965000 | 1.028 | nucleus | K14779 | A | KOG0344 | ATP-dependent RNA helicase |
| Q8N3V7 | SYNPO | 99.462 | 22.52 | 5 | 3 | 3 | 3 | 20215000 | 43332000 | 1.024 | nucleus | K21112 | | | |
| Q9BZE4 | GTPBP4 | 73.964 | 11.716 | 5.5 | 2 | 2 | 2 | 20259000 | 43118000 | 1.022 | cytoplasm | K06943 | R | KOG1490 | GTP-binding protein CRFG/NOG1 (ODN superfamily) |
| P00338 | LDHA | 36.688 | 87.39 | 58.7 | 12 | 13 | 11 | 2.07E+08 | 4.46E+08 | 1.014 | cytoplasm | K00016 | C | KOG1495 | Lactate dehydrogenase |
| Q969Q0 | RPL36AL | 12.469 | 12.836 | 16 | 2 | 2 | 2 | 28577000 | 49748000 | 1.012 | extracellular | K02929 | J | KOG3464 | 60S ribosomal protein L44 |
| P54727 | RAD23B | 43.171 | 11.76 | 5.9 | 2 | 2 | 2 | 14483000 | 21159000 | 1.01 | cytoplasm | K10839 | L | KOG0011 | Nucleotide excision repair factor NEF2, RAD23 component |
| P67809 | YBX1 | 35.924 | 31.287 | 20.4 | 3 | 3 | 2 | 49915000 | 77928000 | 1.007 | nucleus | K09276 | J | KOG3070 | Predicted RNA-binding protein containing PIN domain |
| P13073 | COX4I1 | 19.576 | 32.266 | 26 | 4 | 4 | 4 | 1.28E+08 | 3.7E+08 | 1.003 | mitochondria | K02263 | C | KOG4075 | Cytochrome c oxidase, subunit IV/COX5b |
| Q96A72 | MAGOHB | 17.276 | 23.805 | 29.7 | 3 | 4 | 3 | 94435000 | 87526000 | 1 | cytoplasm | K12877 | A | KOG3392 | Exon-exon junction complex, Magoh component |
| O14950 | MYL12B | 19.779 | 5.634 | 83.7 | 15 | 51 | 1 | 1.73E+08 | 1.29E+09 | 0.999 | mitochondria | K12757 | Z | KOG0031 | Myosin regulatory light chain, EF-Hand protein superfamily |
| P46782 | RPS5 | 22.876 | 25.987 | 30.9 | 3 | 6 | 3 | 1.28E+08 | 3.48E+08 | 0.992 | cytoplasm | K02989 | J | KOG3291 | Ribosomal protein S7 |
| Q15061 | WDR43 | 74.89 | 11.627 | 5.3 | 2 | 2 | 2 | 18118000 | 51542000 | 0.983 | cytoplasm | K14546 | R | KOG4547 | WD40 repeat-containing protein |
| P62701 | RPS4X | 29.597 | 72.339 | 37.3 | 9 | 10 | 9 | 2.4E+08 | 4.93E+08 | 0.968 | cytoplasm | K02987 | J | KOG0378 | 40S ribosomal protein S4 |
| P51858 | HDGF | 26.788 | 101.39 | 51.2 | 10 | 14 | 10 | 6.04E+08 | 9.68E+08 | 0.962 | nucleus | K16641 | | | |
| Q1KMD3 | HNRNPUL2 | 85.104 | 45.15 | 15.4 | 6 | 6 | 6 | 2.61E+09 | 1.86E+08 | 0.957 | nucleus | | | | |
| P36873 | PPP1CC | 36.983 | 36.626 | 23.2 | 5 | 5 | 5 | 22655000 | 87564000 | 0.954 | cytoplasm | K06269 | | | |
| O75569 | PRKRA | 34.404 | 31.566 | 18.2 | 4 | 4 | 4 | 56338000 | 1.37E+08 | 0.953 | cytoplasm | | UK | KOG3732 | Staufen and related double-stranded-RNA-binding proteins |
| P08195 | SLC3A2 | 67.993 | 42.579 | 13.3 | 5 | 5 | 5 | 64992000 | 1.09E+08 | 0.951 | endoplasmic reticulu | K06519 | G | KOG0471 | Alpha-amylase |
| P14678 | SNRPB | 24.61 | 49.941 | 21.7 | 7 | 9 | 7 | 1.55E+08 | 3.59E+08 | 0.95 | cytoplasm | K11086 | K | KOG3168 | U1 snRNP component |
| P09382 | LGALS1 | 14.716 | 65.318 | 60.7 | 7 | 9 | 7 | 2.77E+08 | 9.05E+08 | 0.947 | extracellular | K06830 | W | KOG3587 | Galectin, galactose-binding lectin |
| P36578 | RPL4 | 47.697 | 49.215 | 20.6 | 6 | 7 | 6 | 2.33E+08 | 5.65E+08 | 0.944 | cytoplasm | K02930 | A | KOG1475 | Ribosomal protein RPL1/RPL2/RL4L4 |
| P18669 | PGAM1 | 28.804 | 29.875 | 32.7 | 4 | 5 | 4 | 1.25E+08 | 2.47E+08 | 0.941 | cytoplasm | K01834 | G | KOG0235 | Phosphoglycerate mutase |
| P23528 | CFL1 | 18.502 | 92.828 | 71.1 | 11 | 16 | 8 | 6.03E+08 | 1.75E+09 | 0.941 | mitochondria | K05765 | Z | KOG1735 | Actin depolymerizing factor |
| P30050 | RPL12 | 17.818 | 56.55 | 49.7 | 6 | 6 | 6 | 1.31E+08 | 3.33E+08 | 0.935 | cytoplasm | K02870 | J | KOG0886 | 40S ribosomal protein S2 |
| P62910 | RPL32 | 15.86 | 32.515 | 33.3 | 4 | 6 | 4 | 41599000 | 70131000 | 0.935 | cytoplasm | K02912 | J | KOG0878 | 60S ribosomal protein L32 |
| P41208 | CETN2 | 19.738 | 15.793 | 17.4 | 2 | 3 | 2 | 22533000 | 47340000 | 0.933 | nucleus | K10840 | ZD | KOG0028 | Ca2+-binding protein (centrin/caltractin), EF-Hand superfamily protein |
| Q99848 | EBNA1BP2 | 34.852 | 44.547 | 22.2 | 5 | 8 | 5 | 1.56E+08 | 3.33E+08 | 0.933 | nucleus | K14823 | A | KOG3080 | Nucleolar protein-like/EBNA1-binding protein |
| Q9Y277 | VDAC3 | 30.658 | 84.84 | 45.2 | 9 | 13 | 9 | 1.88E+08 | 4.46E+08 | 0.933 | cytoplasm | K15041 | P | KOG3126 | Porin/voltage-dependent anion-selective channel protein |
| Q9NY12 | GAR1 | 22.348 | 39.287 | 27.2 | 6 | 7 | 6 | 2.28E+08 | 6.74E+08 | 0.93 | cytoplasm | K11128 | J | KOG3262 | H/ACA small nucleolar RNP component GAR1 |
| Q9UJZ1 | STOML2 | 38.534 | 79.115 | 37.1 | 8 | 12 | 8 | 1.66E+08 | 3.87E+08 | 0.93 | mitochondria | | C | KOG2620 | Prohibitins and stomatins of the PID superfamily |
| P62913 | RPL11 | 20.252 | 28.274 | 19.7 | 3 | 3 | 3 | 58378000 | 1.04E+08 | 0.924 | cytoplasm | K02868 | J | KOG0397 | 60S ribosomal protein L11 |
| P18077 | RPL35A | 12.538 | 12.206 | 14.5 | 2 | 2 | 2 | 30936000 | 45372000 | 0.921 | cytoplasm | K02917 | J | KOG0887 | 60S ribosomal protein L35A/L37 |
| P27635 | RPL10 | 24.604 | 20.039 | 23.8 | 3 | 4 | 3 | 60694000 | 1.39E+08 | 0.92 | cytoplasm | K02866 | J | KOG0857 | 60s ribosomal protein L10 |
| O43169 | CYB5B | 16.332 | 16.541 | 23.3 | 1 | 2 | 1 | 69509000 | 2.15E+08 | 0.917 | peroxisome | | C | KOG0537 | Cytochrome b5 |
| P11940 | PABPC1 | 70.67 | 15.494 | 6.3 | 2 | 2 | 2 | 24395000 | 44159000 | 0.917 | nucleus | K13126 | AJ | KOG0123 | Polyadenylate-binding protein (RRM superfamily) |
| O43707 | ACTN4 | 104.85 | 323.31 | 71.2 | 48 | 80 | 33 | 3.96E+09 | 1.17E+10 | 0.915 | cytoplasm | K05699 | Z | KOG0035 | Ca2+-binding actin-bundling protein (actinin), alpha chain |

| | | | | | | | | | | | | | | |
|---|---|---|---|---|---|---|---|---|---|---|---|---|---|---|
| P50395 | GDI2 | 50.663 | 15.426 | 4.5 | 2 | 3 | 2 | 53585000 | 88082000 | 0.913 | endoplasmic reticulu | K17255 | O | KOG1439 RAB proteins geranylgeranyltransferase component A (RAB escort protein) |
| P52209 | PGD | 53.139 | 31.17 | 13.7 | 5 | 6 | 5 | 53902000 | 1.39E+08 | 0.913 | cytoplasm,nucleus | K00033 | G | KOG2653 6-phosphogluconate dehydrogenase |
| Q16891 | IMMT | 83.677 | 211.6 | 43.3 | 24 | 32 | 24 | 8.46E+08 | 1.83E+09 | 0.903 | mitochondria | K17785 | M | KOG1854 Mitochondrial inner membrane protein (mitofilin) |
| Q9Y512 | SAMM50 | 51.976 | 51.727 | 16.6 | 6 | 6 | 6 | 1.05E+08 | 1.59E+08 | 0.898 | cytoplasm | K07277 | R | KOG2602 Predicted cell surface protein homologous to bacterial outer membrane proteins |
| P17676 | CEBPB | 36.105 | 38.469 | 19.1 | 4 | 5 | 4 | 35107000 | 74370000 | 0.895 | nucleus | K10048 | | |
| P62318 | SNRPD3 | 13.916 | 51.628 | 53.2 | 5 | 8 | 5 | 2.81E+08 | 6.62E+08 | 0.892 | cytoplasm | K11088 | A | KOG3172 Small nuclear ribonucleoprotein Sm D3 |
| Q9NR30 | DDX21 | 87.343 | 11.949 | 4.2 | 2 | 2 | 2 | 17984000 | 41203000 | 0.888 | cytoplasm,nucleus | K16911 | A | KOG0331 ATP-dependent RNA helicase |
| P13639 | EEF2 | 95.337 | 39.816 | 13.6 | 6 | 7 | 6 | 46310000 | 97633000 | 0.886 | cytoplasm | K03234 | J | KOG0469 Elongation factor 2 |
| P46783 | RPS10 | 18.898 | 48.493 | 38.2 | 5 | 10 | 5 | 2.29E+08 | 2.15E+09 | 0.886 | nucleus | K02947 | J | KOG3344 40s ribosomal protein s10 |
| P18754 | RCC1 | 44.969 | 115.39 | 50.8 | 11 | 14 | 11 | 3.86E+08 | 8.69E+08 | 0.882 | nucleus | K11493 | | |
| Q9NVP1 | DDX18 | 75.406 | 24.057 | 6.3 | 3 | 3 | 3 | 45458000 | 76925000 | 0.879 | nucleus | K13179 | A | KOG0342 ATP-dependent RNA helicase pitchoune |
| Q15287 | RNPS1 | 34.208 | 48.054 | 23.3 | 5 | 8 | 5 | 1.43E+08 | 3.32E+08 | 0.875 | nucleus | K14325 | A | KOG4209 Splicing factor RNPS1, SR protein superfamily |
| P25398 | RPS12 | 14.515 | 25.109 | 34.1 | 3 | 5 | 3 | 49025000 | 71815000 | 0.874 | cytoplasm | K02951 | J | KOG3406 40S ribosomal protein S12 |
| P40429 | RPL13A | 23.577 | 19.102 | 14.8 | 3 | 3 | 3 | 29247000 | 57904000 | 0.869 | cytoplasm | K02872 | J | KOG3204 60S ribosomal protein L13a |
| P14618 | PKM | 57.936 | 294.48 | 72.5 | 27 | 40 | 27 | 1.68E+09 | 4.14E+09 | 0.867 | cytoplasm | K00873 | G | KOG2323 Pyruvate kinase |
| P55072 | VCP | 89.321 | 27.984 | 7.4 | 4 | 5 | 4 | 46812000 | 81607000 | 0.847 | cytoplasm,nucleus | K13525 | O | KOG0730 AAA+-type ATPase |
| P06733 | ENO1 | 47.168 | 186.92 | 53.5 | 15 | 22 | 15 | 5.24E+08 | 1.22E+09 | 0.845 | cytoplasm | K01689 | G | KOG2670 Enolase |
| P07237 | P4HB | 57.116 | 21.837 | 8.5 | 4 | 4 | 4 | 29359000 | 83374000 | 0.838 | extracellular | K09580 | O | KOG0190 Protein disulfide isomerase (prolyl 4-hydroxylase beta subunit) |
| O75367 | H2AFY | 39.617 | 183.62 | 68.8 | 17 | 28 | 15 | 9.41E+08 | 2.56E+09 | 0.836 | mitochondria | K11251 | BK | KOG2633 Hismacro and SEC14 domain-containing proteins |
| Q00688 | FKBP3 | 25.177 | 23.618 | 16.5 | 3 | 3 | 3 | 14188000 | 25173000 | 0.836 | cytoplasm | K09570 | O | KOG0544 FKBP-type peptidyl-prolyl cis-trans isomerase |
| P63261 | ACTG1 | 41.792 | 323.31 | 95.5 | 38 | 130 | 2 | 1.18E+10 | 6.32E+10 | 0.834 | cytoskeleton | K05692 | Z | KOG0676 Actin and related proteins |
| P09972 | ALDOC | 39.455 | 18.737 | 16.8 | 4 | 6 | 2 | 86404000 | 2.53E+08 | 0.832 | cytoplasm | K01623 | G | KOG1557 Fructose-biphosphate aldolase |
| Q96C19 | EFHD2 | 26.697 | 84.193 | 43.3 | 11 | 12 | 11 | 3.51E+08 | 9.06E+08 | 0.829 | cytoplasm | | R | KOG0041 Predicted Ca2+-binding protein, EF-Hand protein superfamily |
| P31930 | UQCRC1 | 52.645 | 37.454 | 12.5 | 4 | 5 | 4 | 82151000 | 3.28E+08 | 0.821 | mitochondria | K00414 | O | KOG0960 Mitochondrial processing peptidase, beta subunit, and related enzymes |
| P07900 | HSP90AA1 | 84.659 | 35.3 | 16 | 10 | 13 | 4 | 41650000 | 78295000 | 0.82 | cytoplasm | K04079 | O | KOG0020 Endoplasmic reticulum glucose-regulated protein (GRP94/endoplasmin) |
| Q6DKI1 | RPL7L1 | 28.661 | 38.509 | 31.3 | 6 | 6 | 6 | 1.08E+08 | 2.08E+08 | 0.815 | cytoplasm | | J | KOG3184 60S ribosomal protein L7 |
| P08238 | HSP90AB1 | 83.263 | 106.07 | 25 | 14 | 16 | 6 | 2.42E+08 | 3.57E+08 | 0.814 | cytoplasm | K04079 | O | KOG0020 Endoplasmic reticulum glucose-regulated protein (GRP94/endoplasmin) |
| Q99729 | HNRNPAB | 36.224 | 22.316 | 12.7 | 3 | 4 | 3 | 61885000 | 99963000 | 0.812 | nucleus | K13044 | R | KOG0118 FOG: RRM domain |
| P28331 | NDUFS1 | 79.467 | 36.535 | 13.6 | 5 | 5 | 5 | 42678000 | 1.37E+08 | 0.81 | mitochondria | K03934 | C | KOG2282 NADH-ubiquinone oxidoreductase, NDUFS1/75 kDa subunit |
| Q15149 | PLEC | 531.78 | 323.31 | 29.7 | 100 | 118 | 100 | 2.25E+09 | 4.12E+09 | 0.807 | cytoplasm,nucleus | K10388 | J | KOG3344 40s ribosomal protein s10 |
| P56182 | RRP1 | 52.839 | 26.155 | 13.7 | 4 | 4 | 4 | 38706000 | 92684000 | 0.804 | nucleus | K14849 | A | KOG3911 Nucleolar protein NOP52/RRP1 |
| Q00059 | TFAM | 29.096 | 66.881 | 30.9 | 8 | 13 | 8 | 5.17E+08 | 8.46E+08 | 0.798 | mitochondria | K11830 | R | KOG0381 HMG box-containing protein |
| Q15427 | SF3B4 | 44.385 | 17.566 | 11.1 | 2 | 3 | 2 | 1.01E+08 | 2.84E+08 | 0.795 | nucleus | K12831 | A | KOG0131 Splicing factor 3b, subunit 4 |
| Q86V81 | ALYREF | 26.888 | 108.35 | 49.4 | 11 | 15 | 11 | 6.28E+08 | 1.66E+09 | 0.792 | cytoplasm | K12881 | A | KOG0533 RRM motif-containing protein |
| P45880 | VDAC2 | 31.566 | 142.4 | 60.9 | 14 | 22 | 14 | 5.72E+08 | 1.22E+09 | 0.788 | cytoplasm | K15040 | P | KOG3126 Porin/voltage-dependent anion-selective channel protein |
| Q7Z406 | MYH14 | 227.87 | 37.842 | 8.1 | 16 | 26 | 5 | 1.67E+08 | 2E+09 | 0.788 | nucleus | K10352 | Z | KOG0161 Myosin class II heavy chain |
| P05388 | RPLP0 | 34.273 | 26.453 | 20.8 | 4 | 4 | 4 | 68020000 | 1.36E+08 | 0.78 | cytoplasm,nucleus | K02941 | J | KOG0815 60S acidic ribosomal protein P0 |
| Q07021 | C1QBP | 31.362 | 23.571 | 21.6 | 3 | 3 | 3 | 32473000 | 1.04E+08 | 0.778 | mitochondria | K15414 | V | KOG4024 Complement component 1, Q subcomponent binding protein |
| P21796 | VDAC1 | 30.772 | 176.48 | 79.2 | 16 | 29 | 16 | 1.01E+09 | 2.48E+09 | 0.77 | cytoplasm | K05862 | P | KOG3126 Porin/voltage-dependent anion-selective channel protein |
| P39023 | RPL3 | 46.108 | 18.71 | 14.4 | 3 | 3 | 3 | 39271000 | 1.08E+08 | 0.77 | cytoplasm | K02925 | J | KOG0746 60S ribosomal protein L3 and related proteins |
| P60660 | MYL6 | 16.93 | 135.52 | 81.5 | 12 | 37 | 10 | 2.94E+09 | 1.82E+10 | 0.767 | cytoplasm | K12751 | Z | KOG0030 Myosin essential light chain, EF-Hand protein superfamily |
| P15880 | RPS2 | 31.324 | 78.112 | 42.7 | 10 | 11 | 10 | 2.5E+08 | 4.63E+08 | 0.76 | cytoplasm | K02981 | J | KOG0877 40S ribosomal protein S2/30S ribosomal protein S5 |
| P15559 | NQO1 | 30.867 | 38.08 | 22.6 | 4 | 6 | 4 | 1.7E+08 | 3.5E+08 | 0.759 | cytoplasm | K00355 | | |
| P36542 | ATP5F1C | 32.996 | 12.471 | 6 | 2 | 2 | 2 | 20420000 | 31931000 | 0.757 | mitochondria | K02136 | C | KOG1531 F0F1-type ATP synthase, gamma subunit |
| P62424 | RPL7A | 29.995 | 84.3 | 37.6 | 10 | 11 | 10 | 2.38E+08 | 3.96E+08 | 0.757 | cytoplasm | K02936 | J | KOG3166 60S ribosomal protein L7A |
| P05204 | HMGN2 | 9.3926 | 27.167 | 16.7 | 4 | 6 | 4 | 56696000 | 69429000 | 0.756 | nucleus | K11300 | | |
| Q8TDN6 | BRIX1 | 41.401 | 61.244 | 30.6 | 8 | 11 | 8 | 1.79E+08 | 4.4E+08 | 0.754 | nucleus | K14820 | J | KOG2971 RNA-binding protein required for biogenesis of the ribosomal 60S subunit |
| Q9Y2X3 | NOP58 | 59.578 | 122.37 | 36.1 | 13 | 18 | 13 | 3.39E+08 | 6.82E+08 | 0.753 | cytoplasm | K14565 | AJ | KOG2572 Ribosome biogenesis protein - Nop58p/Nop5p |
| Q9Y221 | NIP7 | 20.462 | 22.283 | 29.4 | 3 | 3 | 3 | 46442000 | 69574000 | 0.752 | cytoplasm | K07565 | J | KOG3492 Ribosome biogenesis protein NIP7 |
| P04406 | GAPDH | 36.053 | 139.2 | 68.7 | 15 | 28 | 15 | 1.03E+09 | 2.3E+09 | 0.74 | cytoplasm | K00134 | G | KOG0657 Glyceraldehyde 3-phosphate dehydrogenase |
| P61353 | RPL27 | 15.798 | 31.496 | 36 | 4 | 5 | 4 | 1.35E+08 | 2.7E+08 | 0.737 | mitochondria | K02901 | J | KOG3418 60S ribosomal protein L27 |

| | | | | | | | | | | | | | | |
|---|---|---|---|---|---|---|---|---|---|---|---|---|---|---|
| O00422 | SAP18 | 17.561 | 71.412 | 62.1 | 9 | 12 | 9 | 3.05E+08 | 5.32E+08 | 0.722 | nucleus | K14324 | K | KOG3391 Transcriptional co-repressor component |
| P56537 | EIF6 | 26.599 | 27.552 | 17.1 | 2 | 3 | 2 | 72516000 | 1.24E+08 | 0.717 | cytoplasm,nucleus | K03264 | J | KOG3185 Translation initiation factor 6 (eIF-6) |
| O76021 | RSL1D1 | 54.972 | 134.21 | 30 | 15 | 20 | 15 | 7.98E+08 | 1.65E+09 | 0.714 | nucleus | K14775 | S | KOG1685 Uncharacterized conserved protein |
| P23396 | RPS3 | 26.688 | 125.98 | 67.5 | 16 | 18 | 16 | 5.34E+08 | 9.96E+08 | 0.711 | cytoplasm | K02985 | J | KOG3181 40S ribosomal protein S3 |
| Q5JTH9 | RRP12 | 143.7 | 23.91 | 3.9 | 4 | 4 | 4 | 34147000 | 37577000 | 0.71 | nucleus | K14794 | S | KOG1248 Uncharacterized conserved protein |
| Q15397 | PUM3 | 73.584 | 40.877 | 13.1 | 5 | 5 | 5 | 1.07E+08 | 2.28E+08 | 0.709 | nucleus | K14844 | J | KOG2050 Puf family RNA-binding protein |
| Q9BYG3 | NIFK | 34.222 | 48.093 | 27.3 | 5 | 5 | 5 | 88678000 | 1.41E+08 | 0.708 | nucleus | K14838 | R | KOG4208 Nucleolar RNA-binding protein NIFK |
| P46779 | RPL28 | 15.747 | 17.872 | 18.2 | 3 | 3 | 3 | 22805000 | 45762000 | 0.701 | mitochondria | K02903 | J | KOG3412 60S ribosomal protein L28 |
| P06703 | S100A6 | 10.18 | 35.47 | 74.4 | 5 | 7 | 5 | 5.37E+08 | 1.11E+09 | 0.697 | cytoplasm | | | |
| P18621 | RPL17 | 21.397 | 25.202 | 23.9 | 3 | 5 | 3 | 95331000 | 1.43E+08 | 0.693 | cytoplasm | K02880 | J | KOG3353 60S ribosomal protein L22 |
| Q02878 | RPL6 | 32.728 | 81.771 | 37.2 | 11 | 16 | 11 | 2.96E+08 | 4.13E+08 | 0.693 | cytoplasm | K02934 | J | KOG1694 60s ribosomal protein L6 |
| Q12905 | ILF2 | 43.062 | 174.06 | 65.6 | 17 | 25 | 17 | 1.44E+09 | 3.32E+09 | 0.689 | cytoplasm | K13089 | K | KOG3793 Transcription factor NFAT, subunit NF45 |
| P62280 | RPS11 | 18.431 | 49.374 | 39.2 | 6 | 8 | 6 | 1.97E+08 | 3.53E+08 | 0.684 | cytoplasm | K02949 | J | KOG1728 40S ribosomal protein S11 |
| P62888 | RPL30 | 12.784 | 42.476 | 59.1 | 5 | 6 | 5 | 1.13E+08 | 1.62E+08 | 0.678 | cytoplasm | K02908 | J | KOG2988 60S ribosomal protein L30 |
| Q9UNX3 | RPL26L1 | 17.256 | 25.098 | 22.8 | 4 | 4 | 4 | 90882000 | 1.21E+08 | 0.674 | nucleus | K02898 | J | KOG3401 60S ribosomal protein L26 |
| P55209 | NAP1L1 | 45.374 | 21.934 | 11.8 | 3 | 3 | 3 | 61753000 | 64006000 | 0.67 | cytoplasm,nucleus | K11279 | BD | KOG1507 Nucleosome assembly protein NAP-1 |
| P62906 | RPL10A | 24.831 | 53.658 | 31.3 | 6 | 10 | 6 | 2.77E+08 | 4.28E+08 | 0.665 | cytoplasm,nucleus | K02865 | J | KOG1570 60S ribosomal protein L10A |
| Q9UKM9 | RALY | 32.463 | 93.93 | 35.9 | 10 | 13 | 10 | 8.7E+08 | 1.77E+09 | 0.663 | nucleus | K12895 | R | KOG0118 FOG: RRM domain |
| P12956 | XRCC6 | 69.842 | 12.043 | 3.3 | 2 | 2 | 2 | 17643000 | 19575000 | 0.661 | cytoplasm | K10884 | L | KOG2327 DNA-binding subunit of a DNA-dependent protein kinase (Ku70 autoantigen) |
| P00403 | MT-CO2 | 25.565 | 38.93 | 36.1 | 6 | 6 | 6 | 1.62E+08 | 4.85E+08 | 0.657 | plasma membrane | K02261 | C | KOG4767 Cytochrome c oxidase, subunit II, and related proteins |
| P62269 | RPS18 | 17.718 | 60.012 | 39.5 | 8 | 9 | 8 | 3.41E+08 | 6.87E+08 | 0.655 | cytoplasm | K02964 | J | KOG3311 Ribosomal protein S18 |
| P09661 | SNRPA1 | 28.415 | 59.861 | 25.5 | 5 | 9 | 5 | 91111000 | 2.69E+08 | 0.651 | cytoplasm,nucleus | K11092 | A | KOG1644 U2-associated snRNP A' protein |
| P63167 | DYNLL1 | 10.366 | 21.284 | 37.1 | 2 | 4 | 1 | 61457000 | 1.12E+08 | 0.651 | cytoplasm | K10418 | Z | KOG3430 Dynein light chain type 1 |
| Q9NR12 | PDLIM7 | 49.844 | 27.028 | 17.9 | 4 | 4 | 4 | 86490000 | 1.08E+08 | 0.649 | nucleus | | TZ | KOG1703 Adaptor protein Enigma and related PDZ-LIM proteins |
| Q07020 | RPL18 | 21.634 | 36.453 | 25 | 4 | 4 | 4 | 1.55E+08 | 3.12E+08 | 0.646 | cytoplasm | K02883 | J | KOG1714 60s ribosomal protein L18 |
| Q9Y265 | RUVBL1 | 50.227 | 20.391 | 11.6 | 3 | 4 | 3 | 44830000 | 55353000 | 0.628 | cytoplasm | K04499 | L | KOG1942 DNA helicase, TBP-interacting protein |
| P18124 | RPL7 | 29.225 | 42.776 | 28.6 | 6 | 7 | 6 | 2.75E+08 | 3.73E+08 | 0.625 | cytoplasm | K02937 | J | KOG3184 60S ribosomal protein L7 |
| P19105 | MYL12A | 19.794 | 179.35 | 84.2 | 15 | 50 | 1 | 3.05E+09 | 2.09E+10 | 0.625 | mitochondria | K12757 | Z | KOG0031 Myosin regulatory light chain, EF-Hand protein superfamily |
| Q9UKD2 | MRTO4 | 27.56 | 12.488 | 9.6 | 2 | 3 | 2 | 57565000 | 74974000 | 0.623 | nucleus | K14815 | A | KOG0816 Protein involved in mRNA turnover |
| Q71UM5 | RPS27L | 9.4771 | 6.5039 | 15.5 | 1 | 2 | 1 | 60770000 | 88218000 | 0.622 | extracellular | K02978 | J | KOG1779 40s ribosomal protein S27 |
| P63241 | EIF5A | 16.832 | 57.264 | 51.3 | 6 | 8 | 6 | 1.28E+08 | 1.85E+08 | 0.621 | cytoplasm | K03263 | J | KOG3271 Translation initiation factor 5A (eIF-5A) |
| O96008 | TOMM40 | 37.893 | 85.027 | 53.7 | 9 | 13 | 9 | 4.5E+08 | 7.87E+08 | 0.62 | nucleus | K11518 | U | KOG3296 Translocase of outer mitochondrial membrane complex, subunit TOM40 |
| P06748 | NPM1 | 32.575 | 144.37 | 46.9 | 12 | 29 | 12 | 3.07E+09 | 5.91E+09 | 0.618 | nucleus | K11276 | | |
| P31946 | YWHAB | 28.082 | 22.27 | 22.8 | 4 | 4 | 2 | 57969000 | 57058000 | 0.618 | cytoplasm | K16197 | O | KOG0841 Multifunctional chaperone (14-3-3 family) |
| P08708 | RPS17 | 15.55 | 24.623 | 40 | 3 | 6 | 3 | 2.92E+08 | 4.32E+08 | 0.615 | cytoplasm | K02962 | J | KOG0187 40S ribosomal protein S17 |
| P49207 | RPL34 | 13.293 | 12.89 | 12.8 | 2 | 2 | 2 | 44205000 | 58632000 | 0.614 | nucleus | K02915 | J | KOG1790 60s ribosomal protein L34 |
| P62841 | RPS15 | 17.04 | 38.447 | 55.2 | 4 | 8 | 4 | 3.17E+08 | 6.08E+08 | 0.612 | cytoplasm | K02958 | J | KOG0898 40S ribosomal protein S15 |
| P26447 | S100A4 | 11.728 | 12.93 | 18.8 | 2 | 2 | 2 | 93639000 | 1.8E+08 | 0.61 | extracellular | | | |
| P62277 | RPS13 | 17.222 | 51.239 | 39.1 | 7 | 9 | 7 | 5.5E+08 | 1.2E+09 | 0.604 | cytoplasm | K02953 | J | KOG0400 40S ribosomal protein S13 |
| Q9H7B2 | RPF2 | 35.582 | 42.586 | 21.9 | 6 | 8 | 6 | 1.4E+08 | 2.08E+08 | 0.603 | nucleus | K14847 | J | KOG3031 Protein required for biogenesis of the ribosomal 60S subunit |
| P62854 | RPS26 | 13.015 | 15.56 | 23.5 | 2 | 3 | 2 | 48127000 | 1E+08 | 0.602 | cytoplasm | | J | KOG1768 40s ribosomal protein S26 |
| Q13765 | NACA | 23.384 | 20.383 | 13 | 2 | 2 | 2 | 24197000 | 28341000 | 0.602 | cytoplasm | K03626 | K | KOG2239 Transcription factor containing NAC and TS-N domains |
| P62829 | RPL23 | 14.865 | 44.385 | 40 | 5 | 6 | 5 | 2.54E+08 | 3.93E+08 | 0.599 | cytoplasm | K02894 | J | KOG0901 60S ribosomal protein L14/L17/L23 |
| P42766 | RPL35 | 14.551 | 25.603 | 26 | 3 | 4 | 3 | 1.54E+08 | 1.98E+08 | 0.598 | cytoplasm | K02918 | J | KOG3436 60S ribosomal protein L35 |
| P46781 | RPS9 | 22.591 | 44.901 | 26.8 | 7 | 8 | 7 | 1.33E+08 | 2.24E+08 | 0.593 | cytoplasm | K02997 | J | KOG3301 Ribosomal protein S4 |
| P39019 | RPS19 | 16.06 | 55.907 | 38.6 | 8 | 9 | 8 | 2.98E+08 | 5.26E+08 | 0.588 | cytoplasm | K02966 | J | KOG3411 40S ribosomal protein S19 |
| P83731 | RPL24 | 17.779 | 13.67 | 13.4 | 2 | 2 | 2 | 61961000 | 82579000 | 0.587 | cytoplasm | K02896 | J | KOG1722 60s ribosomal protein L24 |
| Q15366 | PCBP2 | 38.58 | 73.114 | 40.5 | 8 | 12 | 7 | 4.08E+08 | 6.04E+08 | 0.577 | cytoskeleton | K13162 | AR | KOG2190 PolyC-binding proteins alphaCP-1 and related KH domain proteins |
| P19338 | NCL | 76.613 | 125.15 | 22.4 | 12 | 18 | 12 | 7.46E+08 | 1.24E+09 | 0.574 | nucleus | K11294 | | |
| Q14974 | KPNB1 | 97.169 | 12.635 | 2.7 | 2 | 2 | 2 | 24081000 | 32380000 | 0.574 | cytoplasm | K14293 | YU | KOG1241 Karyopherin (importin) beta 1 |
| Q9H6F5 | CCDC86 | 40.235 | 22.937 | 14.7 | 4 | 4 | 4 | 42139000 | 52210000 | 0.574 | nucleus | K14822 | R | KOG4538 Predicted coiled-coil protein |

| | | | | | | | | | | | | | | | |
|---|---|---|---|---|---|---|---|---|---|---|---|---|---|---|---|
| P05387 | RPLP2 | 11.665 | 34.534 | 55.7 | 3 | 5 | 3 | 2.14E+08 | 3.2E+08 | 0.571 | mitochondria | K02943 | J | KOG3449 | 60S acidic ribosomal protein P2 |
| P84243 | H3F3A | 15.328 | 26.518 | 61.8 | 12 | 32 | 2 | 2.33E+09 | 5.39E+09 | 0.567 | nucleus | K11253 | B | KOG1745 | Histones H3 and H4 |
| O75475 | PSIP1 | 60.103 | 84.796 | 19.8 | 8 | 14 | 8 | 5.52E+08 | 7.73E+08 | 0.562 | nucleus | | K | KOG1904 | Transcription coactivator |
| P62750 | RPL23A | 17.695 | 52.689 | 43.6 | 7 | 8 | 7 | 5.71E+08 | 8.69E+08 | 0.56 | cytoplasm | K02893 | J | KOG1751 | 60s ribosomal protein L23 |
| O75531 | BANF1 | 10.058 | 59.517 | 70.8 | 7 | 14 | 7 | 2.79E+09 | 5.27E+09 | 0.556 | extracellular | K21870 | BL | KOG4233 | DNA-bridging protein BAF |
| P62861 | FAU | 6.6478 | 13.507 | 18.6 | 2 | 2 | 2 | 83690000 | 1.29E+08 | 0.556 | nucleus | K02983 | JO | KOG0009 | Ubiquitin-like/40S ribosomal S30 protein fusion |
| Q9Y3U8 | RPL36 | 12.254 | 21.984 | 21 | 3 | 3 | 3 | 1.07E+08 | 1.59E+08 | 0.556 | nucleus | K02920 | J | KOG3452 | 60S ribosomal protein L36 |
| P05386 | RPLP1 | 11.514 | 13.495 | 51.8 | 2 | 2 | 2 | 1.49E+08 | 2.32E+08 | 0.549 | mitochondria | K02942 | J | KOG1762 | 60s acidic ribosomal protein P1 |
| O14880 | MGST3 | 16.516 | 60.521 | 53.3 | 6 | 8 | 6 | 2.27E+08 | 1.09E+09 | 0.533 | extracellular | K00799 | | | |
| P17096 | HMGA1 | 11.676 | 45.363 | 23.4 | 4 | 7 | 4 | 3.18E+08 | 6.19E+08 | 0.533 | nucleus | K09282 | | | |
| P46776 | RPL27A | 16.561 | 26.317 | 22.3 | 4 | 4 | 4 | 1.45E+08 | 1.4E+08 | 0.528 | cytoplasm | K02900 | J | KOG1742 | 60s ribosomal protein L15/L27 |
| P68104 | EEF1A1 | 50.14 | 166.43 | 52.2 | 17 | 29 | 8 | 1.61E+09 | 2.65E+09 | 0.526 | cytoplasm | K03231 | J | KOG0052 | Translation elongation factor EF-1 alpha/Tu |
| P62899 | RPL31 | 14.463 | 21.781 | 24.8 | 3 | 4 | 3 | 3.06E+08 | 4.24E+08 | 0.518 | cytoplasm | K02910 | J | KOG0893 | 60S ribosomal protein L31 |
| P61247 | RPS3A | 29.945 | 61.179 | 34.5 | 8 | 10 | 8 | 2.94E+08 | 3.69E+08 | 0.511 | nucleus | K02984 | J | KOG1628 | 40S ribosomal protein S3A |
| P46778 | RPL21 | 18.565 | 15.276 | 16.2 | 2 | 2 | 2 | 66782000 | 69890000 | 0.504 | nucleus | K02889 | J | KOG1732 | 60S ribosomal protein L21 |
| P62081 | RPS7 | 22.127 | 60.573 | 52.1 | 8 | 11 | 8 | 6.25E+08 | 9.42E+08 | 0.499 | cytoplasm | K02993 | J | KOG3320 | 40S ribosomal protein S7 |
| Q92979 | EMG1 | 26.72 | 18.191 | 18 | 3 | 3 | 3 | 62994000 | 62197000 | 0.496 | nucleus | K14568 | | | |
| P45973 | CBX5 | 22.225 | 21.197 | 30.9 | 4 | 6 | 3 | 1.21E+08 | 99991000 | 0.48 | nucleus | K11587 | | | |
| P62266 | RPS23 | 15.807 | 21.79 | 27.3 | 3 | 3 | 3 | 1.02E+08 | 1.63E+08 | 0.478 | cytoplasm | K02973 | J | KOG1749 | 40S ribosomal protein S23 |
| P62249 | RPS16 | 16.445 | 31.819 | 26.7 | 4 | 5 | 4 | 2.13E+08 | 3.16E+08 | 0.477 | cytoplasm | K02960 | J | KOG1753 | 40S ribosomal protein S16 |
| P60866 | RPS20 | 13.373 | 24.91 | 22.7 | 3 | 6 | 3 | 4.49E+08 | 6.5E+08 | 0.471 | cytoplasm | K02969 | J | KOG0900 | 40S ribosomal protein S20 |
| P62917 | RPL8 | 28.024 | 12.285 | 10.5 | 2 | 2 | 2 | 17638000 | 20150000 | 0.464 | cytoplasm | K02938 | J | KOG2309 | 60s ribosomal protein L2/L8 |
| P35268 | RPL22 | 14.787 | 23.328 | 30.5 | 2 | 2 | 2 | 2.33E+08 | 4.18E+08 | 0.459 | nucleus | K02891 | J | KOG3434 | 60S ribosomal protein L22 |
| P62847 | RPS24 | 15.423 | 18.4 | 20.3 | 2 | 2 | 2 | 1.89E+08 | 3.15E+08 | 0.456 | cytoplasm | K02974 | J | KOG3424 | 40S ribosomal protein S24 |
| P26373 | RPL13 | 24.261 | 22.164 | 11.4 | 2 | 3 | 2 | 1.42E+08 | 4.25E+08 | 0.446 | cytoplasm | K02873 | J | KOG3295 | 60S Ribosomal protein L13 |
| P07910 | HNRNPC | 33.67 | 153.89 | 50.7 | 17 | 34 | 17 | 3.5E+09 | 7.76E+09 | 0.441 | nucleus | K12884 | R | KOG0118 | FOG: RRM domain |
| P62979 | RPS27A | 17.965 | 55.233 | 42.3 | 6 | 14 | 6 | 7.7E+08 | 1.55E+09 | 0.44 | extracellular | K02977 | J | KOG0004 | Ubiquitin/40S ribosomal protein S27a fusion |
| P62851 | RPS25 | 13.742 | 21.177 | 22.4 | 3 | 3 | 3 | 90047000 | 90490000 | 0.434 | nucleus | K02975 | J | KOG1767 | 40S ribosomal protein S25 |
| P57053 | H2BFS | 13.944 | 110.97 | 60.3 | 12 | 62 | 6 | 1.35E+10 | 1.44E+10 | 0.421 | nucleus | | B | KOG1744 | Histone H2B |
| P26599 | PTBP1 | 57.221 | 220.5 | 64.2 | 21 | 33 | 18 | 2.72E+09 | 4.72E+09 | 0.411 | cytoplasm,nucleus | K13218 | A | KOG1190 | Polypyrimidine tract-binding protein |
| P62805 | HIST1H4A | 11.367 | 271.06 | 86.4 | 27 | 70 | 27 | 2.16E+10 | 2.32E+10 | 0.344 | nucleus | K11254 | B | KOG3467 | Histone H4 |
| P35659 | DEK | 42.674 | 16.472 | 6.7 | 2 | 2 | 2 | 70443000 | 47831000 | 0.331 | nucleus | K17046 | B | KOG2266 | Chromatin-associated protein Dek and related proteins |
| Q13185 | CBX3 | 20.811 | 43.747 | 45.9 | 5 | 6 | 3 | 4.77E+08 | 6.11E+08 | 0.331 | nucleus | K11586 | B | KOG1911 | Heterochromatin-associated protein HP1 and related CHROMO domain proteins |
| P09429 | HMGB1 | 24.893 | 53.2 | 39.1 | 7 | 8 | 5 | 3.32E+08 | 1.89E+08 | 0.297 | nucleus | K10802 | R | KOG0381 | HMG box-containing protein |
| Q71DI3 | HIST2H3A | 15.388 | 114.43 | 72.1 | 14 | 39 | 3 | 2.17E+10 | 2.3E+10 | 0.276 | nucleus | K11253 | B | KOG1745 | Histones H3 and H4 |
| P62244 | RPS15A | 14.839 | 23.841 | 29.2 | 4 | 4 | 4 | 1.77E+08 | 2.36E+08 | 0.264 | cytoplasm | K02957 | J | KOG1754 | 40S ribosomal protein S15/S22 |
| O15347 | HMGB3 | 22.98 | 14.08 | 14.5 | 2 | 3 | 2 | 1.31E+08 | 56663000 | 0.242 | nucleus | K11296 | R | KOG0381 | HMG box-containing protein |
| P68431 | HIST1H3A | 15.404 | 34.78 | 72.1 | 14 | 36 | 3 | 6.68E+09 | 6.98E+09 | 0.225 | nucleus | K11253 | B | KOG1745 | Histones H3 and H4 |
| Q71UI9 | H2AFV | 13.509 | 50.682 | 53.9 | 6 | 16 | 4 | 2.26E+09 | 6.77E+09 | 0.201 | nucleus | K11251 | B | KOG1757 | Histone 2A |
| Q16777 | HIST2H2AC | 13.988 | 87.492 | 58.1 | 8 | 37 | 1 | 5.48E+09 | 6.18E+09 | 0.17 | nucleus | K11251 | B | KOG1756 | Histone 2A |
| P16401 | HIST1H1B | 22.58 | 45.529 | 25.2 | 7 | 19 | 6 | 4.31E+08 | 2.27E+08 | 0.168 | nucleus | K11275 | B | KOG4012 | Histone H1 |
| P68366 | TUBA4A | 49.924 | 20.498 | 54 | 17 | 28 | 3 | 3.6E+08 | 1.84E+08 | 0.165 | cytoskeleton | K07374 | Z | KOG1376 | Alpha tubulin |
| Q16778 | HIST2H2BE | 13.92 | 25.117 | 54.8 | 9 | 55 | 1 | 2.35E+09 | 2.8E+09 | 0.15 | nucleus | K11252 | B | KOG1744 | Histone H2B |
| P16403 | HIST1H1C | 21.364 | 70.975 | 28.6 | 9 | 22 | 8 | 2.44E+09 | 1.11E+09 | 0.107 | nucleus | K11275 | B | KOG4012 | Histone H1 |
| A6NHQ2 | FBLL1 | 34.803 | 13.4 | 6.9 | 3 | 5 | 2 | 1.28E+08 | 4.59E+08 | | nucleus | K14563 | | | |
| A6NMZ7 | COL6A6 | 247.17 | 5.6244 | 0.3 | 1 | 2 | 1 | 8863100 | 32431000 | | endoplasmic reticulur | K06238 | W | KOG3544 | Collagens (type IV and type XIII), and related proteins |
| O00299 | CLIC1 | 26.922 | 6.2216 | 9.1 | 1 | 1 | 1 | 0 | 14455000 | | cytoplasm | K05021 | | | |
| O00425 | IGF2BP3 | 63.704 | 5.7634 | 4.7 | 2 | 2 | 1 | 7740500 | 25114000 | | cytoplasm | K13197 | AR | KOG2193 | IGF-II mRNA-binding protein IMP, contains RRM and KH domains |
| O00764 | PDXK | 35.102 | 7.3517 | 6.7 | 1 | 1 | 1 | 0 | 15943000 | | cytoplasm | K00868 | H | KOG2599 | Pyridoxal/pyridoxine/pyridoxamine kinase |
| O14548 | COX7A2L | 12.615 | 7.4336 | 11.4 | 1 | 1 | 1 | 26040000 | 8935100 | | cytoplasm | K02270 | | | |
| O14561 | NDUFAB1 | 17.417 | 6.665 | 9.6 | 1 | 1 | 1 | 19076000 | 42846000 | | mitochondria | K03955 | CIQ | KOG1748 | Acyl carrier protein/NADH-ubiquinone oxidoreductase |

| | | | | | | | | | | | | | | |
|---|---|---|---|---|---|---|---|---|---|---|---|---|---|---|
| O14949 | UQCRQ | 9.9062 | 19.184 | 24.4 | 2 | 2 | 2 | 16497000 | 33371000 | mitochondria | K00418 | C | KOG4116 | Ubiquinol cytochrome c reductase, subunit QCR8 |
| O14974 | PPP1R12A | 115.28 | 18.455 | 3.5 | 2 | 3 | 2 | 1496200 | 40535000 | nucleus | K06270 | OT | KOG0505 | Myosin phosphatase, regulatory subunit |
| O14979 | HNRNPDL | 46.437 | 9.3153 | 5.7 | 2 | 2 | 1 | 7332800 | 26663000 | nucleus | K13044 | | | |
| O15143 | ARPC1B | 40.949 | 6.4187 | 2.7 | 1 | 1 | 1 | 0 | 6607200 | cytoplasm | K05757 | Z | KOG1523 | Actin-related protein Arp2/3 complex, subunit ARPC1/p41-ARC |
| O15144 | ARPC2 | 34.333 | 5.7582 | 3.7 | 1 | 1 | 1 | 0 | 8704700 | cytoplasm | K05758 | Z | KOG2826 | Actin-related protein Arp2/3 complex, subunit ARPC2 |
| O15514 | POLR2D | 16.311 | 8.7078 | 14.1 | 1 | 1 | 1 | 8240600 | 41519000 | cytoplasm | K03012 | K | KOG2351 | RNA polymerase II, fourth largest subunit |
| O15523 | DDX3Y | 73.153 | 8.749 | 2.6 | 1 | 1 | 1 | 0 | 8379300 | nucleus | K17642 | A | KOG0335 | ATP-dependent RNA helicase |
| O43143 | DHX15 | 90.932 | 5.6074 | 1.5 | 1 | 1 | 1 | 0 | 15867000 | nucleus | K12820 | A | KOG0925 | mRNA splicing factor ATP-dependent RNA helicase |
| O43159 | RRP8 | 50.714 | 9.3315 | 3.5 | 1 | 1 | 1 | 0 | 0 | nucleus | K14850 | A | KOG3045 | Predicted RNA methylase involved in rRNA processing |
| O43504 | LAMTOR5 | 9.6138 | 14.13 | 15.4 | 1 | 1 | 1 | 0 | 15379000 | cytoskeleton | K16344 | | | |
| O43570 | CA12 | 39.45 | 15.077 | 9.9 | 2 | 2 | 2 | 5477700 | 56606000 | extracellular | K01672 | R | KOG0382 | Carbonic anhydrase |
| O43678 | NDUFA2 | 10.921 | 5.7938 | 21.2 | 1 | 1 | 1 | 5403700 | 13281000 | mitochondria | K03946 | C | KOG3446 | NADH:ubiquinone oxidoreductase NDUFA2/B8 subunit |
| O43852 | CALU | 37.106 | 6.8954 | 6 | 1 | 1 | 1 | 3746800 | 14911000 | extracellular | | TU | KOG4223 | Reticulocalbin, calumenin, DNA supercoiling factor of the CREC family (EF-Hand protein superfamily) |
| O60664 | PLIN3 | 47.074 | 10.538 | 3.7 | 1 | 1 | 1 | 17069000 | 26341000 | cytoplasm | K20287 | | | |
| O60784 | TOM1 | 53.818 | 6.8881 | 3.5 | 1 | 1 | 1 | 0 | 24041000 | cytoplasm | | U | KOG1087 | Cytosolic sorting protein GGA2/TOM1 |
| O60828 | PQBP1 | 30.472 | 12.519 | 13.2 | 2 | 2 | 2 | 0 | 28920000 | nucleus | K12865 | K | KOG3427 | Polyglutamine tract-binding protein PQBP-1 |
| O75251 | NDUFS7 | 23.563 | 11.116 | 6.6 | 1 | 1 | 1 | 6197300 | 23206000 | mitochondria | K03940 | C | KOG1687 | NADH-ubiquinone oxidoreductase, NUFS7/PSST/20 kDa subunit |
| O75306 | NDUFS2 | 52.545 | 5.6926 | 1.5 | 1 | 1 | 1 | 0 | 4280500 | mitochondria | K03935 | C | KOG2870 | NADH:ubiquinone oxidoreductase, NDUFS2/49 kDa subunit |
| O75396 | SEC22B | 24.593 | 7.6914 | 6.5 | 1 | 1 | 1 | 3243400 | 24880000 | endoplasmic reticulu | K08517 | | | |
| O75431 | MTX2 | 29.763 | 11.571 | 8.4 | 1 | 2 | 1 | 52956000 | 1.37E+08 | cytoplasm | K17776 | MU | KOG3027 | Mitochondrial outer membrane protein Metaxin 2, Metaxin 1-binding protein |
| O75533 | SF3B1 | 145.83 | 17.897 | 2.4 | 2 | 2 | 2 | 7789400 | 37550000 | plasma membrane | K12828 | A | KOG0213 | Splicing factor 3b, subunit 1 |
| O75683 | SURF6 | 41.45 | 5.4869 | 1.9 | 1 | 1 | 1 | 0 | 0 | nucleus | | S | KOG2885 | Uncharacterized conserved protein |
| O75874 | IDH1 | 46.659 | 12.158 | 4.8 | 2 | 2 | 2 | 4102100 | 36785000 | cytoplasm | K00031 | C | KOG1526 | NADP-dependent isocitrate dehydrogenase |
| O75880 | SCO1 | 33.814 | 6.1446 | 5.3 | 1 | 1 | 1 | 9553500 | 40285000 | mitochondria | K07152 | C | KOG2792 | Putative cytochrome C oxidase assembly protein |
| O75940 | SMNDC1 | 26.711 | 15.222 | 13 | 2 | 3 | 2 | 6937200 | 22542000 | nucleus | K12839 | A | KOG3026 | Splicing factor SPF30 |
| O94826 | TOMM70 | 67.454 | 12.805 | 5.8 | 2 | 2 | 2 | 10158000 | 64091000 | endoplasmic reticulu | K17768 | U | KOG0547 | Translocase of outer mitochondrial membrane complex, subunit TOM70/TOM72 |
| O94889 | KLHL18 | 63.638 | 5.5906 | 1.2 | 1 | 1 | 1 | 20703000 | 2E+08 | nucleus | K10455 | TR | KOG4441 | Proteins containing BTB/POZ and Kelch domains |
| O94973 | AP2A2 | 103.96 | 11.892 | 2.6 | 2 | 2 | 1 | 2567400 | 12611000 | cytoplasm | K11824 | U | KOG1077 | Vesicle coat complex AP-2, alpha subunit |
| O95182 | NDUFA7 | 12.551 | 6.7433 | 15.9 | 1 | 1 | 1 | 13278000 | 41743000 | mitochondria | K03951 | C | KOG4630 | NADH:ubiquinone oxidoreductase, NDUFA7/B14.5A subunit |
| O95425 | SVIL | 247.74 | 34.27 | 3.4 | 4 | 4 | 4 | 3083000 | 66999000 | nucleus | K10369 | Z | KOG0445 | Actin regulatory protein supervillin (gelsolin/villin family) |
| O95613 | PCNT | 378.03 | -2 | 0.4 | 1 | 1 | 1 | 0 | 0 | nucleus | K16481 | | | |
| O95782 | AP2A1 | 107.54 | 6.3704 | 3.9 | 2 | 2 | 1 | 3958200 | 38005000 | cytoplasm | K11824 | U | KOG1077 | Vesicle coat complex AP-2, alpha subunit |
| O95793 | STAU1 | 63.182 | 11.882 | 4.5 | 2 | 2 | 2 | 12110000 | 23381000 | nucleus | K17597 | UK | KOG3732 | Staufen and related double-stranded-RNA-binding proteins |
| O95816 | BAG2 | 23.772 | 5.4999 | 10.4 | 1 | 1 | 1 | 3141000 | 13503000 | cytoplasm | K09556 | O | KOG3633 | BAG family molecular chaperone regulator 2 |
| P00387 | CYB5R3 | 34.234 | 17.366 | 14.3 | 3 | 3 | 3 | 0 | 49408000 | extracellular | K00326 | HC | KOG0534 | NADH-cytochrome b-5 reductase |
| P00450 | CP | 122.2 | 5.9239 | 1.6 | 1 | 1 | 1 | 0 | 17352000 | endoplasmic reticulu | K13624 | Q | KOG1263 | Multicopper oxidases |
| P01116 | KRAS | 21.656 | 7.7576 | 13.2 | 1 | 1 | 1 | 15914000 | 39660000 | cytoplasm | K07827 | R | KOG0395 | Ras-related GTPase |
| P02792 | FTL | 20.019 | 9.622 | 9.1 | 1 | 1 | 1 | 16635000 | 15554000 | cytoplasm | K13625 | | | |
| P02794 | FTH1 | 21.225 | 6.2798 | 8.2 | 1 | 1 | 1 | 7740200 | 10069000 | cytoplasm | K00522 | P | KOG2332 | Ferritin |
| P04080 | CSTB | 11.139 | 9.9269 | 21.4 | 1 | 2 | 1 | 0 | 14073000 | cytoplasm | K13907 | | | |
| P04156 | PRNP | 27.661 | 9.9192 | 4.7 | 1 | 1 | 1 | 11905000 | 38649000 | extracellular | K05634 | | | |
| P04259 | KRT6B | 60.066 | 12.221 | 13.1 | 10 | 0 | 0 | 0 | 77065000 | nucleus | K07605 | | | |
| P04632 | CAPNS1 | 28.315 | 32.99 | 19.4 | 3 | 3 | 3 | 11330000 | 70495000 | cytoplasm | K08583 | T | KOG0037 | Ca2+-binding protein, EF-Hand protein superfamily |
| P05976 | MYL1 | 21.145 | 20.768 | 8.2 | 2 | 7 | 2 | 0 | 0 | cytoplasm | K05738 | Z | KOG0030 | Myosin essential light chain, EF-Hand protein superfamily |
| P06454 | PTMA | 12.203 | 6.9758 | 12.6 | 1 | 1 | 1 | 9756100 | 8035600 | cytoplasm | K13784 | | | |
| P07948 | LYN | 58.573 | 5.4748 | 4.3 | 2 | 2 | 1 | 0 | 0 | cytoplasm,nucleus | K05854 | T | KOG0197 | Tyrosine kinases |
| P08865 | RPSA | 32.854 | 26.808 | 23.1 | 3 | 3 | 3 | 8428400 | 42520000 | cytoplasm | K02998 | J | KOG0830 | 40S ribosomal protein SA (P40)/Laminin receptor 1 |
| P09622 | DLD | 54.177 | 7.496 | 3.7 | 1 | 1 | 1 | 6189300 | 44160000 | mitochondria | K00382 | C | KOG1335 | Dihydrolipoamide dehydrogenase |
| P0DPI3 | CENPVL2 | 29.887 | 6.8835 | 2.9 | 1 | 1 | 1 | 9927200 | 40970000 | mitochondria | | | | |
| P10515 | DLAT | 68.996 | 5.861 | 2.6 | 1 | 1 | 1 | 0 | 0 | mitochondria | K00627 | C | KOG0557 | Dihydrolipoamide acetyltransferase |
| P11217 | PYGM | 97.091 | 11.054 | 3.4 | 2 | 2 | 2 | 3901300 | 32647000 | cytoplasm | K00688 | G | KOG2099 | Glycogen phosphorylase |

| | | | | | | | | | | | | | | |
|---|---|---|---|---|---|---|---|---|---|---|---|---|---|---|
| P12235 | SLC25A4 | 33.064 | 6.7433 | 31.9 | 8 | 10 | 1 | 2799000 | 15242000 | mitochondria | K05863 | C | KOG0749 | Mitochondrial ADP/ATP carrier proteins |
| P12931 | SRC | 59.834 | 13.226 | 5.2 | 2 | 2 | 1 | 6490200 | 57499000 | cytoplasm | K05704 | T | KOG0197 | Tyrosine kinases |
| P13010 | XRCC5 | 82.704 | 9.5646 | 2.2 | 1 | 1 | 1 | 18904000 | 23683000 | cytoplasm | K10885 | L | KOG2326 | DNA-binding subunit of a DNA-dependent protein kinase (Ku80 autoantigen) |
| P13473 | LAMP2 | 44.96 | 17.701 | 4.9 | 2 | 2 | 2 | 5613100 | 53356000 | extracellular | K06528 | | | |
| P14174 | MIF | 12.476 | 6.4327 | 7.8 | 1 | 1 | 1 | 9031500 | 1.17E+08 | cytoplasm,nucleus | K07253 | V | KOG1759 | Macrophage migration inhibitory factor |
| P14550 | AKR1A1 | 36.573 | 7.7634 | 3.1 | 1 | 1 | 1 | 12381000 | 88726000 | cytoplasm | K00002 | R | KOG1577 | Aldo/keto reductase family proteins |
| P15531 | NME1 | 17.149 | 7.425 | 31.6 | 3 | 3 | 1 | 18359000 | 39178000 | cytoplasm | K00940 | F | KOG0888 | Nucleoside diphosphate kinase |
| P15954 | COX7C | 7.2454 | 6.3913 | 27 | 1 | 1 | 1 | 23051000 | 63299000 | mitochondria | K02272 | C | KOG4527 | Cytochrome c oxidase, subunit VIIc/COX8 |
| P16104 | H2AFX | 15.144 | 7.5736 | 52.4 | 6 | 19 | 0 | 0 | 0 | nucleus | K11251 | B | KOG1756 | Histone 2A |
| P16989 | YBX3 | 40.089 | 13.295 | 9.7 | 2 | 2 | 1 | 13092000 | 17262000 | nucleus | K06099 | J | KOG3070 | Predicted RNA-binding protein containing PIN domain |
| P17066 | HSPA6 | 71.027 | 9.0598 | 11.7 | 6 | 9 | 1 | 47196000 | 1.16E+08 | cytoplasm | K03283 | O | KOG0101 | Molecular chaperones HSP70/HSC70, HSP70 superfamily |
| P17480 | UBTF | 89.405 | 14.708 | 4.1 | 2 | 3 | 2 | 19316000 | 28691000 | nucleus | K09273 | R | KOG0381 | HMG box-containing protein |
| P17844 | DDX5 | 69.147 | 7.0362 | 2 | 1 | 1 | 1 | 3239200 | 8602000 | nucleus | K12823 | A | KOG0331 | ATP-dependent RNA helicase |
| P17931 | LGALS3 | 26.152 | 6.1018 | 6 | 1 | 1 | 1 | 0 | 0 | cytoplasm,nucleus | K06831 | W | KOG3587 | Galectin, galactose-binding lectin |
| P18085 | ARF4 | 20.511 | 33.278 | 16.7 | 2 | 2 | 1 | 14266000 | 67340000 | cytoplasm | K07939 | U | KOG0070 | GTP-binding ADP-ribosylation factor Arf1 |
| P19404 | NDUFV2 | 27.391 | 6.2706 | 5.2 | 1 | 1 | 1 | 7539100 | 27798000 | mitochondria | K03943 | C | KOG3196 | NADH:ubiquinone oxidoreductase, NDUFV2/24 kD subunit |
| P19525 | EIF2AK2 | 62.094 | 6.3859 | 2.2 | 1 | 1 | 1 | 0 | 19915000 | cytoplasm | K16195 | J | KOG1033 | eIF-2alpha kinase PEK/EIF2AK3 |
| P21291 | CSRP1 | 20.567 | 16.763 | 16.6 | 2 | 2 | 2 | 0 | 20787000 | nucleus | K09377 | | | |
| P23497 | SP100 | 100.42 | 6.9185 | 2.2 | 2 | 2 | 1 | 0 | 5457100 | nucleus | K15413 | O | KOG2177 | Predicted E3 ubiquitin ligase |
| P24539 | ATP5F1 | 28.908 | 7.529 | 4.7 | 1 | 1 | 1 | 2932800 | 20836000 | mitochondria | K02127 | | | |
| P25787 | PSMA2 | 25.898 | 7.4953 | 6 | 1 | 2 | 1 | 6262800 | 12291000 | cytoplasm | K02726 | O | KOG0181 | 20S proteasome, regulatory subunit alpha type PSMA2/PRE8 |
| P27449 | ATP6V0C | 15.736 | 5.8043 | 11.6 | 1 | 1 | 1 | 7629700 | 28872000 | plasma membrane | K02155 | C | KOG0232 | Vacuolar H+-ATPase V0 sector, subunits c/c' |
| P27816 | MAP4 | 121 | 17.429 | 6.5 | 3 | 3 | 3 | 6617700 | 45590000 | nucleus | K10431 | | | |
| P28066 | PSMA5 | 26.411 | 6.9936 | 5 | 1 | 1 | 1 | 8327000 | 11158000 | cytoplasm | K02729 | O | KOG0176 | 20S proteasome, regulatory subunit alpha type PSMA5/PUP2 |
| P28289 | TMOD1 | 40.569 | 18.487 | 15 | 4 | 4 | 3 | 12675000 | 63133000 | cytoplasm | K10370 | Z | KOG3735 | Tropomodulin and leiomodulin |
| P29692 | EEF1D | 31.121 | 19.271 | 8.5 | 1 | 1 | 1 | 8186300 | 21900000 | cytoplasm | K15410 | K | KOG1668 | Elongation factor 1 beta/delta chain |
| P29966 | MARCKS | 31.554 | 5.5636 | 5.4 | 1 | 1 | 1 | 1372200 | 6158400 | nucleus | K12561 | | | |
| P30041 | PRDX6 | 25.035 | 10.178 | 7.6 | 1 | 2 | 1 | 0 | 42061000 | mitochondria | K11188 | O | KOG0854 | Alkyl hydroperoxide reductase, thiol specific antioxidant and related enzymes |
| P30048 | PRDX3 | 27.692 | 9.6957 | 5.5 | 1 | 1 | 1 | 5081600 | 17652000 | mitochondria | K20011 | O | KOG0852 | Alkyl hydroperoxide reductase, thiol specific antioxidant and related enzymes |
| P30049 | ATP5F1D | 17.49 | 6.259 | 8.3 | 1 | 1 | 1 | 1980600 | 34064000 | mitochondria | K02134 | C | KOG1758 | Mitochondrial F1F0-ATP synthase, subunit delta/ATP16 |
| P30086 | PEBP1 | 21.057 | 11.763 | 27.8 | 2 | 2 | 2 | 1853800 | 29433000 | cytoplasm | | R | KOG3346 | Phosphatidylethanolamine binding protein |
| P30153 | PPP2R1A | 65.308 | 18.219 | 7 | 3 | 3 | 3 | 12909000 | 22634000 | cytoplasm | K03456 | T | KOG0211 | Protein phosphatase 2A regulatory subunit A and related proteins |
| P31689 | DNAJA1 | 44.868 | 6.1034 | 6.3 | 1 | 1 | 1 | 0 | 11114000 | cytoplasm | K09502 | O | KOG0712 | Molecular chaperone (DnaJ superfamily) |
| P31942 | HNRNPH3 | 36.926 | 6.3913 | 4 | 1 | 1 | 1 | 3575000 | 10546000 | cytoplasm | K12898 | A | KOG4211 | Splicing factor hnRNP-F and related RNA-binding proteins |
| P31947 | SFN | 27.774 | 12.352 | 20.6 | 3 | 3 | 2 | 9397600 | 40503000 | cytoplasm,nucleus | K06644 | O | KOG0841 | Multifunctional chaperone (14-3-3 family) |
| P32969 | RPL9 | 21.863 | 17.691 | 38.5 | 3 | 3 | 3 | 10233000 | 40700000 | cytoplasm | K02940 | J | KOG3255 | 60S ribosomal protein L9 |
| P34932 | HSPA4 | 94.33 | 14.049 | 3.9 | 2 | 2 | 2 | 7851300 | 32912000 | cytoplasm | K09489 | O | KOG0103 | Molecular chaperones HSP105/HSP110/SSE1, HSP70 superfamily |
| P35221 | CTNNA1 | 100.07 | 5.5057 | 1.4 | 1 | 1 | 1 | 4134500 | 11142000 | nucleus | K05691 | W | KOG3681 | Alpha-catenin |
| P35754 | GLRX | 11.776 | 10.667 | 39.6 | 2 | 2 | 2 | 0 | 47555000 | cytoplasm | K03676 | O | KOG1752 | Glutaredoxin and related proteins |
| P36543 | ATP6V1E1 | 26.145 | 7.1745 | 6.2 | 1 | 1 | 1 | 0 | 0 | cytoplasm | K02150 | C | KOG1664 | Vacuolar H+-ATPase V1 sector, subunit E |
| P37108 | SRP14 | 14.57 | 14.274 | 10.3 | 1 | 1 | 1 | 80715000 | 96060000 | cytoplasm | K03104 | U | KOG1761 | Signal recognition particle, subunit Srp14 |
| P40261 | NNMT | 29.574 | 6.7658 | 7.2 | 1 | 1 | 1 | 48815000 | 58823000 | cytoplasm | K00541 | | | |
| P40925 | MDH1 | 36.426 | 10.518 | 5.1 | 1 | 1 | 1 | 4149900 | 19134000 | cytoplasm | K00025 | C | KOG1496 | Malate dehydrogenase |
| P40939 | HADHA | 82.999 | 8.1466 | 2.8 | 1 | 1 | 1 | 0 | 8626500 | mitochondria | K07515 | I | KOG1683 | Hydroxyacyl-CoA dehydrogenase/enoyl-CoA hydratase |
| P41223 | BUD31 | 17 | 5.9224 | 5.6 | 1 | 1 | 1 | 0 | 15367000 | extracellular | K12873 | K | KOG3404 | G10 protein/predicted nuclear transcription regulator |
| P42167 | TMPO | 50.67 | 9.4494 | 2.6 | 1 | 1 | 1 | 10621000 | 9145500 | cytoplasm | | | | |
| P42704 | LRPPRC | 157.9 | 18.324 | 2.4 | 3 | 3 | 3 | 7661400 | 18666000 | mitochondria | K17964 | A | KOG4318 | Bicoid mRNA stability factor |
| P43307 | SSR1 | 32.235 | 9.1812 | 5.2 | 1 | 1 | 1 | 31455000 | 71071000 | endoplasmic reticulu | K13249 | U | KOG1631 | Translocon-associated complex TRAP, alpha subunit |
| P46777 | RPL5 | 34.362 | 11.266 | 10.4 | 2 | 2 | 2 | 88553 | 32137000 | nucleus | K02932 | J | KOG0875 | 60S ribosomal protein L5 |
| P47914 | RPL29 | 17.752 | 9.2572 | 9.4 | 1 | 1 | 1 | 62899000 | 83623000 | nucleus | K02905 | J | KOG3504 | 60S ribosomal protein L29 |
| P49458 | SRP9 | 10.112 | 9.0865 | 12.8 | 1 | 1 | 1 | 28588000 | 28095000 | cytoplasm | K03109 | U | KOG3465 | Signal recognition particle, subunit Srp9 |

| | | | | | | | | | | | | | | |
|---|---|---|---|---|---|---|---|---|---|---|---|---|---|---|
| P49821 | NDUFV1 | 50.817 | 12.5 | 4.7 | 2 | 3 | 2 | 5127600 | 17039000 | mitochondria | K03942 | C | KOG2658 | NADH:ubiquinone oxidoreductase, NDUFV1/51kDa subunit |
| P50238 | CRIP1 | 8.5328 | 5.7614 | 36.4 | 1 | 1 | 1 | 3685900 | 22691000 | nucleus | | TZ | KOG1700 | Regulatory protein MLP and related LIM proteins |
| P50914 | RPL14 | 23.432 | 15.847 | 10.2 | 2 | 2 | 2 | 13057000 | 44241000 | mitochondria | K02875 | J | KOG3421 | 60S ribosomal protein L14 |
| P50990 | CCT8 | 59.62 | 5.5967 | 2.2 | 1 | 1 | 1 | 7544000 | 8919100 | cytoplasm | K09500 | O | KOG0362 | Chaperonin complex component, TCP-1 theta subunit (CCT8) |
| P51148 | RAB5C | 23.482 | 5.5558 | 6.5 | 1 | 1 | 1 | 0 | 32160000 | cytoplasm | K07889 | U | KOG0092 | GTPase Rab5/YPT51 and related small G protein superfamily GTPases |
| P51571 | SSR4 | 18.998 | 8.4155 | 6.4 | 1 | 1 | 1 | 5025600 | 23569000 | plasma membrane | K04571 | U | KOG4088 | Translocon-associated complex TRAP, delta subunit |
| P51572 | BCAP31 | 27.991 | 13.625 | 7.3 | 2 | 2 | 2 | 5162900 | 26827000 | endoplasmic reticulu | K14009 | V | KOG1962 | B-cell receptor-associated protein and related proteins |
| P52292 | KPNA2 | 57.861 | 7.8933 | 2.8 | 1 | 1 | 1 | 6840100 | 11673000 | cytoplasm,nucleus | K15043 | U | KOG0166 | Karyopherin (importin) alpha |
| P52597 | HNRNPF | 45.671 | 12.146 | 15.7 | 2 | 2 | 2 | 5163300 | 61309000 | cytoplasm | K12898 | A | KOG4211 | Splicing factor hnRNP-F and related RNA-binding proteins |
| P52815 | MRPL12 | 21.348 | 11.69 | 11.6 | 2 | 2 | 2 | 0 | 0 | mitochondria | K02935 | J | KOG1715 | Mitochondrial/chloroplast ribosomal protein L12 |
| P53396 | ACLY | 120.84 | 5.6466 | 1.3 | 1 | 1 | 1 | 0 | 0 | cytoplasm | K01648 | C | KOG1254 | ATP-citrate lyase |
| P53567 | CEBPG | 16.408 | 6.3625 | 20.7 | 1 | 1 | 1 | 0 | 0 | nucleus | K10049 | | | |
| P54652 | HSPA2 | 70.02 | 7.8145 | 12.5 | 6 | 6 | 1 | 11149000 | 18344000 | cytoplasm | K03283 | O | KOG0101 | Molecular chaperones HSP70/HSC70, HSP70 superfamily |
| P54855 | UGT2B15 | 61.036 | 6.1825 | 1.3 | 1 | 1 | 1 | 67849000 | 2.55E+08 | peroxisome | K00699 | GC | KOG1192 | UDP-glucuronosyl and UDP-glucosyl transferase |
| P55084 | HADHB | 51.294 | 5.9778 | 5.5 | 1 | 1 | 1 | 0 | 67202000 | nucleus | K07509 | I | KOG1392 | Acetyl-CoA acetyltransferase |
| P60228 | EIF3E | 52.22 | 6.771 | 2.9 | 1 | 1 | 1 | 0 | 0 | cytoplasm | K03250 | J | KOG2758 | Translation initiation factor 3, subunit e (eIF-3e) |
| P60602 | ROMO1 | 8.1828 | 11.904 | 21.5 | 1 | 1 | 1 | 13480000 | 33517000 | extracellular | | S | KOG4096 | Uncharacterized conserved protein |
| P60709 | ACTB | 41.736 | 15.56 | 95.5 | 37 | 129 | 1 | 0 | 0 | cytoskeleton | K05692 | Z | KOG0676 | Actin and related proteins |
| P61026 | RAB10 | 22.541 | 8.4155 | 11 | 2 | 2 | 1 | 11795000 | 36198000 | cytoplasm | K07903 | TU | KOG0078 | GTP-binding protein SEC4, small G protein superfamily |
| P61225 | RAP2B | 20.504 | 6.0578 | 6.6 | 1 | 1 | 1 | 0 | 11994000 | cytoplasm | K07838 | R | KOG0395 | Ras-related GTPase |
| P61513 | RPL37A | 10.275 | 7.9589 | 19.6 | 1 | 1 | 1 | 12424000 | 13074000 | extracellular | K02921 | J | KOG0402 | 60S ribosomal protein L37 |
| P61586 | RHOA | 21.768 | 5.5395 | 6.2 | 1 | 1 | 1 | 2490000 | 6892600 | cytoplasm | K04513 | R | KOG0393 | Ras-related small GTPase, Rho type |
| P62263 | RPS14 | 16.273 | 12.951 | 13.9 | 2 | 2 | 2 | 8468600 | 13448000 | cytoplasm | K02955 | J | KOG0407 | 40S ribosomal protein S14 |
| P62273 | RPS29 | 6.6767 | 5.4488 | 14.3 | 1 | 1 | 1 | 9848100 | 21171000 | extracellular | K02980 | J | KOG3506 | 40S ribosomal protein S29 |
| P62304 | SNRPE | 10.803 | 14.313 | 40.2 | 2 | 2 | 2 | 1.16E+08 | 2.77E+08 | mitochondria | K11097 | A | KOG1774 | Small nuclear ribonucleoprotein E |
| P62328 | TMSB4X | 5.0526 | 5.7749 | 31.8 | 1 | 1 | 1 | 6968900 | 29006000 | nucleus | K05764 | N | KOG4794 | Thymosin beta |
| P62834 | RAP1A | 20.987 | 6.8051 | 5.4 | 1 | 1 | 1 | 2849400 | 18419000 | cytoplasm | K04353 | R | KOG0395 | Ras-related GTPase |
| P62857 | RPS28 | 7.8409 | 9.8793 | 17.4 | 1 | 2 | 1 | 34313000 | 69242000 | mitochondria | K02979 | J | KOG3502 | 40S ribosomal protein S28 |
| P62942 | FKBP1A | 11.951 | 14.963 | 28.7 | 2 | 2 | 2 | 2785200 | 32668000 | cytoplasm | K09568 | O | KOG0544 | FKBP-type peptidyl-prolyl cis-trans isomerase |
| P62995 | TRA2B | 33.665 | 6.9499 | 5.6 | 1 | 1 | 1 | 5514300 | 13524000 | nucleus | K12897 | | | |
| P63092 | GNAS | 45.664 | 10.791 | 9.4 | 3 | 3 | 2 | 2889400 | 21121000 | cytoplasm | K04632 | T | KOG0099 | G protein subunit Galphas, small G protein superfamily |
| P63173 | RPL38 | 8.2178 | 12.651 | 18.6 | 1 | 1 | 1 | 61710000 | 88103000 | nucleus | K02923 | J | KOG3499 | 60S ribosomal protein L38 |
| P63313 | TMSB10 | 5.0256 | 8.543 | 31.8 | 1 | 1 | 1 | 61494000 | 1.11E+08 | cytoplasm | K13785 | N | KOG4794 | Thymosin beta |
| P67812 | SEC11A | 20.625 | 9.0012 | 6.7 | 1 | 1 | 1 | 0 | 0 | cytoplasm | K13280 | U | KOG3342 | Signal peptidase I |
| P78417 | GSTO1 | 27.566 | 5.8036 | 3.3 | 1 | 1 | 1 | 0 | 0 | cytoplasm | K00799 | O | KOG0406 | Glutathione S-transferase |
| P84077 | ARF1 | 20.697 | 7.3522 | 21 | 2 | 2 | 1 | 0 | 17636000 | cytoplasm | K07937 | U | KOG0070 | GTP-binding ADP-ribosylation factor Arf1 |
| P84098 | RPL19 | 23.466 | 11.447 | 8.7 | 1 | 1 | 1 | 0 | 0 | cytoplasm | K02885 | J | KOG1696 | 60s ribosomal protein L19 |
| P84103 | SRSF3 | 19.329 | 6.3039 | 12.8 | 1 | 1 | 1 | 0 | 28029000 | nucleus | K12892 | A | KOG0107 | Alternative splicing factor SRp20/9G8 (RRM superfamily) |
| P98187 | CYP4F8 | 59.994 | 5.7738 | 1.5 | 1 | 1 | 1 | 0 | 0 | plasma membrane | K17728 | | | |
| Q01082 | SPTBN1 | 274.61 | 34.147 | 2.8 | 5 | 5 | 5 | 5241600 | 27824000 | cytoplasm | K06115 | Z | KOG0035 | Ca2+-binding actin-bundling protein (actinin), alpha chain |
| Q01469 | FABP5 | 15.164 | 22.266 | 25.9 | 2 | 2 | 2 | 1.89E+08 | 77514000 | extracellular | K08754 | I | KOG4015 | Fatty acid-binding protein FABP |
| Q01518 | CAP1 | 51.901 | 5.6686 | 2.7 | 1 | 2 | 1 | 0 | 0 | cytoplasm | K17261 | ZT | KOG2675 | Adenylate cyclase-associated protein (CAP/Srv2p) |
| Q01995 | TAGLN | 22.611 | 19.333 | 17.4 | 3 | 3 | 3 | 1352200 | 87694000 | cytoplasm | K20526 | Z | KOG2046 | Calponin |
| Q03701 | CEBPZ | 120.97 | 5.7113 | 2.1 | 1 | 1 | 1 | 0 | 0 | nucleus | K14832 | JK | KOG2038 | CAATT-binding transcription factor/60S ribosomal subunit biogenesis protein |
| Q05682 | CALD1 | 93.23 | 6.7682 | 2.3 | 1 | 1 | 1 | 5371700 | 40718000 | nucleus | K12327 | | | |
| Q06323 | PSME1 | 28.723 | 18.841 | 12 | 3 | 3 | 3 | 7276200 | 54218000 | nucleus | K06696 | O | KOG4470 | Proteasome activator subunit |
| Q07666 | KHDRBS1 | 48.227 | 5.9412 | 6.5 | 1 | 1 | 1 | 15118000 | 80492000 | cytoplasm | K13198 | A | KOG1588 | RNA-binding protein Sam68 and related KH domain proteins |
| Q08211 | DHX9 | 140.96 | 13.825 | 2.8 | 2 | 2 | 2 | 8517900 | 69707000 | nucleus | K13184 | A | KOG0920 | ATP-dependent RNA helicase A |
| Q12797 | ASPH | 85.862 | 8.0841 | 2.4 | 1 | 1 | 1 | 9173100 | 22070000 | plasma membrane | K00476 | O | KOG3696 | Aspartyl beta-hydroxylase |
| Q12860 | CNTN1 | 113.32 | 11.369 | 2.8 | 2 | 2 | 2 | 0 | 10733000 | plasma membrane | K06759 | T | KOG3513 | Neural cell adhesion molecule L1 |
| Q13011 | ECH1 | 35.816 | 7.2928 | 6.7 | 1 | 1 | 1 | 0 | 32763000 | mitochondria | K12663 | I | KOG1681 | Enoyl-CoA isomerase |

| | | | | | | | | | | | | | | |
|---|---|---|---|---|---|---|---|---|---|---|---|---|---|---|
| Q13423 | NNT | 113.89 | 23.343 | 5.9 | 4 | 4 | 4 | 0 | 52574000 | plasma membrane | K00323 | | | |
| Q13478 | IL18R1 | 62.303 | 6.4925 | 1.3 | 1 | 1 | 1 | 7134500 | 28674000 | peroxisome | K05173 | | | |
| Q13492 | PICALM | 70.754 | 12.388 | 8.1 | 2 | 2 | 2 | 7122700 | 63954000 | nucleus | K20044 | TU | KOG0251 | Clathrin assembly protein AP180 and related proteins, contain ENTH domain |
| Q13601 | KRR1 | 43.664 | 14.695 | 7.3 | 1 | 1 | 1 | 3981300 | 17204000 | nucleus | K06961 | JD | KOG2874 | rRNA processing protein |
| Q13617 | CUL2 | 86.982 | 5.8013 | 1.2 | 1 | 1 | 1 | 34062000 | 2.99E+08 | cytoplasm | K03870 | O | KOG2284 | E3 ubiquitin ligase, Cullin 2 component |
| Q13895 | BYSL | 49.601 | 5.7002 | 3 | 1 | 1 | 1 | 0 | 0 | cytoplasm | K14797 | W | KOG3871 | Cell adhesion complex protein bystin |
| Q14126 | DSG2 | 122.29 | 9.9498 | 2.1 | 1 | 1 | 1 | 6665900 | 26805000 | plasma membrane | K07597 | S | KOG3594 | FOG: Cadherin repeats |
| Q14573 | ITPR3 | 304.1 | 11.896 | 1.2 | 2 | 2 | 2 | 5187700 | 30066000 | plasma membrane | K04960 | T | KOG3533 | Inositol 1,4,5-trisphosphate receptor |
| Q14847 | LASP1 | 29.717 | 6.2428 | 5 | 1 | 1 | 1 | 24887000 | 51365000 | nucleus | | Z | KOG1702 | Nebulin repeat protein |
| Q14914 | PTGR1 | 35.869 | 11.436 | 7 | 1 | 1 | 1 | 10108000 | 43854000 | cytoplasm | K13948 | R | KOG1196 | Predicted NAD-dependent oxidoreductase |
| Q15005 | SPCS2 | 25.003 | 10.968 | 8.4 | 1 | 1 | 1 | 4438600 | 38679000 | plasma membrane | K12947 | U | KOG4072 | Signal peptidase complex, subunit SPC25 |
| Q15019 | SEPT2 | 41.487 | 5.4982 | 6.4 | 1 | 1 | 1 | 0 | 0 | cytoplasm | K16942 | DZU | KOG2655 | Septin family protein (P-loop GTPase) |
| Q15181 | PPA1 | 32.66 | 5.8487 | 5.5 | 1 | 1 | 1 | 4432800 | 23881000 | cytoplasm | K01507 | C | KOG1626 | Inorganic pyrophosphatase/Nucleosome remodeling factor, subunit NURF38 |
| Q15363 | TMED2 | 22.761 | 18.166 | 24.9 | 3 | 3 | 3 | 0 | 1.37E+08 | extracellular | K20347 | U | KOG1692 | Putative cargo transport protein EMP24 (p24 protein family) |
| Q15388 | TOMM20 | 16.298 | 9.0043 | 13.8 | 1 | 1 | 1 | 11250000 | 30756000 | extracellular | K17770 | U | KOG4056 | Translocase of outer mitochondrial membrane complex, subunit TOM20 |
| Q15637 | SF1 | 68.329 | 5.8963 | 3.3 | 1 | 1 | 1 | 17632000 | 15211000 | nucleus | K13095 | A | KOG0119 | Splicing factor 1/branch point binding protein (RRM superfamily) |
| Q15836 | VAMP3 | 11.309 | 8.0807 | 24 | 1 | 1 | 1 | 3027900 | 14594000 | plasma membrane | K13505 | U | KOG0860 | Synaptobrevin/VAMP-like protein |
| Q16352 | INA | 55.39 | 17.4 | 6 | 3 | 3 | 2 | 6064500 | 41792000 | nucleus | K07608 | | | |
| Q16555 | DPYSL2 | 62.293 | 8.5422 | 3.5 | 1 | 1 | 1 | 9338300 | 37294000 | cytoplasm,nucleus | K07528 | F | KOG2584 | Dihydroorotase and related enzymes |
| Q3ZCM7 | TUBB8 | 49.775 | 6.568 | 18.5 | 6 | 9 | 1 | 28679000 | 70993000 | cytoplasm | K07375 | Z | KOG1375 | Beta tubulin |
| Q53GQ0 | HSD17B12 | 34.324 | 6.6243 | 4.8 | 1 | 1 | 1 | 0 | 27252000 | cytoplasm | K10251 | I | KOG1014 | 17 beta-hydroxysteroid dehydrogenase type 3, HSD17B3 |
| Q58FF8 | HSP90AB2P | 44.348 | 9.9593 | 12.1 | 4 | 5 | 1 | 7454500 | 24015000 | cytoplasm | K04079 | | | |
| Q5GLZ8 | HERC4 | 118.56 | 5.9429 | 1.1 | 1 | 1 | 1 | 0 | 0 | nucleus | K10615 | O | KOG0941 | E3 ubiquitin protein ligase |
| Q5M775 | SPECC1 | 118.58 | 7.8801 | 1.2 | 1 | 1 | 1 | 0 | 51071000 | nucleus | | R | KOG4678 | FOG: Calponin homology domain |
| Q5RI15 | COX20 | 13.291 | 9.5323 | 19.5 | 1 | 1 | 1 | 0 | 24102000 | cytoplasm | K18184 | | | |
| Q5T2N8 | ATAD3C | 46.379 | 6.2077 | 3.4 | 1 | 1 | 1 | 3982500 | 17031000 | cytoplasm | | O | KOG0742 | AAA+-type ATPase |
| Q5TGY3 | AHDC1 | 168.35 | -2 | 0.4 | 1 | 1 | 1 | 0 | 0 | nucleus | K22592 | | | |
| Q5TGZ0 | MINOS1 | 8.8081 | 5.5134 | 41 | 1 | 1 | 1 | 0 | 0 | mitochondria | K17784 | S | KOG4604 | Uncharacterized conserved protein |
| Q5UIP0 | RIF1 | 274.46 | 5.6115 | 0.3 | 1 | 1 | 1 | 0 | 0 | cytoplasm | K11138 | | | |
| Q6PIJ6 | FBXO38 | 133.94 | 5.5397 | 0.8 | 1 | 1 | 1 | 31275000 | 36486000 | nucleus | K10313 | | | |
| Q6R327 | RICTOR | 192.22 | -2 | 0.6 | 1 | 1 | 1 | 0 | 0 | nucleus | K08267 | D | KOG3694 | Protein required for meiosis |
| Q6UW68 | TMEM205 | 21.198 | 14.074 | 6.9 | 1 | 1 | 1 | 6643700 | 31303000 | extracellular | | S | KOG2886 | Uncharacterized conserved protein |
| Q71U36 | TUBA1A | 50.135 | 12.117 | 59.9 | 18 | 28 | 1 | 0 | 0 | cytoskeleton | K07374 | Z | KOG1376 | Alpha tubulin |
| Q7Z434 | MAVS | 56.527 | 5.6769 | 4.3 | 1 | 1 | 1 | 0 | 0 | cytoplasm | K12648 | | | |
| Q7Z6B0 | CCDC91 | 49.971 | 6.3999 | 2 | 1 | 1 | 1 | 0 | 0 | cytoplasm | | | | |
| Q7Z6I8 | C5orf24 | 20.132 | 6.5904 | 11.2 | 1 | 1 | 1 | 12990000 | 20774000 | nucleus | | | | |
| Q7Z7K0 | CMC1 | 12.49 | 6.0891 | 9.4 | 1 | 1 | 1 | 3305100 | 16535000 | extracellular | K18171 | S | KOG4624 | Uncharacterized conserved protein |
| Q86U42 | PABPN1 | 32.749 | 8.1029 | 9.2 | 1 | 1 | 1 | 0 | 36300000 | nucleus | K14396 | A | KOG4209 | Splicing factor RNPS1, SR protein superfamily |
| Q86VV8 | RTTN | 248.63 | 6.4476 | 0.3 | 1 | 1 | 1 | 0 | 0 | plasma membrane | K16484 | | | |
| Q86XN8 | MEX3D | 64.882 | 6.1853 | 1.1 | 1 | 1 | 1 | 23343000 | 29115000 | cytoplasm,nucleus | K15686 | R | KOG2113 | Predicted RNA binding protein, contains KH domain |
| Q86Y39 | NDUFA11 | 14.852 | 7.763 | 10.6 | 1 | 1 | 1 | 5290400 | 25968000 | mitochondria | K03956 | | | |
| Q8IUE6 | HIST2H2AB | 13.995 | 6.7166 | 57.7 | 5 | 11 | 1 | 2.01E+08 | 1.68E+08 | nucleus | K11251 | B | KOG1756 | Histone 2A |
| Q8IVF2 | AHNAK2 | 616.62 | 13.338 | 5.2 | 2 | 2 | 2 | 1453600 | 76244000 | nucleus | | | | |
| Q8N257 | HIST3H2BB | 13.908 | 6.2488 | 50.8 | 8 | 53 | 1 | 6876600 | 5978800 | nucleus | K11252 | B | KOG1744 | Histone H2B |
| Q8N283 | ANKRD35 | 109.96 | 6.71 | 0.8 | 1 | 1 | 1 | 4123600 | 39366000 | mitochondria | | | | |
| Q8N2U0 | TMEM256 | 11.741 | 8.4805 | 24.8 | 1 | 1 | 1 | 11093000 | 56113000 | mitochondria | | S | KOG3472 | Predicted small membrane protein |
| Q8N3E9 | PLCD3 | 89.257 | 10.162 | 2.9 | 1 | 1 | 1 | 5049500 | 31490000 | mitochondria | K05857 | T | KOG0169 | Phosphoinositide-specific phospholipase C |
| Q8N4H5 | TOMM5 | 6.0352 | 6.8738 | 19.6 | 1 | 1 | 1 | 8737100 | 14948000 | cytoplasm | K17773 | | | |
| Q8N8Y2 | ATP6V0D2 | 40.426 | 11.358 | 5.1 | 1 | 1 | 1 | 3995900 | 19494000 | cytoplasm | K02146 | C | KOG2957 | Vacuolar H+-ATPase V0 sector, subunit d |
| Q8NAV1 | PRPF38A | 37.476 | 5.7403 | 3.8 | 1 | 1 | 1 | 0 | 13461000 | nucleus | K12849 | S | KOG2889 | Predicted PRP38-like splicing factor |
| Q8ND76 | CCNY | 39.336 | 5.8103 | 4.4 | 1 | 1 | 1 | 0 | 16965000 | nucleus | | R | KOG1675 | Predicted cyclin |

| | | | | | | | | | | | | | |
|---|---|---|---|---|---|---|---|---|---|---|---|---|---|
| Q8NE86 | MCU | 39.866 | 10.867 | 5.4 | 1 | 1 | 1 | 13576000 | 45060000 | mitochondria | K20858 | R | KOG2966 Uncharacterized conserved protein |
| Q8WTT2 | NOC3L | 92.547 | 13.589 | 4.4 | 2 | 2 | 2 | 9971800 | 40562000 | nucleus | K14834 | JU | KOG2153 Protein involved in the nuclear export of pre-ribosomes |
| Q8WWI1 | LMO7 | 192.69 | 9.0215 | 0.8 | 1 | 1 | 1 | 21694000 | 9338000 | nucleus | K06084 | TZR | KOG1704 FOG: LIM domain |
| Q8WXE9 | STON2 | 101.16 | 7.198 | 1 | 1 | 1 | 1 | 17216000 | 1.33E+08 | plasma membrane | K20067 | U | KOG2677 Stoned B synaptic vesicle biogenesis protein |
| Q8WZ42 | TTN | 3816 | 5.516 | 0 | 1 | 1 | 1 | 0 | 24624000 | | K12567 | Z | KOG0613 Projectin/twitchin and related proteins |
| Q92614 | MYO18A | 233.11 | 17.6 | 1.9 | 2 | 2 | 2 | 2518800 | 29474000 | nucleus | K10362 | Z | KOG0161 Myosin class II heavy chain |
| Q96A26 | FAM162A | 17.342 | 6.5455 | 13 | 1 | 1 | 1 | 0 | 15158000 | plasma membrane | | | |
| Q96AG4 | LRRC59 | 34.93 | 5.876 | 3.9 | 1 | 2 | 1 | 21430000 | 24854000 | cytoplasm,peroxisome | | S | KOG0473 Leucine-rich repeat protein |
| Q96AY2 | EME1 | 63.251 | 5.5701 | 1.6 | 1 | 1 | 1 | 0 | 2.56E+08 | nucleus | K10882 | | |
| Q96B49 | TOMM6 | 8.0019 | 7.1713 | 18.9 | 1 | 1 | 1 | 0 | 0 | cytoplasm,nucleus | K17772 | | |
| Q96CS3 | FAF2 | 52.623 | 16.557 | 10.1 | 2 | 2 | 2 | 9925200 | 53915000 | cytoplasm | K18726 | T | KOG1363 Predicted regulator of the ubiquitin pathway (contains UAS and UBX domains) |
| Q96CW1 | AP2M1 | 49.654 | 5.7793 | 3.7 | 1 | 1 | 1 | 0 | 6015300 | cytoplasm | K11826 | U | KOG0937 Adaptor complexes medium subunit family |
| Q96DI7 | SNRNP40 | 39.31 | 17.086 | 8.4 | 1 | 1 | 1 | 5436800 | 39994000 | nucleus | K12857 | A | KOG0265 U5 snRNP-specific protein-like factor and related proteins |
| Q96EU6 | RRP36 | 29.823 | 5.5604 | 5.8 | 1 | 1 | 1 | 0 | 0 | nucleus | K14795 | S | KOG3190 Uncharacterized conserved protein |
| Q96FJ2 | DYNLL2 | 10.35 | 7.7422 | 37.1 | 2 | 3 | 1 | 0 | 18194000 | cytoplasm | K10418 | Z | KOG3430 Dynein light chain type 1 |
| Q96IX5 | USMG5 | 6.4575 | 7.9504 | 25.9 | 1 | 1 | 1 | 8224600 | 38931000 | cytoplasm | K18194 | | |
| Q96LW4 | PRIMPOL | 64.411 | 5.5479 | 2.1 | 1 | 1 | 1 | 0 | 0 | cytoplasm,nucleus | K22761 | | |
| Q96S97 | MYADM | 35.273 | 11.327 | 7.1 | 1 | 1 | 1 | 23897000 | 94638000 | plasma membrane | | V | KOG4788 Members of chemokine-like factor super family and related proteins |
| Q96ST3 | SIN3A | 145.17 | 5.6198 | 1 | 1 | 1 | 1 | 0 | 0 | nucleus | K11644 | B | KOG4204 Histone deacetylase complex, SIN3 component |
| Q99439 | CNN2 | 33.697 | 7.0667 | 5.2 | 1 | 1 | 1 | 0 | 0 | cytoplasm | | Z | KOG2046 Calponin |
| Q99497 | PARK7 | 19.891 | 6.1061 | 13.8 | 1 | 1 | 1 | 0 | 0 | cytoplasm | K05687 | RV | KOG2764 Putative transcriptional regulator DJ-1 |
| Q99640 | PKMYT1 | 54.521 | 5.6913 | 1.8 | 1 | 1 | 1 | 1.94E+09 | 2.39E+09 | nucleus | K06633 | D | KOG0601 Cyclin-dependent kinase WEE1 |
| Q99878 | HIST1H2AJ | 13.936 | 10.603 | 58.6 | 8 | 37 | 0 | 0 | 0 | nucleus | K11251 | B | KOG1756 Histone 2A |
| Q9BPU6 | DPYSL5 | 61.421 | 6.1601 | 1.2 | 1 | 2 | 1 | 6.27E+08 | 7.39E+08 | cytoplasm | K07529 | F | KOG2584 Dihydroorotase and related enzymes |
| Q9BQE3 | TUBA1C | 49.895 | 11.763 | 56.3 | 17 | 24 | 1 | 40820000 | 65044000 | cytoskeleton | K07374 | Z | KOG1376 Alpha tubulin |
| Q9BRT6 | LLPH | 15.225 | 7.468 | 12.4 | 1 | 1 | 1 | 11444000 | 11424000 | nucleus | | S | KOG4811 Uncharacterized conserved protein |
| Q9BTV4 | TMEM43 | 44.875 | 19.897 | 7.5 | 3 | 3 | 3 | 2836700 | 16768000 | plasma membrane | | | |
| Q9BUF5 | TUBB6 | 49.857 | 6.74 | 16.4 | 7 | 10 | 1 | 0 | 6710500 | cytoplasm,nucleus | K07375 | Z | KOG1375 Beta tubulin |
| Q9BVA1 | TUBB2B | 49.953 | 7.6727 | 29.2 | 10 | 17 | 1 | 6663000 | 21205000 | cytoplasm | K07375 | Z | KOG1375 Beta tubulin |
| Q9BVI4 | NOC4L | 58.467 | 6.9148 | 4.1 | 1 | 1 | 1 | 16125000 | 24721000 | cytoplasm | K14771 | J | KOG2154 Predicted nucleolar protein involved in ribosome biogenesis |
| Q9BVJ6 | UTP14A | 87.977 | 10.913 | 3 | 2 | 2 | 2 | 0 | 0 | nucleus | K14567 | S | KOG2172 Uncharacterized conserved protein |
| Q9BVK6 | TMED9 | 27.277 | 19.201 | 8.9 | 2 | 2 | 2 | 4324000 | 23939000 | peroxisome | K20346 | U | KOG1690 emp24/gp25L/p24 family of membrane trafficking proteins |
| Q9BW72 | HIGD2A | 11.528 | 9.03 | 23.6 | 1 | 1 | 1 | 21147000 | 63616000 | cytoplasm,mitochondria | | R | KOG4431 Uncharacterized protein, induced by hypoxia |
| Q9BWJ5 | SF3B5 | 10.135 | 11.428 | 27.9 | 2 | 2 | 2 | 3381200 | 16432000 | extracellular | K12832 | S | KOG3485 Uncharacterized conserved protein |
| Q9BXY0 | MAK16 | 35.368 | 5.7458 | 6.3 | 1 | 1 | 1 | 4782900 | 17969000 | nucleus | K14831 | A | KOG3064 RNA-binding nuclear protein (MAK16) containing a distinct C4 Zn-finger |
| Q9BZI7 | UPF3B | 57.761 | -2 | 1.9 | 1 | 1 | 1 | 0 | 0 | cytoplasm,nucleus | K14328 | A | KOG1295 Nonsense-mediated decay protein Upf3 |
| Q9BZZ5 | API5 | 59.004 | 19.212 | 3.4 | 1 | 1 | 1 | 19785000 | 58117000 | cytoplasm | | T | KOG2213 Apoptosis inhibitor 5/fibroblast growth factor 2-interacting factor 2 |
| Q9C002 | NMES1 | 9.6172 | 13.611 | 32.5 | 2 | 2 | 2 | 0 | 0 | extracellular | | | |
| Q9C005 | DPY30 | 11.25 | 17.119 | 36.4 | 2 | 2 | 2 | 32286000 | 80674000 | cytoplasm | K14965 | K | KOG4109 Histone H3 (Lys4) methyltransferase complex, subunit CPS25/DPY-30 |
| Q9H061 | TMEM126A | 21.527 | 5.5456 | 10.3 | 1 | 1 | 1 | 3464600 | 17546000 | cytoplasm | K18157 | | |
| Q9H307 | PNN | 81.627 | 8.5561 | 2.4 | 1 | 1 | 1 | 4483000 | 26169000 | nucleus | K13114 | Z | KOG3756 Pinin (desmosome-associated protein) |
| Q9H8H0 | NOL11 | 81.123 | 7.5053 | 1.9 | 1 | 1 | 1 | 10165000 | 34734000 | cytoplasm | | | |
| Q9HAW7 | UGT1A7 | 59.818 | 5.8913 | 2.3 | 1 | 1 | 1 | 0 | 0 | plasma membrane | K00699 | GC | KOG1192 UDP-glucuronosyl and UDP-glucosyl transferase |
| Q9HBG6 | IFT122 | 141.82 | 10.434 | 1.5 | 2 | 2 | 2 | 26461000 | 2507200 | cytoplasm | K19656 | R | KOG1538 Uncharacterized conserved protein WDR10, contains WD40 repeats |
| Q9HDC9 | APMAP | 46.48 | 6.7424 | 6.5 | 1 | 1 | 1 | 11600000 | 44347000 | plasma membrane | K21407 | R | KOG1520 Predicted alkaloid synthase/Surface mucin Hemomucin |
| Q9NPE3 | NOP10 | 7.7059 | 12.458 | 20.3 | 1 | 1 | 1 | 44365000 | 1.7E+08 | cytoplasm | K11130 | A | KOG3503 H/ACA snoRNP complex, subunit NOP10 |
| Q9NPG4 | PCDH12 | 128.99 | 5.6904 | 0.8 | 1 | 1 | 1 | 37543000 | 0 | endoplasmic reticulu | K16499 | S | KOG3594 FOG: Cadherin repeats |
| Q9NQC3 | RTN4 | 129.93 | 11.755 | 4.9 | 2 | 2 | 2 | 2.46E+08 | 2.87E+08 | plasma membrane | K20720 | U | KOG1792 Reticulon |
| Q9NQG5 | RPRD1B | 36.899 | 12.844 | 7.4 | 2 | 2 | 2 | 4327400 | 14111000 | nucleus | K15559 | A | KOG2669 Regulator of nuclear mRNA |
| Q9NSI2 | FAM207A | 25.456 | 5.9571 | 6.1 | 1 | 1 | 1 | 0 | 0 | nucleus | | | |
| Q9NX40 | OCIAD1 | 27.626 | 12.761 | 18.8 | 2 | 2 | 2 | 7684700 | 47680000 | nucleus | | | |

| | | | | | | | | | | | | | | |
|---|---|---|---|---|---|---|---|---|---|---|---|---|---|---|
| Q9NY93 | DDX56 | 61.589 | 9.3496 | 2.7 | 1 | 1 | 1 | 7122800 | 11793000 | nucleus | K14810 | A | KOG0346 | RNA helicase |
| Q9NZ45 | CISD1 | 12.199 | 5.9327 | 12 | 1 | 1 | 1 | 0 | 14742000 | extracellular | | R | KOG3461 | CDGSH-type Zn-finger containing protein |
| Q9NZN4 | EHD2 | 61.161 | 20.895 | 10.3 | 3 | 3 | 3 | 5540300 | 80564000 | cytoplasm | K12469 | TU | KOG1954 | Endocytosis/signaling protein EHD1 |
| Q9P035 | HACD3 | 43.159 | 6.1709 | 6.1 | 1 | 1 | 1 | 3325100 | 47841000 | cytoplasm | K10703 | R | KOG3187 | Protein tyrosine phosphatase-like protein PTPLA (contains Pro) |
| Q9P0J0 | NDUFA13 | 16.698 | 18.473 | 16 | 2 | 2 | 2 | 7295600 | 37105000 | cytoplasm | K11353 | CD | KOG3300 | NADH:ubiquinone oxidoreductase, B16.6 subunit/cell death-regulatory protein |
| Q9P2E9 | RRBP1 | 152.45 | 5.6389 | 1.2 | 1 | 1 | 1 | 0 | 0 | endoplasmic reticulu | K14000 | | | |
| Q9UBD6 | RHCG | 53.178 | 23.575 | 9.2 | 2 | 3 | 2 | 3858700 | 70541000 | plasma membrane | K06580 | UR | KOG3796 | Ammonium transporter RHBG |
| Q9UDW1 | UQCR10 | 7.3084 | 7.2677 | 27 | 1 | 1 | 1 | 4543400 | 12839000 | mitochondria | K00419 | C | KOG3494 | Ubiquinol cytochrome c oxidoreductase, subunit QCR9 |
| Q9UJC5 | SH3BGRL2 | 12.326 | 5.5437 | 12.1 | 1 | 1 | 1 | 0 | 21047000 | cytoplasm | | S | KOG4023 | Uncharacterized conserved protein |
| Q9UM00 | TMCO1 | 21.175 | 8.3169 | 8 | 1 | 1 | 1 | 7150400 | 45256000 | extracellular | K21891 | S | KOG3312 | Predicted membrane protein |
| Q9UMY1 | NOL7 | 29.426 | 6.7256 | 5.4 | 1 | 1 | 1 | 15385000 | 21291000 | nucleus | | | | |
| Q9UQ80 | PA2G4 | 43.786 | 6.1233 | 5.8 | 1 | 1 | 1 | 11917000 | 13548000 | nucleus | | R | KOG2776 | Metallopeptidase |
| Q9Y224 | RTRAF | 28.068 | 14.607 | 13.9 | 2 | 2 | 2 | 18730000 | 57139000 | cytoplasm | K15433 | R | KOG4380 | Carnitine deficiency associated protein |
| Q9Y241 | HIGD1A | 10.143 | 7.1218 | 19.4 | 1 | 1 | 1 | 0 | 15337000 | cytoplasm | | R | KOG4431 | Uncharacterized protein, induced by hypoxia |
| Q9Y281 | CFL2 | 18.736 | 10.625 | 39.2 | 5 | 8 | 2 | 7478900 | 33080000 | mitochondria | K05765 | Z | KOG1735 | Actin depolymerizing factor |
| Q9Y295 | DRG1 | 40.542 | 5.5475 | 3 | 1 | 1 | 1 | 0 | 0 | cytoplasm | | T | KOG1487 | GTP-binding protein DRG1 (ODN superfamily) |
| Q9Y333 | LSM2 | 10.834 | 13.51 | 28.4 | 2 | 3 | 2 | 14493000 | 76749000 | cytoplasm | K12621 | A | KOG3448 | Predicted snRNP core protein |
| Q9Y3B3 | TMED7 | 25.171 | 7.7208 | 5.8 | 1 | 1 | 1 | 0 | 18200000 | mitochondria | K20349 | U | KOG1693 | emp24/gp25L/p24 family of membrane trafficking proteins |
| Q9Y3B4 | SF3B6 | 14.585 | 5.5693 | 11.2 | 1 | 1 | 1 | 14344000 | 36025000 | nucleus | K12833 | R | KOG0114 | Predicted RNA-binding protein (RRM superfamily) |
| Q9Y3B9 | RRP15 | 31.484 | 5.5586 | 3.9 | 1 | 1 | 1 | 0 | 0 | nucleus | | S | KOG2974 | Uncharacterized conserved protein |
| Q9Y3C1 | NOP16 | 21.188 | 6.2276 | 3.9 | 1 | 1 | 1 | 8380600 | 12981000 | nucleus | | S | KOG4706 | Uncharacterized conserved protein |
| Q9Y3T9 | NOC2L | 84.918 | 8.1003 | 2.1 | 1 | 1 | 1 | 2564200 | 15010000 | nucleus | K14833 | J | KOG2256 | Predicted protein involved in nuclear export of pre-ribosomes |
| Q9Y5J1 | UTP18 | 62.003 | 6.0631 | 3.1 | 1 | 1 | 1 | 0 | 11635000 | nucleus | K14553 | R | KOG2055 | WD40 repeat protein |

**Supplementary Table 3. A representative list of proteins subject to remarkable methylation or demethylation in senescent cells. Related to Fig. 1.**

**A. Differentially modified sites summary (Filtered with threshold value of expression fold change, dimethylation)**

| Compare group | Regulated type | Fold change >1.2 | Fold change >1.3 | Fold change >1.5 | Fold change >2 |
|---|---|---|---|---|---|
| SEN/CTRL | up-regulated | (0) | (0) | (0) | (0) |
| | down-regulated | 1 (1) | 1 (1) | 1 (1) | (0) |

| Protein accession | Position | Amino acid | Protein description | Gene name | Localization probability | PEP | Score | Modified sequence | Charge | Mass error [ppm] | MS/MS Count | CTRL | SEN | SEN/CTRL Ratio |
|---|---|---|---|---|---|---|---|---|---|---|---|---|---|---|
| Q71DI3 | **36** | **K** | **Histone H3.2** | HIST2H3A | 0.837398 | 0.0267953 | 64.22 | KSAPATGGVK(0.837)K(0.16 | 4 | -2.4441 | 1 | 5175500 | 15592000 | **0.554** |

**B. Differentially modified sites summary (Filtered with threshold value of expression fold change, trimethylation)**

| Compare group | Regulated type | Fold change >1.2 | Fold change >1.3 | Fold change >1.5 | Fold change >2 |
|---|---|---|---|---|---|
| BLEO/CTRL | up-regulated | 1 (1) | 1 (1) | 1 (1) | (0) |
| | down-regulated | 3 (2) | 3 (2) | 2 (2) | (0) |

| Protein accession | Position | Amino acid | Protein description | Gene name | Localization probability | PEP | Score | Modified sequence | Charge | Mass error [ppm] | MS/MS Count | CTRL | SEN | SEN/CTRL Ratio |
|---|---|---|---|---|---|---|---|---|---|---|---|---|---|---|
| P05141 | 52 | K | ADP/ATP translocase 2 | SLC25A5 | 1 | 0.001817 | 129 | QYK(1)GIIDCVVR | 2 | -0.78278 | 2 | 42366000 | 1.75E+08 | 1.66 |
| P0DP25 | 116 | K | Calmodulin-3 | CALM3 | 1 | 4.783E-18 | 231.9 | HVMTNLGEK(1)LTDEEVDEI | 3 | -0.4112 | 11 | 1.54E+09 | 7.47E+09 | 0.863 |
| P68104 | 318 | K | Elongation factor 1-alpha 1 | EEF1A1 | 1 | 0.0643065 | 63.82 | NVSVK(1)DVR | 2 | -0.84943 | 1 | 36018000 | 46913000 | 0.526 |
| P68104 | 79 | K | Elongation factor 1-alpha 1 | EEF1A1 | 1 | 8.348E-07 | 141 | GITIDISLWK(1)FETSK | 3 | -0.05163 | 2 | 46087000 | 54182000 | 0.701 |
| Q71DI3 | **27** | **K** | **Histone H3.2** | HIST2H3A | 0.999996 | 0.0267953 | 64.22 | K(1)SAPATGGVKKPHR | 4 | -2.4441 | 1 | 5175500 | 15592000 | **0.554** |
| Q9HBG6 | 711 | K | Intraflagellar transport protein 122 homolo | IFT122 | 1 | 0.0500242 | 78.52 | FHEAAK(1)LYK | 2 | 0.015077 | 1 | 0 | 1.32E+08 | |

**Supplementary Table 4. Primer sequences for qRT-PCR assays. Related to Fig. 3, 4, 5, and 7.**

| Gene name | Forward (5'-3') | Reverse (5'-3') |
|---|---|---|
| ***IL6*** | TTCTGCGCAGCTTTAAGGAG | AGGTGCCCATGCTACATTTG |
| ***CXCL8*** | ATGACTTCCAAGCTGGCCGTG | TGTGTTGGCGCAGTGTGGTC |
| ***IL-1α*** | AATGACGCCCTCAATCAAAG | TGGGTATCTCAGGCATCTCC |
| ***IL-1β*** | TGGGTATCTCAGGCATCTCC | TTCTGCTTGAGAGGTGCTGA |
| ***GM-CSF*** | ATGTGAATGCCATCCAGGAG | AGGGCAGTGCTGCTTGTAGT |
| ***AREG*** | AGCTGCCTTTATGTCTGCTG | TTTCGTTCCTCAGCTTCTCC |
| ***CXCL1*** | CACCCCAAGAACATCCAAAG | TAACTATGGGGGATGCAGGA |
| ***CXCL3*** | GGAGCACCAACTGACAGGAG | CCTTTCCAGCTGTCCCTAGA |
| ***SPINK1*** | CCTTGGCCCTGTTGAGTCTA | GCCCAGATTTTTGAATGAGG |
| ***WNT16B*** | GCTCCTGTGCTGTGAAAACA | TGCATTCTCTGCCTTGTGTC |
| ***MMP3*** | AGGGAACTTGAGCGTGAATC | TCACTTGTCTGTTGCACACG |
| $p16^{INK4a}$ | CTTCCTGGACACGCTGGT | ATCTATGCGGGCATGGTTAC |
| $P21^{CIP1}$ | ATGAAATTCACCCCCTTTCC | CCCTAGGCTGTGCTCACTTC |
| ***KDM4A*** | TGGACTTGGTGGAAAAGGAG | GTCTTCAGTGTGCCAAGCAA |
| ***KDM4B*** | GGACTGACGGCAACCTCTAC | CGTCCTCAAACTCCACCTG |
| ***KDM4C*** | TGCCTGTCGTGTTTTCTCAG | CATGTCGAGCAACTTCAGGA |
| ***KDM4D*** | TTTCCCTATGGCTACCATGC | TCATAGCGTTCAGGTTGCAG |
| ***SUV39H1*** | GTCATGGAGTACGTGGGAGAG | CCTGACGGTCGTAGATCTGG |
| ***RPL13A*** | GTACGCTGTGAAGGCATCAA | CGCTTTTTCTTGTCGTAGGG |
